# H2A.Z facilitates Sox2-nucleosome interaction by promoting DNA and histone H3 tail mobility

**DOI:** 10.1101/2025.03.06.641691

**Authors:** Helen K. Moos, Rutika Patel, Sophie K. Flaherty, Sharon M. Loverde, Evgenia N. Nikolova

**Author notes:** Correspondence to Evgenia N. Nikolova.

## Abstract

Epigenetic regulation of chromatin structure is strongly influenced by histone variants and post-translational modifications. The conserved histone variant H2A.Z has been functionally linked to the pioneer factors Sox2 and Oct4, which open chromatin and activate cell fate-specific transcriptional programs. However, the molecular basis of this interaction is not well understood. Here, we combine biochemistry, NMR spectroscopy, and molecular dynamics (MD) simulations to investigate how H2A.Z nucleosome dynamics influence pioneer factor binding. We find that H2A.Z enhances Sox2 and Oct4 association at distinct positions within 601 nucleosomes, correlating with increased DNA accessibility and altered H3 N-terminal tail dynamics. We show that the H3 tail competes with Sox2 for DNA binding and is more efficiently displaced with H2A.Z, while also allowing for unique Sox2-H3 tail interactions. MD simulations reveal that H2A.Z promotes DNA unwrapping, increases inter-gyre spacing, and enhances H3 tail flexibility, while simultaneously reducing contacts with DNA and with the H2A.Z C-terminal tail. This destabilizing effect is DNA-sequence dependent and prominent in the less stable Lin28B nucleosome, which Sox2 appears to substantially reshape. Together, our results suggest that H2A.Z promotes pioneer factor binding by increasing DNA accessibility and reducing histone tail competition, with broad implications for epigenetic regulation and chromatin recognition.

## Introduction

Epigenetic changes such as post-translational modifications (PTMs) and histone variants play critical roles in regulating the structure and function of chromatin.^1, 2^ H2A.Z is a highly conserved histone variant that is crucial for transcriptional regulation, genome integrity, DNA damage repair, and chromosome segregation.^3-6^ H2A.Z is required for embryonic stem cell renewal and differentiation,^7^ and its dysregulation has been linked to embryonic abnormalities and cancer.^6,8^ Despite extensive studies, how H2A.Z regulates chromatin structure remains to be characterized on a molecular level.

Within the genome, H2A.Z is enriched at regulatory regions of both active and inactive genes^9^ and, depending on its location, can either activate or repress transcription.^10-12^ At actively transcribing genes, H2A.Z is frequently acetylated^13^ and its occupancy strongly correlates with epigenetic marks such as H3K4 methylation.^14,15^ Functionally, H2A.Z alters chromatin architecture by reducing nucleosome occupancy and increasing chromatin accessibility *in vivo*.^16-18^ This variant has been closely linked to pioneer transcription factors^7,19-22^ that promote chromatin opening and remodeling during embryonic stem cell (ESC) renewal and differentiation.^23,24^ For example, nucleosome-occupied regions of the pioneer factors Sox2 and Oct4 are marked by H2A.Z and its depletion compromises Oct4 binding to target genes.^7, 21, 25, 26^ H2A.Z can also cooperate with Sox2 in recruiting the repressive complex PRC2, ^25^ underscoring its versatile role in mediating chromatin structure and its interaction with regulatory proteins that orchestrate cell fate.

Mechanistically, these effects may relate to the impact of H2A.Z on nucleosome dynamics. Increases in nucleosome “breathing” or DNA unwrapping enhance transcription factor binding *in vitro*,^27-30^ further supported by computational studies.^31-34^ Nuclear magnetic resonance (NMR) spectroscopy and molecular dynamics (MD) simulations show that DNA unwrapping is coupled to the dynamics of the flexible histone tails protruding from the nucleosome core, whose transient DNA interactions can hinder regulatory protein binding.^35-42^ Interactions of the H3 N-terminal and H2A C-terminal tail at super-helical locations (SHLs) near entry/exit DNA are especially important because these regions serve as primary docking sites for transcription factors.^38,43-45^Post-translational modifications (PTMs), proteolytic cleavage, and oncogenic mutations that frequently affect H3 and H2A tails^46-48^ further tune these interactions and modulate chromatin accessibility.^1,34,36,42,49^ Together, these findings suggest that pioneer factors exploit variations in nucleosome dynamics to efficiently recognize their targets, promoting chromatin opening and the recruitment of additional regulatory complexes.

Building on this model, structural and biochemical evidence directly supports the role of H2A.Z in promoting chromatin destabilization. Incorporating H2A.Z in nucleosomes leads to lower thermal stability and increased DNA unwrapping compared to canonical systems.^50, 51^ The unique H2A.Z N- and C-terminal regions, which carry fewer positively charged residues, have been implicated in this weakening of histone-DNA contacts.^52,53^Consistent with this, cryo-electron microscopy (cryo-EM) structures of H2A.Z nucleosomes assembled on the Widom 601 positioning sequence reveal enhanced asymmetric DNA unwrapping and weaker interactions between the H2A.Z C-terminal region, DNA, and histone H3.^51^ However, the flexible tails of histones, including H2A.Z, are not resolved by cryo-EM, preventing insight into their conformation, dynamics, and interactions. MD simulations help bridge this gap, showing that H2A.Z incorporation increases histone plasticity and couples DNA unwrapping to loss of H2A.Z C-terminal tail-DNA contacts.^53, 54^ Despite these advances, how additional histone tail perturbations shape H2A.Z nucleosome dynamics and pioneer factor engagement remains poorly understood.

Because H2A.Z frequently co-occupies regulatory regions with pioneer factors, understanding its effect on nucleosome dynamics is crucial for deciphering how these factors engage closed chromatin. Indeed, H2A.Z-associated Sox2 and Oct4 are key pioneer factors that engage silent chromatin in ESCs and initiate transcriptional programs that govern cell-fate decisions.^24, 55^ They are also essential for somatic cell reprogramming into induced pluripotent stem cells (iPSC).^56, 57^ Sox2 and Oct4 can bind synergistically, interacting on adjacent or closely spaced motifs *in vitro*^58,59^and co-localizing on target genes *in vivo*.^56, 60^ Sox2 contains a partially disordered high-mobility group (HMG) box DNA-binding domain that folds upon engaging the DNA minor groove, inducing strong bending and local unwinding of the helix.^61, 62^ The extent of Sox2-nucleosome binding and DNA deformation is determined by multiple factors, including the intrinsic flexibility of Sox2, as well as the DNA sequence, accessibility, and local mobility.^63-66^Biochemical and structural studies,^64-66^ backed by MD simulations and molecular modeling,^67, 68^ have shown that Sox2 preferentially targets solvent-exposed sites on nucleosomes, with strongest enrichment at SHL5 and lower preference for SHL2 and SHL6 on Widom 601 nucleosomes. Oct4, in contrast, recognizes DNA using a bipartite DBD composed of a POU-specific (POU_S_) domain and a POU-homeodomain (POU_HD_) joined by a flexible linker, allowing the two domains to contact opposite faces of the DNA major groove.^61, 62^ Oct4 preferentially engages entry/exit sites, where at least one half-motif is accessible, or linker DNA, where its binding is modulated by interactions with the H3 and H4 N-terminal tails. ^65, 69-71^ In inherently less stable nucleosomes that exhibit greater DNA mobility, such as Lin28B, Oct4 can gain access to more internal sites.^69, 70, 72^ These observations underscore how DNA dynamics dictate pioneer factor interactions with chromatin, emphasizing the need to understand how H2A.Z incorporation and other epigenetic modifications tune this process.

Here, we dissect the mechanisms by which H2A.Z modulates Sox2 and Oct4 engagement with nucleosomes. Using an integrated approach combining NMR spectroscopy, MD simulations, biochemical assays, and targeted mutagenesis, we probe the interplay between DNA mobility, histone tail dynamics, and pioneer factor binding. Our results reveal that H2A.Z enhances Sox2 and Oct4 association with both end-positioned sites (SHL6) and internal sites (SHL2 for Sox2), where DNA unwrapping is increased and H3 tail dynamics and DNA interactions are altered. Notably, Sox2 more effectively displaces the H3 N-terminal tail at these sites in the presence of H2A.Z. Complementary MD simulations show that H2A.Z nucleosomes exhibit greater DNA unwrapping and inter-gyre gapping, which is coupled to more dynamic H3 tail conformations and reduced contacts between the H3 tail, DNA, and the H2A.Z C-terminal tail. Together, these findings suggest that H2A.Z facilitates Sox2 and Oct4 engagement primarily by promoting DNA mobility and loosening H3 tail-DNA interactions, thereby rendering nucleosomal sites more accessible.

## Materials and Methods

### DNA preparation

DNA for mouse Fgf4 (CTTTGTTTGGATGCTAAT) and Nanog (CATTGTAATGCAAAA) Sox2-Oct4 composite motifs were inserted via standard mutagenesis within the Widom 601 sequence in pGEM-3z/601 plasmid (gift from Prof. Gregory Bowman, JHU), which we modified on one end to remove a strong non-cognate Sox2 binding site (Table S1). The Lin28B plasmid in pBluescript II SK(+) was ordered from Gene Universal. Large scale PCR reactions (10-60 ml) were prepared as described^66^ using unlabeled DNA primers or 5′-end labeled fluorescent primers (6-FAM or Cy3) obtained from IDT with HPLC purification. PCR product was purified by vertical gel electrophoresis on a 6% (60:1 Acrylamide/Bisacrylamide) native gel column using a 491 Prep Cell (Bio-Rad) run at 11 W and 4 °C, as described.^73^ Fractions were analyzed via agarose gel, concentrated, and stored at -20 °C.

### Sox2 and Oct4 preparation

The human Sox2 DBD (39-118), and Oct4 DBD (133-296) (Table S2) were designed as described previously.^66^ An Oct4 “CtoS” mutant was created via standard mutagenesis where all cysteine residues were substituted with serine to create a stabilized construct and reduce protein aggregation. DNA binding was tested and found to be equivalent to the wild-type protein. Proteins were expressed in BL21(DE3) cells in TB media. Cells were grown with shaking at 37 °C until OD_600_ ~0.6, induced with 0.5 mM IPTG, then grown for 4 h at 37 °C for Sox2, and 16-18 h at 18 °C for Oct4. Cells were harvested by centrifugation at 3600 RPM for 20 min, and pellets were stored at -20 °C. Purification was done via a two-step Ni^2^+ affinity chromatography on an AKTA FPLC system (Cytiva), as described previously,^66^ where the His-tag was cleaved using an in-house His-tagged 3C protease.

### Histone preparation

Human H2A.Z in pET21a was gifted by Prof. Carl Wu (JHU). *Xenopus laevis* (XL) histones H2A, H2B, H3, and H4 in pET3a were a gift from Prof. Gregory Bowman (JHU). Mutant constructs were prepared using standard mutagenesis. All histone constructs are shown in Table S2. XL histone plasmids and H2A^C-H2A.Z^ mutant were transformed into BL21(DE3) pLysS for expression in 2XTY media (unlabeled) or M9 media (isotopic labelling), except for ^15^N-labeled H4 that was expressed in BL21(DE3) cells. Human H2A.Z and mutant constructs were expressed in BL21(DE3) cells. For ^15^N or ^15^N,^13^C labeling, M9 media was supplemented with 1 g ^15^NH_4_Cl and 2 g ^13^C-glucose (CIL). For XL H2A, H2B, H3, and human H2A.Z, cell pellets were purified on a 20 ml Q-SP HiTrap HP column (Cytiva, 2×5 ml Q on top of 2×5 ml SP), using a Sodium Acetate-Urea buffer, as previously described.^66^ Histone H4 was prepared from inclusion bodies, followed by a desalting column (HiPrep 26/10 (GE)) and a 2X Q-SP column (Cytiva), as previously described.^66, 74^

### Nucleosome preparation

Lyophilized histones were resuspended in 20 mM Tris-HCl pH 7.8, 6 M Guanidine-HCl, 5 mM DTT at roughly 2 mg/mL with approximately 15-20% molar excess of H2A/H2B and refolded by dialyzing 4 times into Refolding Buffer (10 mM Tris-HCl pH 7.8, 2 M NaCl, 1 mM EDTA, 5 mM BME) using a 3.5kDa membrane (Spectra Labs) at 4°C. The refolded octamer (and excess H2A-H2B dimer) was then concentrated and purified by using a HiLoad 16/600 Superdex 200 pg column (Cytiva), as described.^73, 74^ Fractions were analyzed by SDS-page gel, concentrated to 50-150 μM, flash frozen, and stored at - 80°C. Nucleosomes were reconstituted following an adapted salt-dialysis protocol,^73, 74^ where purified octamer (6 μM), dimer (1.8 μM), and 10% excess of DNA (6.6 μM) were combined. Nucleosomes were concentrated and purified via vertical gel electrophoresis as for DNA, except using a 7% (60:1 Acrylamide/Bisacrylamide) native gel column.^66^ Nucleosome purity was assessed by 5% native gel electrophoresis: 601 nucleosomes exhibited high purity, while Lin28B nucleosomes showed a minor second population (~ 17%) and small free DNA contamination. Incubation of Lin28B NCPs at 37°C for 3 hours to remove kinetically trapped states (i.e. heat shifting) did not resolve the minor population (see Fig. S13E). This state was unlikely due to alternate DNA positioning, which was previously determined to occur with shorter reconstitution protocols and the use of XL histones.^75^

### Electrophoretic mobility shift assay (EMSA)

10 nM of 5′-FAM/5′-Cy3-labeled nucleosome (NCP) was mixed with variable concentrations of Sox2 HMG or Oct4 (CtoS mutant) protein (10 nM to 300 nM) in binding buffer (10 mM Tris-HCl pH 7.5, 150 mM KCl, 1 mM EDTA pH 8.0, 1 mM DTT, 0.5 mg/ml Recombinant Albumin (NEB), 5% glycerol) in a total reaction volume of 20 μL and incubated at 25 °C for 45 min. Reactions were loaded (2 μL) on a 5% native polyacrylamide gel and run for 120 min at 100 V on ice. Binding was visualized using a Typhoon 5 imager (GE Amersham) using FAM fluorescence. The integrated peak intensities for free and bound species were quantified using GelAnalyzer 19.1 (available at www.gelanalyzer.com, by Istvan Lazar Jr., PhD and Istvan Lazar Sr., PhD, CSc). For 601* NCPs, the fraction bound for the specific complex at a given gel lane was calculated as I_SC_/(I_SC_+I_NC_+I_F_), where I_SC_ is the signal for the specific complex (discrete shifted band), I_NC_ is the signal for any non-specific complex (smear or supershifted band), I_F_ is the signal for the free nucleosome in that gel lane. For Lin28B NCPs, where multiple Sox2/Oct4 binding sites are present, the “specific” fraction bound was calculated as above using the signal for the first discrete binding event, while the “total” fraction bound was calculated as (I_SC_+I_MC_)/(I_SC_+I_MC_+I_F_), where I_MC_ was the supershifted signal reflecting multi-site or non-specific binding. Fraction bound was plotted as the mean ± standard deviation (s.d.) for 3-4 independent experiments.

### DnaseI footprinting assay

Reactions were set up in 10 mM Tris-HCl pH 7.5, 100 mM KCl, 1 mM DTT, 0.2 mg/ml Recombinant Albumin (NEB), 0.02% Tween-20, 8% Glycerol, containing 100 nM Lin28B DNA or NCP. Binding reactions were set up with or without Sox2 HMG (0.1 μM or 1 μM), and Oct4 CtoS (0.3 μM or 3 μM), and incubated at 25 °C for 45 min. DNaseI (NEB) was added at 0.5 U (or 0.125 U) for NCP and 0.1 U (or 0.025 U) and incubated for 5 min at 25°C, inactivated by 40 μL quench buffer (10 mM Tris-HCl pH 7.5, 50 mM EDTA, 2% SDS, 300 ng/ul Glycogen (Invitrogen)) and heating at 75 °C for 30 min, as previously described.^66^ DNA was purified using Phenol-Chloroform extraction and ethanol precipitated. Samples were run on a denaturing sequencing gel.^66^ Gels were scanned on a Typhoon 5 imager (GE Amersham) as described for EMSA. For DNaseI assay, 2 independent measurements were performed and at 2 different DnaseI concentrations, as described above. Gel lane analysis was performed with ImageJ2^76^ using background subtraction and contrast adjustment. Minor groove analysis was performed using the X3DNA server.^59^ The change in DNA minor groove width for bound versus unbound NCP was obtained using the following structures: PDB ID: 6T79 (unbound) and 6T7B (Sox2-bound) for SHL+2;^64^ PDB ID: 6T93 (unbound) and 6T90 (Sox2-Oct4-bound) for SHL-6; PDB ID: 6T93 (unbound) and 6YOV (Sox2-Oct4-bound) for SHL+5.^65^

### Exonuclease III assay

Reactions were set up in 10 μL of 10 mM Tris-HCl pH 7.5, 150 mM KCl, 1 mM DTT, 1 mM EDTA, 0.2 mg/ml BSA, 0.02% Tween-20, 8% Glycerol containing 100 nM NCP, with or without Sox2 HMG (0.3 to 1 μM), and/or Oct4 (1 to 3 μM), and incubated at 25 °C for 45 min. ExonucleaseIII (NEB) dilutions were created with equal parts stock enzyme (100 U/μL) and reaction buffer supplemented with 76.67 mM MgCl_2_. 1.5 μL of enzyme dilution or reaction buffer was added to each reaction (10 mM MgCl_2_ final) and incubated for the desired time (0 to 60 min). Reaction was stopped with 40 μL quench buffer (10 mM Tris pH 7.5, 20 mM EDTA, 1.2% SDS, 100 ng/μl glycogen) and inactivated at 75 °C for 20 min. DNA was purified using Phenol-Chloroform extraction and ethanol precipitated. Samples were prepared and run on a denaturing sequencing gel and analyzed with ImageJ2^76^ to obtain lane plots as described for DNaseI footprinting. The fraction of DNA at each SHL was quantified using GelAnalyzer 19.1 (mean ± s.d. of 3 independent experiments (2 for H2A/H2A.Z swap mutants)) by dividing the integrated intensity of DNA at each SHL by the combined signal of undigested and digested DNA in each lane. To extract ExoIII decay rates, the DNA fraction at SHL±7 was fit to a mono-exponential decay using a python script and a 5% signal error, and the mean ± s.d. was reported in Table S3.

### Restriction enzyme digest assay

Reactions were set up in 20 μL rCutsmart buffer (NEB) using 10 nM NCP on ice and 10 U of MslI (NEB). Then reactions were incubated for 2 hrs at 25 °C or 37 °C, quenched with 60 μL Quench buffer (50 mM Tris Acetate pH 7.5, 50 mM EDTA pH 8, 2% SDS, 0.2 mg Glycogen), and heated at 85ºC for 20 min. Samples were prepared using Phenol-Chloroform extraction, ethanol precipitated, and resuspended in 10 μL Orange G Formamide dye as described in DnaseI assay. 5 μL of reaction was loaded onto an 8 M Urea/8% Acrylamide gel and run for 2 hrs at 130 V. Gels were imaged as described in EMSA.

### Förster resonance energy transfer (FRET) assay

Reactions were set up in 120 μL EMSA binding buffer containing 20 nM NCP or DNA with 5′-(6-FAM)/5′-Cy3 labeling. For binding, proteins (Sox2 HMG, Oct4 CtoS) were diluted into buffer and added at a 200 nM final concentration (10X). Fluorescence emission scans were measured on an ATF 105 Differential/Ratio Spectrofluorometer (AVIV). For FRET, FAM was excited at 495 nm and data was collected for 505-600 nm range. To correct for contributions from donor bleed-through at the acceptor peak (560 nm) and direct acceptor excitation when exciting at 495 nm, a correction factor α was calculated as follows: α = I_560,DNA_/I_520,DNA_, where I_560,DNA_ and I_520,DNA_ are the intensity at 560 nm and 520 nm, respectively, of doubly labeled DNA, where no significant FRET is expected to occur (DNA contour length L = 145 bp(0.34 nm/bp) = 49.3 nm). For this calculation (α = 0.412 ± 0.015), 3 replicates of 5 different 601* DNAs and Lin28B DNA, which showed nearly identical α values on their own, were averaged. The corrected acceptor intensity (I_A,corr_) and FRET ratio were calculated as follows: I_A,corr_ = I_A_ – αI_D_; FRET = I_A,corr_ /(I_D_ + I_A,corr_), where I_A_ and I_D_ are the measured acceptor and donor intensity. Reported FRET values represent the mean and s.d. of 2-3 technical replicates (same stock, same buffer preparation) and 2-6 independent measurements (different stock dilutions, different buffer preparation). FRET values and associated statistics are reported in Table S4; the combined s.d. was calculated as s.d._total_ = ((s.d._sample_)^2^/n + (s.d._mean_)^2^)^1/2^, where 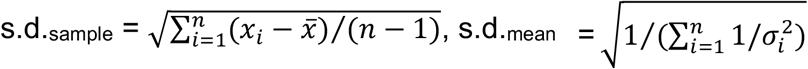 is the propagated error in α), and n is the total number of measurements. Statistical significance between datasets (NCPs) was assessed using a two-tailed Welch’s t-test, with p-values reported in Table S4. We verified that spectra for FAM-only labeled NCP and DNA were identical and showed the same contribution at the acceptor wavelength (560 nm). A Cy3-only labeled DNA, when excited at 495 nm, showed no contribution to I_D_ (i.e. acceptor bleed-through was ~0 at 520 nm) but measurable direct acceptor excitation (560 nm), which was accounted for by the correction factor α.

### NMR spectroscopy

Nucleosome samples were prepared with a single isotopically labeled histone (^15^N-labeled H3 or H4; ^15^N,^13^C-labeled H2A or H2A.Z). Samples were dialyzed 3 times against NMR buffer (20 mM Tris-HCl pH 7.0, 25 mM NaCl, 0.5 mM TCEP, 1 mM EDTA) using a 3.5 kDa membrane (Spectra Labs) and further concentrated to ~10-20 μM (~300 μl) using an Amicon Ultra-0.5 (10 kDa) centrifugal device at 4°C. For Sox2 titrations, 50 μM unlabeled Sox2 HMG was added slowly to NCPs at appropriate ratios and concentrated to ~300 μL as above. The (^15^N-H2A)-H2B or (^15^N-H2A.Z)-H2B dimer sample was prepared at ~20 μM by exchanging into NMR buffer, as for NCPs. To make the histone dimer-DNA complex, dimer was slowly added to a 2-fold excess of a DNA40 duplex containing the mFgf4 motif (see Table S1). The NMR buffer was supplemented with 7.5% D_2_O (CIL) prior to experiments. Data was collected on a Bruker Ascend 800 MHz NMR spectrometer equipped with a triple-resonance cryoprobe. Standard 2D ^1^H-^15^N HSQC experiments (64 scans) were collected at 25 °C and 37 °C for all NCP and histone-DNA samples. Standard 3D HNCO and HN(CA)CO assignment experiments were acquired with non-uniform sampling (20%) for histone-DNA complexes. Data was processed using nmrPipe^77^ and analyzed using Sparky (Goddard, T. D., & Kneller, D. G. **SPARKY 3**, University of California, San Francisco) and in-house scripts. H3 N-tail peak intensities were normalized to A15, a resolved uncharged residue with small cross-sample variation and no persistent DNA contacts in MD simulations; this normalization closely matched concentration-adjusted raw intensity differences between NCP samples. Chemical shift perturbations (CSPs) were calculated from the difference in proton (Δ δ_H_) and nitrogen (Δ δ_N_) chemical shifts in ^1^H-^15^N HSQC spectra, as follows: CSP = (Δ δH^2^ + (0.154Δ δ _N_)^2^)^1/2^.

### Molecular Dynamics (MD) simulation

One microsecond MD simulations were run using (i) the free Widom 601 nucleosome,^78^ (ii) the 601 nucleosome in complex with the Sox2 HMG domain at SHL-6,^65^ and (iii) the Lin28B^70^ nucleosome, each containing either H2A or H2A.Z histones. Three replicas were performed for each system. Summary of the MD simulation setup is reported in Table S5. The 601 DNA sequence and initial configuration of the nucleosome core particle (NCP) were obtained from the crystal structure PDB ID: 3LZ0^78^ missing the histone N-terminal tails and the C-terminal tail of H2A. The N-terminal tails and other missing residues were modeled following our previous work^79^. The canonical nucleosome containing H2A was modeled with the addition of the C-terminal tail (KTESSKSKSK), using Chimera.^80^ The dihedral angles for each residue in the C-terminal histone H2A tails were assigned with φ angle = -60° and ψ angle = 30°. The H2A.Z nucleosome was modeled using PDB ID: 1F66.^81^ by swapping canonical H2A from the above construct with H2A.Z. The Lin28B DNA sequence and initial configuration was obtained from the cryoEM structure PDB:7U0J.^70^ The Sox2 system was obtained from the cryoEM structure PDB:6T90,^65^ which contains Sox2 and Oct4 bound at the SHL-6 location. Oct4 was removed from the structure. The final NCP constructs were visualized using PyMOL and VMD.^82^ All nucleosome constructs were parameterized using AMBER forcefields at 150 mM NaCl concentration under physiological conditions. The histone was simulated with ff19SB,^83^ and DNA was parameterized using OL15.^84^ The OPC water model^85^ was used, with its Lennard-Jones interaction (Na^+^/OW) modification, using the Kulkarni *et al*. method, which provides better estimates of osmotic pressure.^86^ For sodium (Na^+^) and chlorine (Cl^−^) ions, Joung and Cheetham^87^ parameters were used. Mg^+2^ modification was performed using the Li *et al* parameter method.^88^ All systems were initially minimized and heated using Amber22.^89^ Systems were converted to GROMACS using ParmEd^90^, ensuring the correct ion parameters. Three replicas of all systems were equilibrated for 100 ns, followed by production runs for 1 µs using GROMACS 2022.5,^91^ except for the Lin28B systems, which were run on Anton 3.^92^ See SI Methods for more information on MD setup and analysis.

## Results

### H2A.Z stimulates DNA unwrapping in a sequence-dependent manner

Prior studies have reported potential interaction and functional coordination between histone variant H2A.Z and pioneer factors Sox2 and Oct4.^7, 21, 25, 26^ To probe the effect of H2A.Z on the interaction of Sox2 and Oct4 with chromatin, we constructed nucleosome core particles (NCPs) with a modified 145 base pair Widom 601 sequence (601*, where a non-specific Sox2 binding site is removed) containing Sox2-Oct4 composite motifs (Table 1 and Table S1). These motifs were derived from mouse Nanog (mNanog, N)^58^ and Fgf4 (mFgf4, F)^93^ gene regulatory elements and inserted at previously characterized positions at SHL+2, +5 and ±6.^66^ To assess the effect of H2A.Z in the context of naturally occurring, less stable nucleosomes, we used a minimal 147 base pair fragment from the human Lin28B enhancer that contains several putative Sox2 and Oct4 binding sites (Table1 and Table S1).^63^ The Lin28B dyad position was assigned based on a cryo-EM study by Guan *et al*.^70^ and the DNA fragment was truncated to 147 base pairs to minimize non-specific Sox2 interactions with linker DNA^66^ that could complicate binding analysis.

**Table 1.** Nucleosome constructs and corresponding genome origin, nucleotide sequence, and approximate nucleosome position (SHL) of the Sox2 binding site.

| NCP construct | Sox2 binding site |  |  |
| --- | --- | --- | --- |
|  | SHL | Gene | Sequence |
| **601\*** |  |  |  |
| 62<sub>F</sub> | SHL+6 | mFgf4 | CTTTGTT |
| 62<sub>N</sub> | SHL+6 | mNanog | CATTGTA |
| 54R<sub>F</sub> | SHL+5 | mFgf4 | CTTTGTT |
| 54R<sub>N</sub> | SHL+5 | mNanog | CATTGTA |
| 23R<sub>N</sub> | SHL+2 | mFgf4 | CTTTGTT |
| 23R<sub>F</sub> | SHL+2 | mNanog | CATTGTA |
| m62<sub>F</sub> | SHL-6 | mFgf4 | CTTTGTT |
| m62<sub>N</sub> | SHL-6 | mNanog | CATTGTA |
| **Lin28B** |  |  |  |
| Lin28B | SHL+6 | Lin28B | GCTTGTG |
|  | SHL+5 |  | GATTGTG |
|  | SHL+4 |  | AATTGTC |
|  | SHL-3 |  | TATTGTT |

We first asked whether H2A.Z would increase DNA mobility and unwrapping in our constructs, similar to previous reports.^51, 53^ To assess changes in DNA accessibility due to unwrapping, we combined nuclease digestion and bulk Förster resonance energy transfer (FRET) assays (Fig. 1). Digestion by Exonuclease III (ExoIII), which degrades DNA progressively from the 3’ end, was monitored over time by denaturing gel electrophoresis with base-pair resolution for each strand (Fig. 1A and Fig. S1). As a control for enhanced unwrapping, we generated corresponding NCP constructs with a histone H3 R49A mutation that eliminates a key histone-DNA minor groove interaction at nucleosome entry/exit points (Table S2).

**Figure 1.**
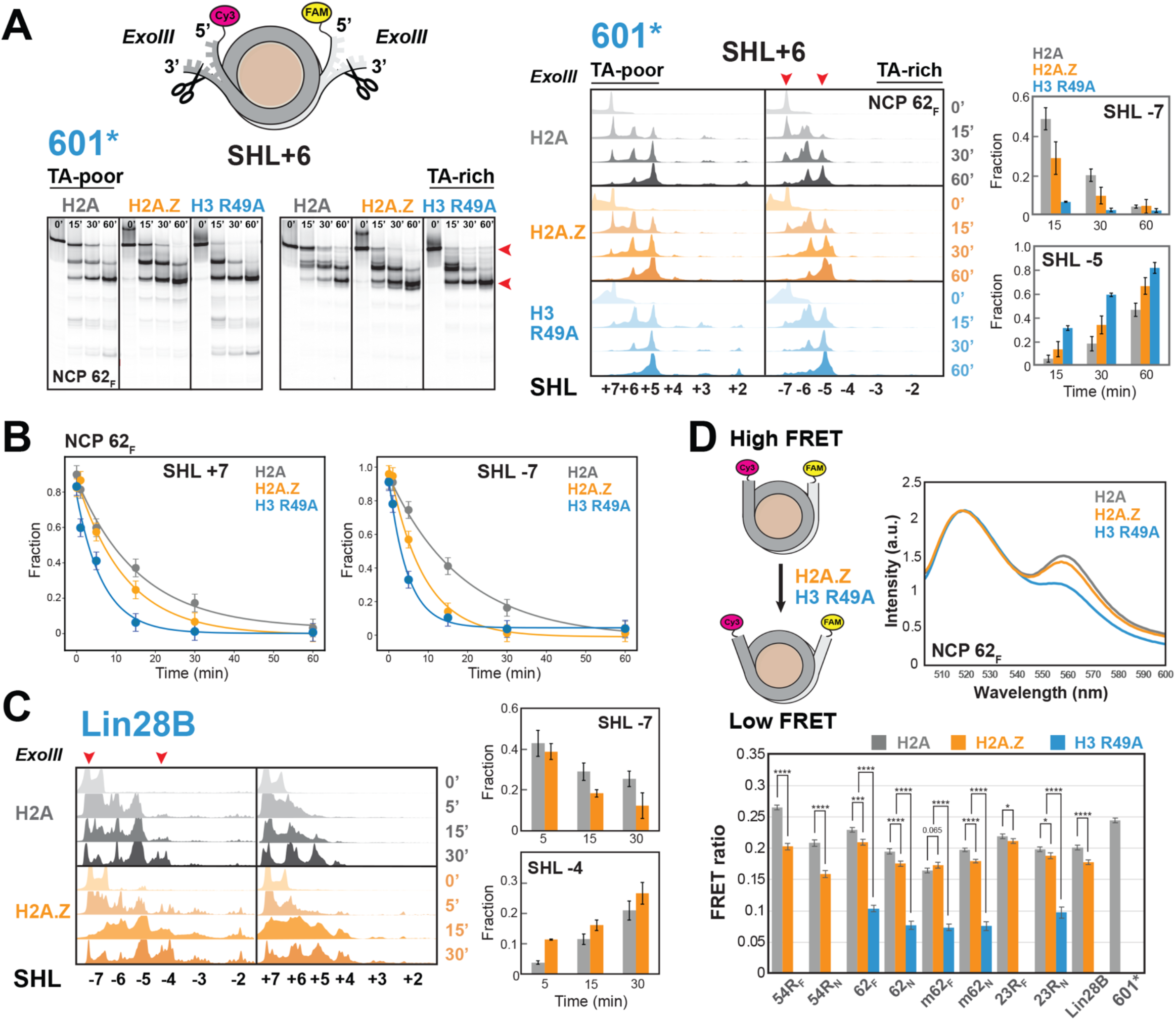
H2A.Z promotes DNA unwrapping in modified 601 and Lin28B nucleosomes. (A) Denaturing gel (left) and corresponding lane profiles (center) from an Exonuclease III (ExoIII) digestion time course of 601* NCPs containing a Sox2-Oct4 motif at SHL+6 (NCP 62_F_) with either H2A, H2A.Z, or an H3 R49A mutant. Quantified DNA fractions at SHL-7 and SHL-5 (indicated by red arrows) showing increased accessibility in H2A.Z and H3 R49A nucleosomes (mean ± s.d.). (B) ExoIII digestion profiles for the TA-poor (SHL+7) and TA-rich (SHL-7) sides of NCP 62F, along with monoexponential fits, showing faster decay with H2A.Z and H3 R49A. (C) ExoIII profiles for Lin28B NCPs, analyzed as in (A), showing enhanced DNA accessibility with H2A.Z. (D) FRET assay of nucleosomes labeled with FAM and Cy3 (left) to monitor DNA unwrapping. Representative fluorescence spectra (right) of NCP 62_F_ upon FAM excitation at 495 nm show reduced Cy3 emission at 560 nm (lower FRET) in H2A.Z and H3 R49A NCPs. FRET ratios (bottom, mean ± s.d.(total)) indicate increased DNA unwrapping for H2A.Z and, particularly, H3 R49A, with effects modulated by DNA sequence (601* denotes the NCP control lacking a Sox2-Oct4 motif). Statistical significance is indicated by *p*-values obtained from a two-tailed t-test(\*\*\*\**p*<0.0001;\*\*\**p*< 0.001;\*\**p*<0.01; \**p*<0.05).

Overall, ExoIII digestion was faster and more extensive with H2A.Z than with H2A for both 601* and Lin28B nucleosomes (Fig. 1, Figs S2 and S3, Table S3). In 601* constructs, the effect of H2A.Z on quantified digestion rates, reported in Table S3, was typically greater on the TA-rich side (SHL-7) of nucleosomes (Fig. 1). For instance, the 62_F_ NCP was digested twice as fast with H2A.Z (0.101 ± 0.026 min^−1^) as compared to H2A (0.050 ± 0.011 min^−1^) at SHL-7, but only 1.2-fold faster (0.082 ± 0.023 min^−1^ versus 0.066 ± 0.018 min^−1^) at SHL+7 (Fig. 1B). As expected, the control H3 R49A NCP displayed markedly faster ExoIII digestion (~0.20 ± 0.05 min^−1^ at SHL±7), consistent with greater unwrapping upon disruption of histone-DNA contacts (Fig. 1B). The effect of H2A.Z also varied across NCPs containing different sequences and placement of Sox2-Oct4 motifs, producing on average 0.8 to 2.0-fold changes in digestion rates (Figs S2 and S3, Table S3). For canonical Lin28B constructs, ExoIII digestion was significantly faster and penetrated deeper into the nucleosome core, indicating greater destabilization than 601* constructs (Fig. 1C). This effect, and especially the increased internal digestion extending to SHL2, was further amplified by H2A.Z. Together, these findings show that H2A.Z promotes DNA unwrapping in a sequence-dependent and asymmetric manner. This conclusion was independently supported by an endonuclease assay, in which the restriction enzyme MslI was used to cleave DNA sequence-specifically at the Sox2-Oct4 motif near SHL±6 (Fig. S1).

ExoIII digestion penetrated internal sites up to SHL+2 on the TA-poor side of H2A 601* nucleosomes, with the extent varying with Sox2-Oct4 motif position (Fig. S1). Surprisingly, this internal cleavage was suppressed in the presence of H2A.Z, but not with the destabilizing H3 R49A mutant, and persisted only when Sox2-Oct4 motifs were located at SHL+5. To test whether enhanced internal DNA mobility was involved, we employed an H3 R83A mutant, which selectively disrupts a histone Arginine-DNA minor groove contact at SHL±2. This mutant exhibited a localized increase in ExoIII digestion at SHL+2 relative to other internal sites (Fig. S1, NCP 23R_N_), consistent with changes in internal DNA dynamics. Together, these findings suggest that H2A.Z can destabilize nucleosomal DNA ends, while simultaneously stabilizing certain internal DNA regions, in a sequence-dependent manner. Our conclusion aligns with previous work showing that H2A.Z can both destabilize peripheral DNA and inhibit H2A-H2B dimer release from the H3-H4 tetramer during thermal denaturation of nucleosomes.^94^

To directly assess H2A.Z-induced DNA unwrapping, we measured FRET between FAM and Cy3 fluorophores covalently attached to the DNA 5′ ends (Fig. 1D). Increased unwrapping was expected to raise the average inter-probe distance, thereby reducing FRET efficiency. Consistent with this, FRET ratios were measurably lower with H2A.Z than with H2A in nearly all NCP constructs (Fig. 1D). The magnitude of the FRET change varied with the position and identity of the Sox2-Oct4 motif (Fig. 1D), being smaller when the Sox2-Oct4 motif was placed at SHL+2 and SHL6, but more pronounced at SHL+5. This pattern suggests that H2A.Z nucleosomes are particularly responsive to A-tract containing sequences at SHL+5, as found in the mFgf4 motif, consistent with the elevated internal ExoIII digestion observed for H2A.Z in NCP 54R_F_ (Fig. S1). Together, these results indicate that H2A.Z enhances entry/exit DNA unwrapping in a manner modulated by the underlying DNA sequence.

Finally, to obtain an atomistic view of changes in DNA conformation and dynamics elicited by H2A.Z, we performed three 1 μs MD simulations for 601 and Lin28B NCPs under physiological salt conditions (Fig. 2). H2A.Z containing systems exhibited increased overall mobility and DNA unwrapping, particularly at SHL-7 for 601 and both SHL±7 regions for Lin28B (Fig. 2A). This was accompanied by a significant increase in the average gapping distance between two parallel DNA gyres (Fig. 2B), increasing accessibility to internal regions. Our results are consistent with prior computational studies showing increased DNA unwrapping and gapping of Widom 601 and TP53(+1) nucleosomes, which were attributed to the altered conformation and dynamics of the H2A.Z C-terminal and N-terminal tails, respectively.^53^

**Figure 2.**
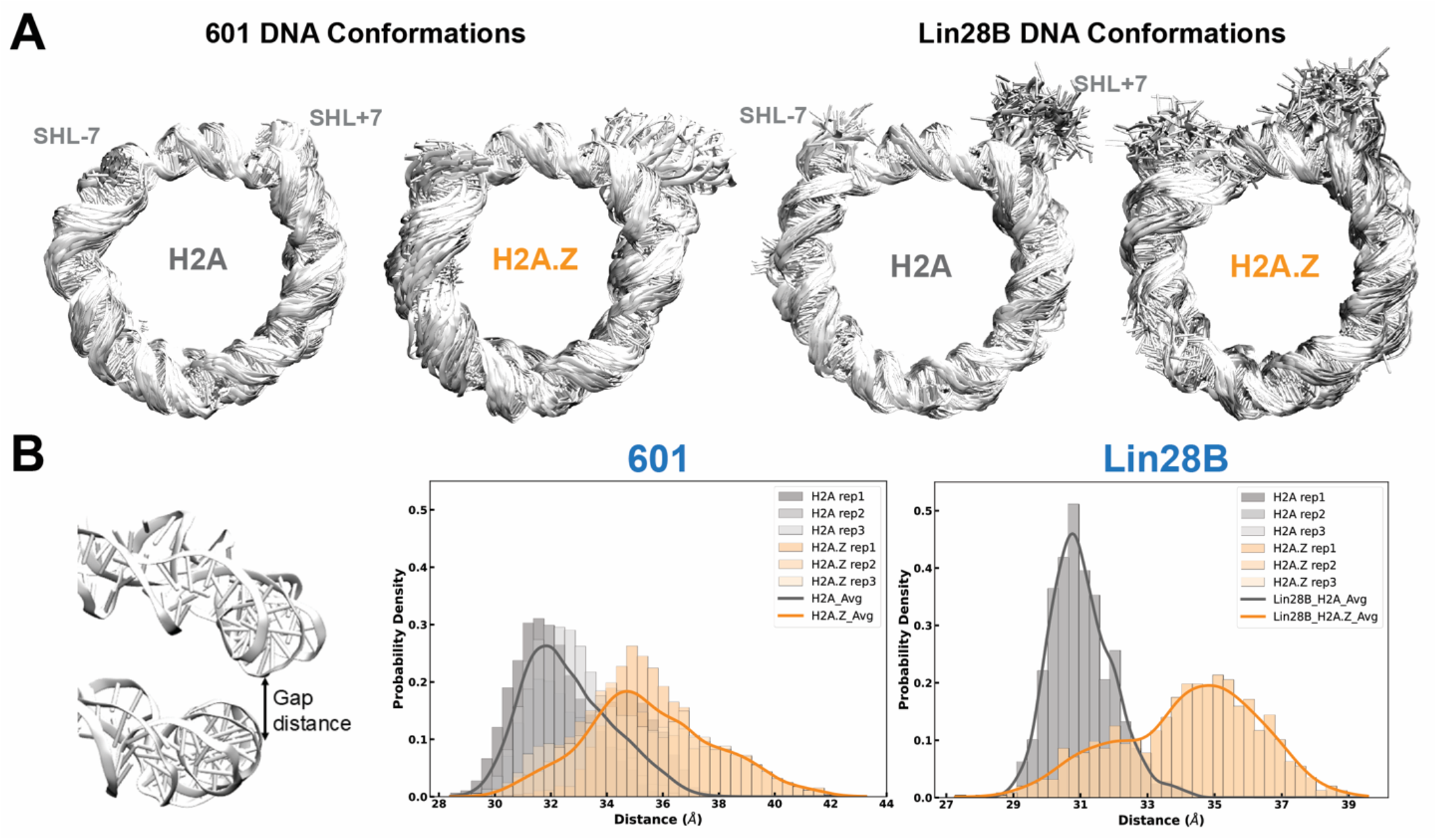
H2A.Z increases DNA unwrapping and inter-gyre distance. (A) The fluctuation of nucleosomal DNA in superimposed conformations for H2A (left) and H2A.Z (right) in MD simulations of 601 and Lin28B nucleosomes, showing larger DNA unwrapping and overall mobility with H2A.Z. (B) Density plots of the gap distance between two DNA gyres in H2A (grey) and H2A.Z (orange) 601 and Lin28B nucleosomes showing a significant increase with the H2A.Z variant.

### H2A.Z alters the conformational dynamics and disrupts DNA interactions of the H3 tail

While the increase in DNA mobility has been previously linked to specific H2A.Z tail dynamics, the impact of the histone H3 N-terminal tails (N-tail), which sit at the nucleosome edge, has not been investigated. To investigate how H2A.Z affects H3 N-tail conformation and dynamics, we first employed solution NMR spectroscopy. We assembled nucleosomes containing either H2A or H2A.Z, each with a single uniformly ^15^N-labeled H3 with the Sox2-Oct4 composite motif positioned at SHL+6 (NCP 62_F/N_), SHL+5 (NCP 54R_F_) or SHL+2 (NCP 23R_N_). In samples prepared at 20 μM, standard ^1^H-^15^N HSQC spectra readily revealed backbone amides from the flexible N-terminal tails of labeled histones.

We compared the backbone amide HSQC spectra of the H3 N-terminal tails of H2A and H2A.Z NCPs (Fig. 3A, Fig. S5). Using published assignments,^95^ we quantified changes in peak intensities and chemical shifts to assess changes in backbone structure and dynamics (Fig. 3B, Fig. S5). While chemical shifts remained largely unaffected (Fig. S5D), we observed more sizable changes in the relative intensities for two H3 N-tail regions – residues 10-13 and 22-34 – particularly for NCP 62_F_ and 54R_F_ containing the mFgf4 motif (Fig. 3B). These regions primarily consist of residues that were previously characterized by NMR as flexible “hinges” due to their higher mobility, flanked by Arg “anchor” residues (Fig. 3B, highlighted in red) that exhibit lower mobility and maintain stronger DNA interactions.^96^ For NCP 62_F_ and 54R_F_, multiple peak intensities were reduced in H2A.Z as compared to H2A NCPs (Fig. 3B), most consistent with increased chemical exchange dynamics on the micro-to-millisecond (*μs-ms*) timescale. Intensities were further attenuated with less stable constructs, a C-tail swap mutant (H2A.Z^C-H2A^ NCP 54R_F_) and, particularly, H3 R49A (NCP 62_F_) (Fig. 3B). This data suggests a correlation between altered H3 N-tail dynamics and increased DNA unwrapping associated with these mutants. One possible scenario is that H2A.Z-induced DNA unwrapping alters N-tail conformational sampling or exposes new DNA surfaces, breaking or redistributing transient tail-DNA contacts. Simultaneously, the selective intensity decrease with H2A.Z cannot be explained solely by an increase in overall tumbling due to larger DNA unwrapping, which would typically influence all residues, nor by increased conformational rigidity resulting in higher transverse relaxation rates (R_2_), which would oppose the expected effect from disrupting histone tail-DNA interactions with unwrapping.

**Figure 3.**
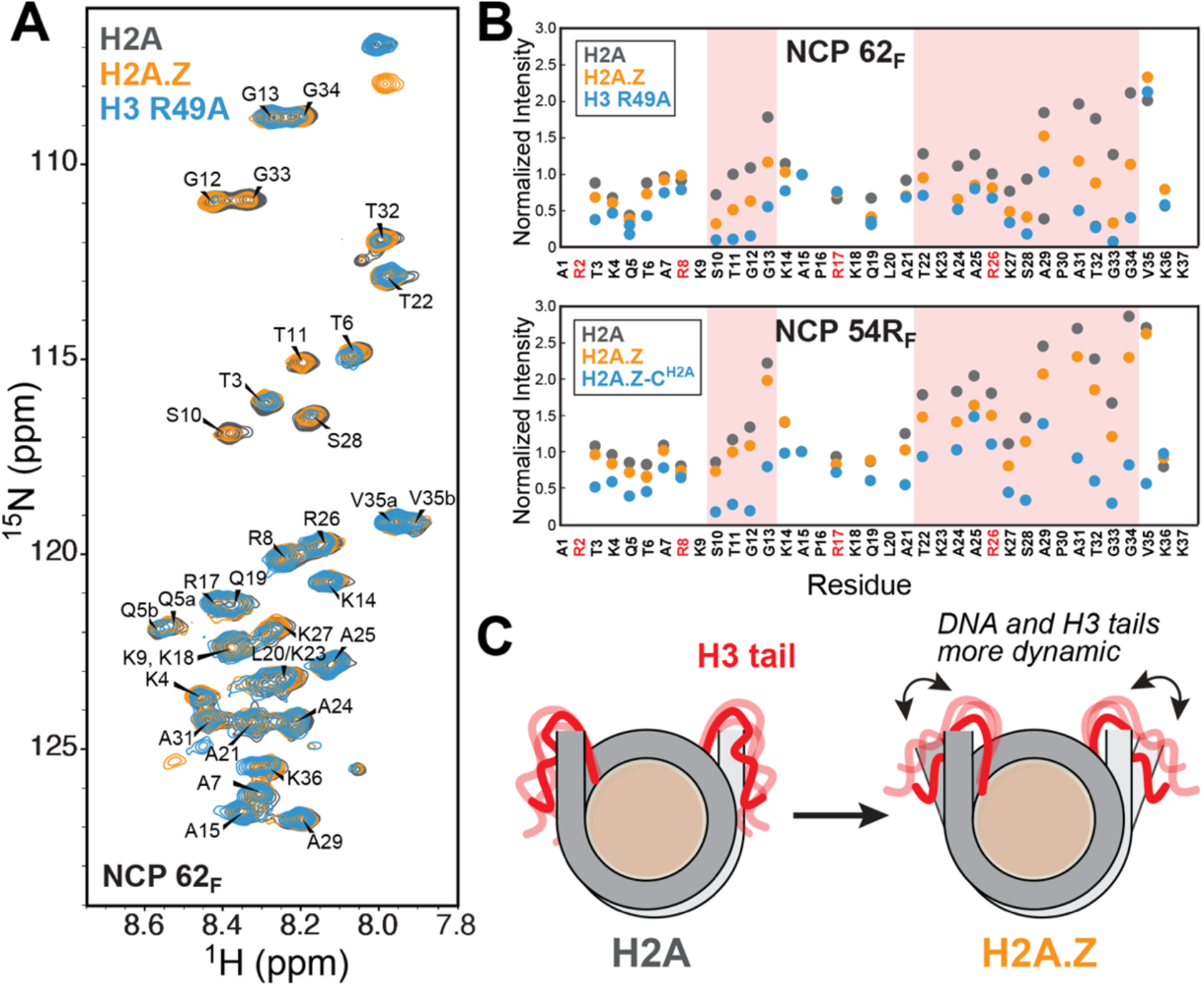
H2A.Z perturbs the dynamics of the H3 N-terminal tail. (A) Overlay of ^1^H-^15^N HSQC spectra of ^15^N-H3 NCP 62_F_ (20 μM) containing H2A, H2A.Z or H3 R49A (37°C) showing differences in peak intensities and chemical shifts. (B) Normalized peak intensities obtained from spectra in (A) (top) showing decreased values for residues in flexible “hinge” regions (S10-G13, T22-A25, S28-G34) with H2A.Z and, especially, with H3 R49A. Similar results are obtained for NCP 54R_F_ (bottom) with H2A.Z and the destabilizing H2A.Z^C-H2A^ mutant (see Fig. S14). (C) Cartoon of H3 N-tails showing perturbed dynamics in the presence of H2A.Z.

For nucleosomes containing the mNanog motif (NCP 62_N_ and 23R_N_), the same regions showed instead an increase in peak intensities with H2A.Z, an effect further amplified by the destabilizing H2A.Z^C-H2A^ (NCP 62_N_) mutant (Fig. S5). This can be interpreted as enhanced fluctuations on the picosecond-to-nanosecond (*ps-ns*) timescale, faster than the predicted rotational correlation time for a free nucleosome (~110 ns),^97^ which are known to dominate H3 tail dynamics in solution.^98^ The opposing intensity trends in mNanog versus mFgf4 NCPs likely reflect differing contributions of slow *(μs-ms*) and fast (*ps-ns*) dynamics. In this case, H2A.Z-induced unwrapping likely disrupts specific transient H3 tail-DNA contacts, reducing R_2_and narrowing linewidths through elimination of chemical exchange between DNA-bound and unbound states and increased local flexibility. We note that, based on FRET efficiencies (Fig. 1D), mNanog NCPs are on average more unwrapped than mFgf4 ones, which could correlate with altered H3 N-tail dynamics. Low sample concentrations here prevented reliable NMR relaxation measurements of *ps-ns* dynamics. Further work will be required to quantitatively characterize these motions across timescales and assess their dependence on DNA sequence and histone variant.

To gain further insight into H3 N-terminal tail dynamics associated with DNA unwrapping in H2A.Z NCPs, we examined H3 tail behavior in our MD simulations. In H2A 601 and Lin28B NCPs, the H3 N-tail adopted more extended conformations that bridged the two DNA gyres and maintained stable contacts with SHL2 (Fig. 4B and Fig. S6). This bridging may suppress DNA unwrapping and inter-gyre gapping in nucleosomes by both enabling direct H3-DNA interactions and screening the electrostatic repulsion between adjacent DNA gyres. By contrast, in the presence of H2A.Z, the H3 N-tail adopted altered conformations that were repositioned away from SHL2 (Fig. 4B and Fig. S6) and were slightly less ordered when compared H2A (e.g. 3.5% versus 2% β-sheets and 3% versus 2.7% α-helices for 601 NCPs). Likewise, ensembles for the H2A.Z C-terminal tail showed increased dynamics and reduced DNA interactions (Fig. 4C and Fig. S6). To further correlate H3 structural dynamics with our NMR results, we calculated root-mean-square fluctuations (RMSF) (Fig. 4D and Fig. S7). In H2A.Z systems, H3 consistently exhibited greater RMSF values, particularly in the N-terminal regions. For the 601 NCPs, the largest N-tail fluctuations associated with greater unwrapping (SHL+7), occurred in residues 13-16 and 23-33 (Fig. 4D). This pattern is consistent with our NMR data, which show pronounced intensity changes for a similar set of residues (Fig. 3 and Fig. S5).

**Figure 4.**
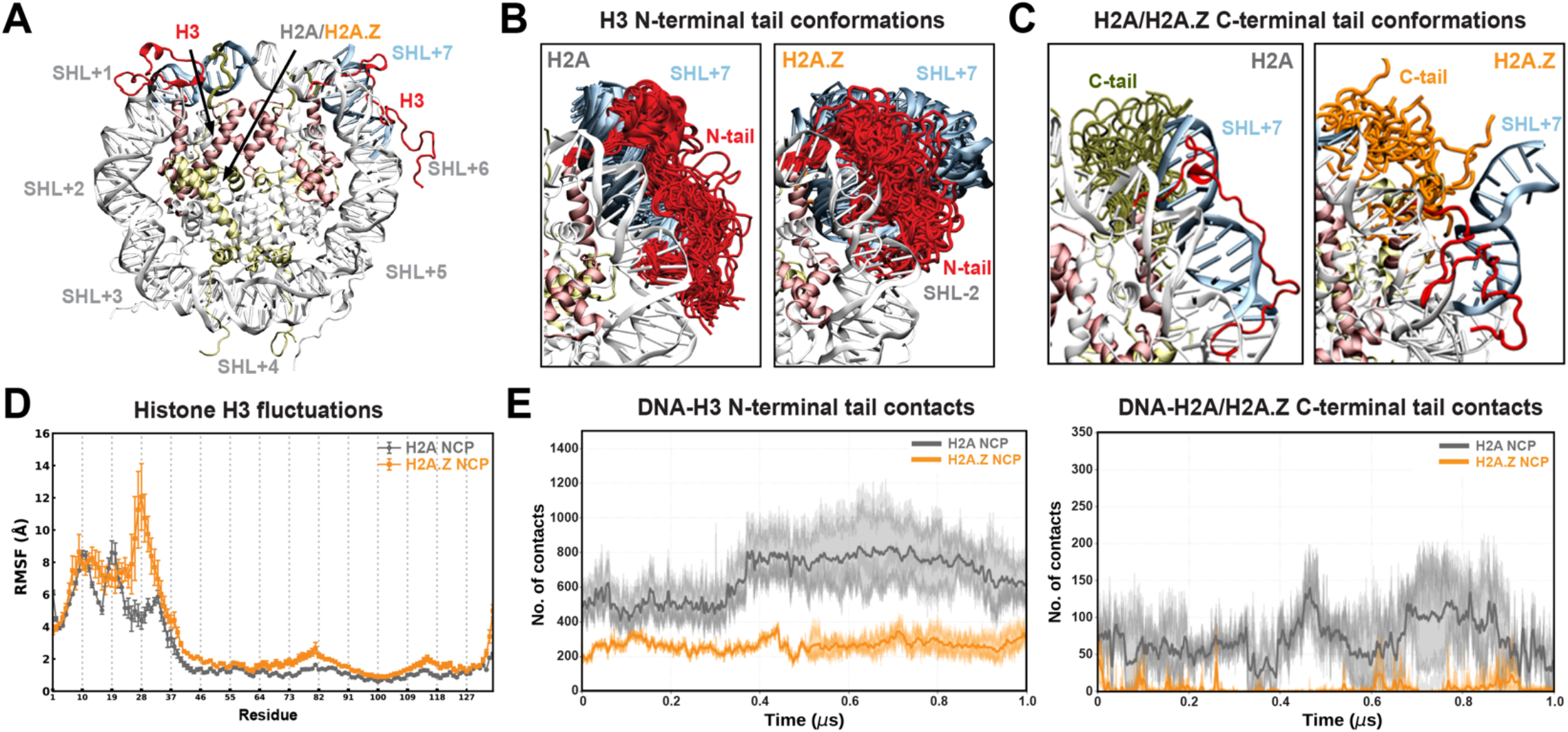
DNA unwrapping in H2A.Z nucleosomes is coupled to increased H3 fluctuations and fewer tail-DNA contacts. (A) The modeled structure of NCP with superhelical locations (SHL+), highlighting H2A (yellow), H3 (red), and DNA at SHL+7 (ice blue). (B) Conformations of H3 N-terminal tails (red) during the MD simulations depicting higher disorder and altered conformations with H2A.Z, shifted away from SHL-2. (C) Conformations of H2A/H2A.Z C-terminal tails (green/orange) during the MD simulations. (D) Root-mean-square fluctuations (RMSF) for Cα atoms of histone H3 for H2A (grey) and H2A.Z (orange) systems (mean ± s.d. of 3 replicas) showing overall greater fluctuations with H2A.Z, particularly in the H3 N-tail. (E) Total contacts (4.5 Å cutoff) between the H3 N-tail (left) or the H2A/H2A.Z C-tail (right) and DNA as a function of time (averaged over 3 replicas), showing fewer contacts in the H2A.Z (orange) as compared to the H2A (gray) system.

Analysis of the total contacts between the H3 N-tail and DNA in MD trajectories revealed substantially fewer interactions with H2A.Z versus H2A in 601 NCPs (Fig. 4E and Fig. S8A), with a smaller decrease in Lin28B NCPs (Fig. S9A). Similarly, contacts between the H2A.Z C-tail and DNA were significantly lower than for the H2A C-tail, consistent with previous reports^53^ and with its shorter length and fewer positively charged residues (Fig. 4E, Figs S8B and S9B). Closer inspection revealed that key hydrogen bonds between positively charged H3 N-tail residues (i.e. R17, R26, and R42) and the DNA backbone were impaired with H2A.Z (Fig. S10). Also, residues surrounding H3 R26 exhibited disrupted DNA interactions, consistent with NMR intensity changes and elevated RMSF values. We also observed several contacts between H3 R42 and the H2A C-tail, including a potential salt bridge between H3 R42 and H2A E121 (Fig. S11), observed in prior structural studies,^99^ that would not be possible with H2A.Z. Together, our results support a model where the H2A.Z variant disrupts key contacts between the H3 N-tail, H2A C-tail, and DNA, increasing DNA mobility and accessibility through enhanced unwrapping and inter-gyre spacing.

### H2A.Z enhances Sox2 and Oct4 binding to modified 601 nucleosomes

We next asked whether the H2A.Z-induced changes in nucleosome conformation and dynamics affect the binding of pioneer factors Sox2 and its functional partner Oct4. To test this, we performed native gel shift assays (EMSA) under near-physiological salt conditions (150 mM KCl) (Fig. 5 and Fig. S12). When Sox2-Oct4 motifs were positioned near the nucleosome edge, at SHL-6 (NCP m62_F_, m62_N_) or SHL+6 (NCP 62_F_, 62_N_) of 601*, H2A.Z visibly increased the stability of the Sox2-NCP complex (Fig. 5B and Fig. S12). This enhancement was observed for both mFgf4 (F) and mNanog (N) motifs and was further amplified in the more unwrapped H3 R49A control. Sox2 binding to H2A.Z nucleosomes was also strengthened when the motif was placed internally at SHL+2 (NCP 23R_F_, 23R_N_) (Fig. 5B). The increased internal mobility at SHL+2 with the H3 R83A mutant similarly promoted Sox2 binding. In contrast, H2A.Z had less detectable impact on Sox2 binding at a high-affinity site near SHL+5 (NCP 54R_F_, 54R_N_) (Fig. S12).^66^ Together, these results indicate that H2A.Z enhances Sox2-NCP binding in a position-dependent manner at both internal and terminal sites. This is consistent with increased DNA accessibility at SHL6 and SHL2, driven by the enhanced unwrapping, inter-gyre gapping, and elevated H3 N-tail dynamics described above.

**Figure 5.**
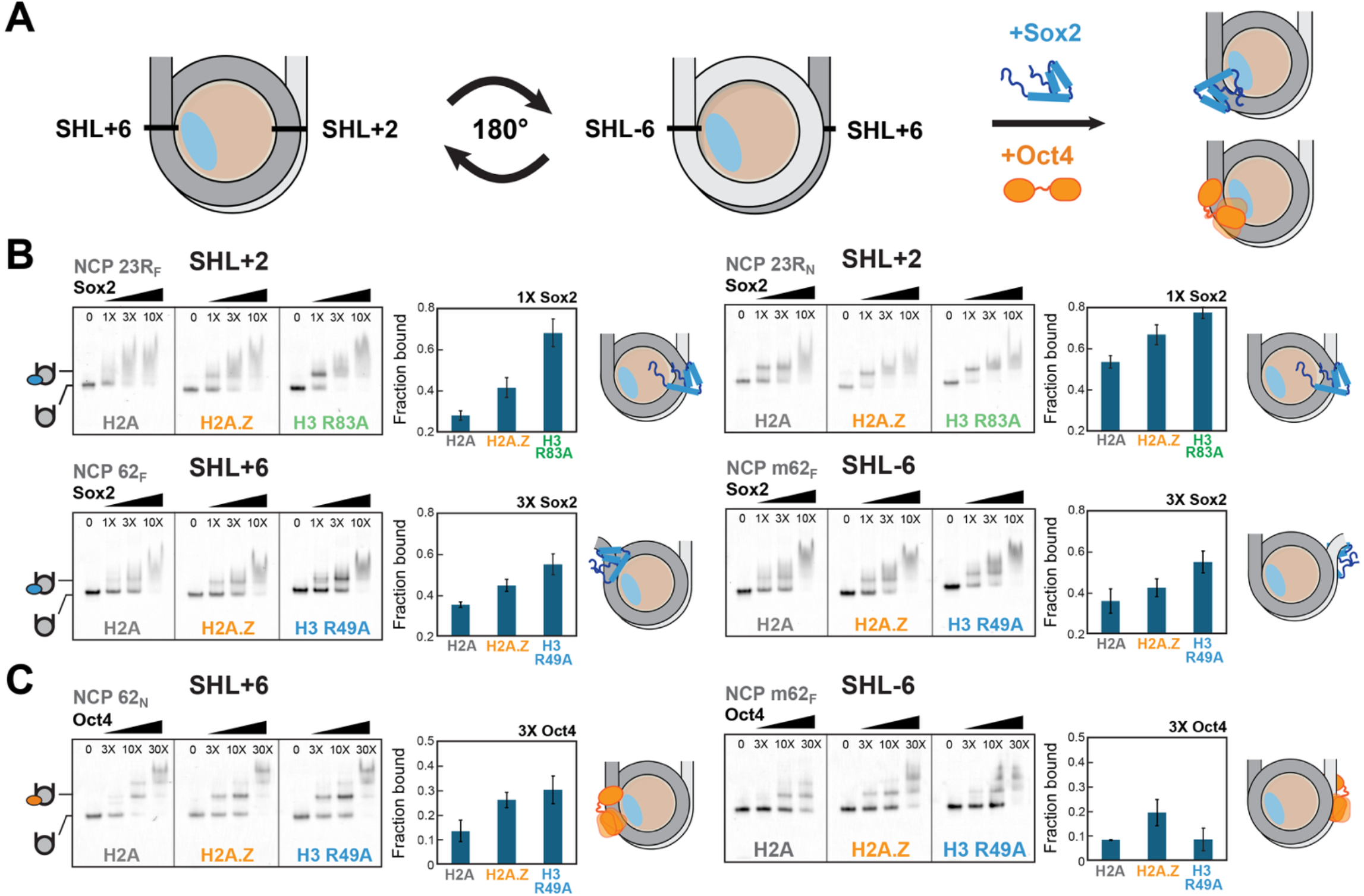
H2A.Z enhances Sox2 and Oct4 binding at end-positioned and internal motifs on 601* nucleosomes. (A) Approximate position of Sox2-Oct4 composite motifs (black lines) near SHL+2, +6, and -6 shown in relation to H2A/H2A.Z (light blue). (B) EMSA analysis of NCPs (10 nM) (left panels) with increasing Sox2 concentrations (1, 3, 10X) showing enhanced Sox2 binding with H2A.Z (SHL+2 and SHL±6) and, especially, with H3 R83A and H3 R49A mutants, used as controls for locally elevated DNA mobility at SHL2 and SHL6. Quantified fractions of Sox2-bound NCPs (right panels) represent mean ± s.d. (C) EMSA analysis of NCPs (10 nM) incubated with increasing concentrations of Oct4 (3, 10, 30X) showing formation of a more stable specific Oct4-NCP complex with H2A.Z and H3 R49A at SHL+6 (TA-poor side).

Our results also point to a potential role of H2A.Z-induced nucleosome dynamics in Oct4-nucleosome binding (Fig. 5C and Fig. S12). For NCP 62_N_ (SHL+6, TA poor side), where the Oct4 mNanog POU_S_half-site is exposed and the POU_HD_half-site occluded, H2A.Z promoted the formation of the specific Oct4-NCP complex (Fig. 5C). By contrast, when the Oct4 motif was placed at SHL-6 (NCP m62_N_) on the TA-rich side, which we observed to be more unwrapped by ExoIII digestion (Fig. S1, S2 and S3), the specific complex formed equally well on canonical and H2A.Z NCPs (Fig. S12). This was also noted with the destabilizing H3 R49A mutant (Fig. 5C). Constructs that contained the mFgf4 motif at SHL±6 (NCP 62_F_, m62_F_) where the POU_HD_ half-site is solvent accessible, only H2A.Z enhanced formation of the specific complex (Fig. 5C and Fig. S12). The pronounced effect of H2A.Z, coupled with the lower sensitivity to DNA unwrapping (H3 R49A), suggests that Oct4 may adopt a distinct binding mode.

Using FRET, we further detected Sox2-induced deformation of nucleosomal entry/exit DNA as reduced FRET ratios (Fig. S4). This deformation was enhanced by the synergistic binding of Sox2 and Oct4 and became more pronounced with H2A.Z. These observations reinforce our previously published findings that the folded conformation of Sox2, and accompanying large DNA bending, is stabilized by Oct4 on adjacent motifs at SHL6.^66^

### H2A.Z modulates an altered Lin28B nucleosome conformation in the presence of Sox2

The Lin28B nucleosome is more prone to unwrapping than 601, as seen by ExoIII digestion (Fig. 1C) and MD simulations (Fig. 2), and is known to exhibit greater dynamics^31^ and heterogenous positioning *in vitro*.^52^ Surprisingly, in our assays, H2A.Z had little effect on Sox2 binding to Lin28B NCPs (Fig. 6B). EMSA revealed a major super-shifted band at an equimolar ratio of NCP and Sox2, with multi-site binding evident at higher Sox2 concentrations. Similar binding profiles were observed for Oct4, with only a modest enhancement in the presence of H2A.Z (Fig. 6B). The lack of substantial H2A.Z effect could be due to the inherently large Lin28B NCP dynamics with H2A or may reflect the suboptimal sequence, orientation or positioning of the Sox2 and Oct4 motifs (Fig. 6A).

**Figure 6.**
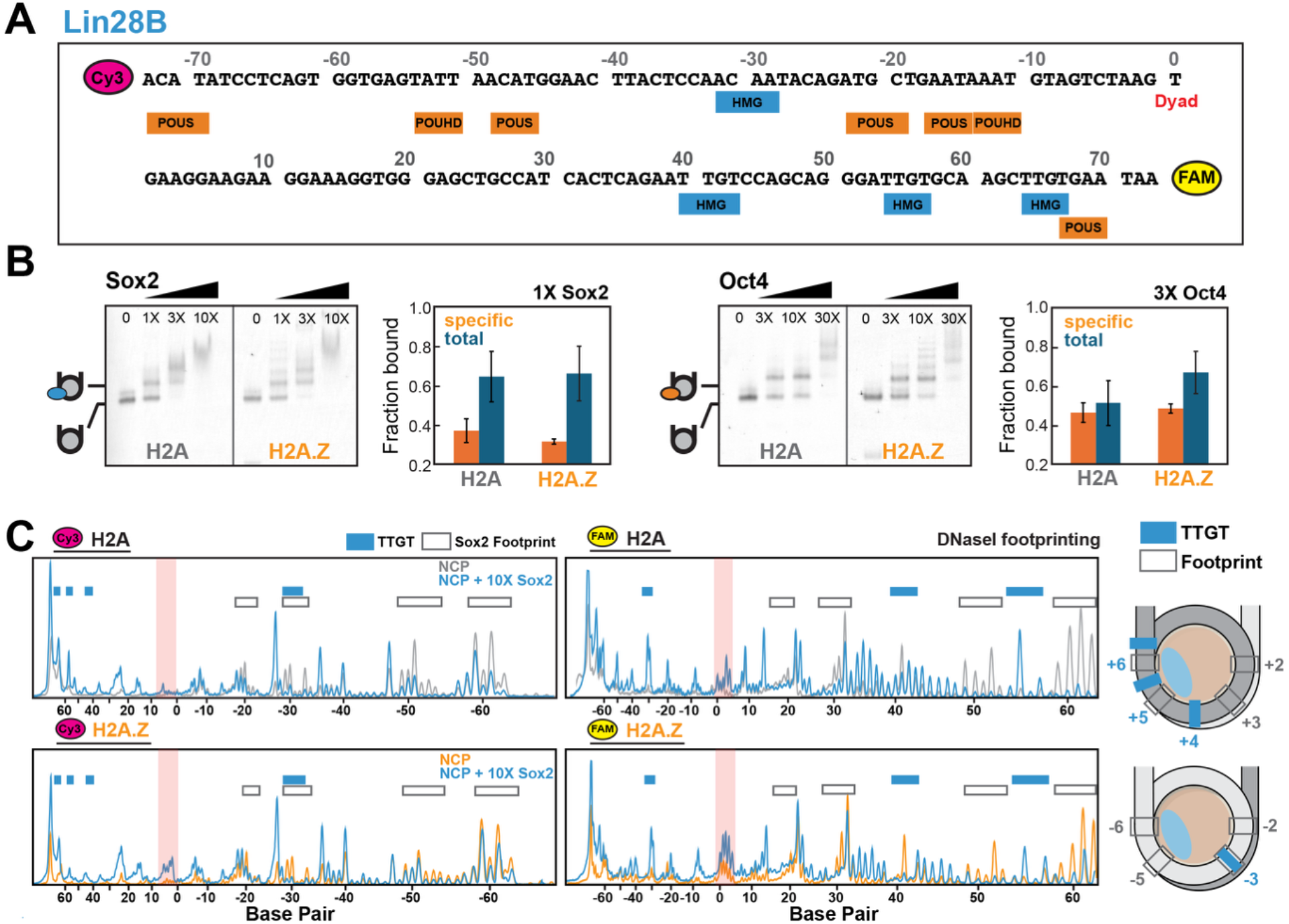
H2A.Z modulates an altered Lin28B NCP conformation in the presence of Sox2. (A) Lin28B sequence depicting potential binding motifs (TTGT) for Sox2 HMG (blue) and Oct4 POU_S_ and POU_HD_ domains (orange). (B) EMSA gels of nucleosomes with increasing Sox2 (left) or Oct4 (right) concentrations and quantified fraction bound (specific complex or total, mean ± s.d.) showing similar binding with both H2A and H2A.Z. The laddering effect with H2A.Z could be an artifact of some free DNA present. (C) DNaseI digestion profiles of H2A and H2A.Z Lin28B nucleosomes (100 nM) with or without Sox2 (10X), showing multiple footprints (decreased signal) and hypersensitive sites (increased signal) with Sox2. The location of Sox2 footprints (grey, open) and TTGT motifs (blue) are indicated. The nucleosome dyad region, showing higher DNaseI and Sox2 sensitivity with H2A.Z, is highlighted in red. NCP cartoons (right) depicting approximate locations of Sox2 motifs and DNaseI footprints, and the position of H2A/H2A.Z (light blue).

To further map Sox2 and Oct4 binding sites on Lin28B nucleosomes, we performed limited DNaseI digestion (Fig. 6C and Fig. S13). Lin28B contains four minimal Sox2 motifs (TTGT) (Fig. 6A) and is known to interact extensively with Sox2.^63^ Both H2A and H2A.Z NCPs displayed alternating regions of reduced DNaseI cleavage (footprints) and increased cleavage (hypersensitive sites), observed at two DNaseI concentrations and unlikely to result from over-digestion (Fig. 6C and Fig. S13A). These Sox2-dependent patterns were absent in naked DNA, no-DNaseI controls, or in 601* NCPs (Fig. S13A,B). Footprints appeared periodically at outward-facing DNA minor groove sites, often extending beyond TTGT motifs, whereas cleavage maxima occurred at inward-facing minor grooves (Fig. 6C). H2A.Z NCPs showed a similar pattern but with enhanced cleavage at the dyad and SHL±1, which became more pronounced with Sox2 binding and was absent in 601* constructs (Fig. S13A).

Oct4 further enhanced the effect of Sox2 in DNaseI assays (Fig. S13A, compare lanes 2 and 4), consistent with cooperative Oct4-Sox2 binding.^66^ While Oct4 alone showed ubiquitous binding in EMSA and DNaseI digestion, it did not generate hypersensitive sites. Cryo-EM structures of Lin28B NCPs with one or three Oct4 molecules likewise showed no major DNA repositioning, apart from asymmetric DNA unwrapping.^70^ Thus, the conformational change in Lin28B nucleosomes is unique to Sox2 and not purely due to DNA unwrapping.

While Sox2 produced numerous footprints on Lin28B consistent with the multi-site binding by EMSA, we did not observe the cleavage expected from binding or nearby DNA deformations (Fig. S13C). Hypersensitive sites appeared semi-regularly at predicted buried minor groove positions (Fig. 6C), rather pointing to changes in nucleosomal DNA mobility or positioning. This is supported by increased ExoIII digestion of Lin28B NCPs in the presence of Sox2 (Fig. S13D). One possibility is that Sox2 shifts the DNA register in less stable nucleosomes such as Lin28B, consistent with prior coarse-grain MD simulation studies of Sox2-Lin28B complexes.^32^ Our initial attempts to detect nucleosome deformation or sliding via histone-DNA crosslinking at SHL±5 were unsuccessful, likely due to extensive unwrapping and low crosslinking efficiency. Further studies with more optimal NCP constructs are currently underway.

### H2A.Z C-terminal tail and L1-loop differentially regulate nucleosome dynamics

The H2A.Z C-tail, which shows higher fluctuations and reduced DNA contacts in simulations, has been linked to increased nucleosome entry/exit dynamics.^51, 53^ To test whether the C-tail promotes unwrapping and contributes to enhanced Sox2 binding, we generated reciprocal C-tail swaps (Table S2, Fig. S14A). Installing the H2A.Z C-tail onto H2A (H2A^C-H2A.Z^) increased DNA internal dynamics and unwrapping in 601* and Lin28B NCPs, as shown by the enhanced ExoIII internal digestion (at SHL+2, +3, and +4) and reduced FRET (Fig. S14B-E). Surprisingly, replacing the H2A.Z C-tail with that of H2A (H2A.Z^C-H2A^) also destabilized nucleosomes, indicating that residues outside the tail influence dynamics (Fig. S14B). Despite elevated unwrapping, H2A^C-H2A.Z^ did not seem to improve Sox2-NCP stability by EMSA (Fig. S14F). This suggests the C-tail alone cannot account for H2A.Z-dependent Sox2 binding, consistent with the broader increase in H2A.Z mobility observed computationally.^53^

To probe additional structural contributions, we introduced two other H2A.Z-to-H2A swaps targeting the N-terminal tail (H2A.Z^N-H2A^) and the L1 loop (H2A.Z^L1-H2A^) (Table S2 and Fig. S14A). Both L1-loop and C-tail mutants exhibited enhanced internal digestion on the TA-poor side (Fig. S14B), which was normally suppressed by H2A.Z (Figs S1 and S14B). These mutants also showed significantly reduced FRET relative to H2A and H2A.Z constructs for NCP m62_N_and m62_F_, but not for NCP 62_F_ (Fig. S14C), indicating that perturbations in these regions are DNA-sequence dependent and can modulate NCP dynamics. The H2A.Z L1 loop, proposed to alter dimer orientation and reinforce hydrogen bonding networks between histones and DNA,^54^ may underlie part of the internal H2A.Z stabilization lost in these mutants. Further targeted mutagenesis and computation will be required to pinpoint the specific residues and allosteric pathways responsible for these effects.

### Sox2 displaces the H3 N-terminal tail from DNA at multiple nucleosome sites

We hypothesized that the reduced H3 N-tail contacts with DNA in H2A.Z nucleosomes may facilitate Sox2 binding. To investigate how Sox2 affects the conformation and dynamics of the H3 N-tail, we performed titrations of canonical and H2A.Z NCPs with Sox2 and monitored H3 amide chemical shifts (Fig. 7 and Fig. S15). The largest chemical shift perturbations (CSPs) were observed when Sox2 bound at SHL+6 (NCP 62_F/N_), near the H3 N-tail-DNA interaction site (Fig. 7B). Notably, we detected the emergence of a second set of peaks shifted in the direction of the free H3 N-tail peptide or higher salt ^42^ (Fig. 7 and Fig. S15), consistent with Sox2 competitively displacing one of the H3 N-tails from DNA (Fig. 7C). Residues at and around Arg “anchor” points (i.e. R26, R17, R8) that maintain stronger DNA interactions, particularly H3 R26, exhibited the largest CSPs (Fig. 7B). Similar trends were observed for H2A.Z and H3 R49A NCPs, with generally larger CSPs at and around R26 (Fig. 7B and Fig. S15).

**Figure 7.**
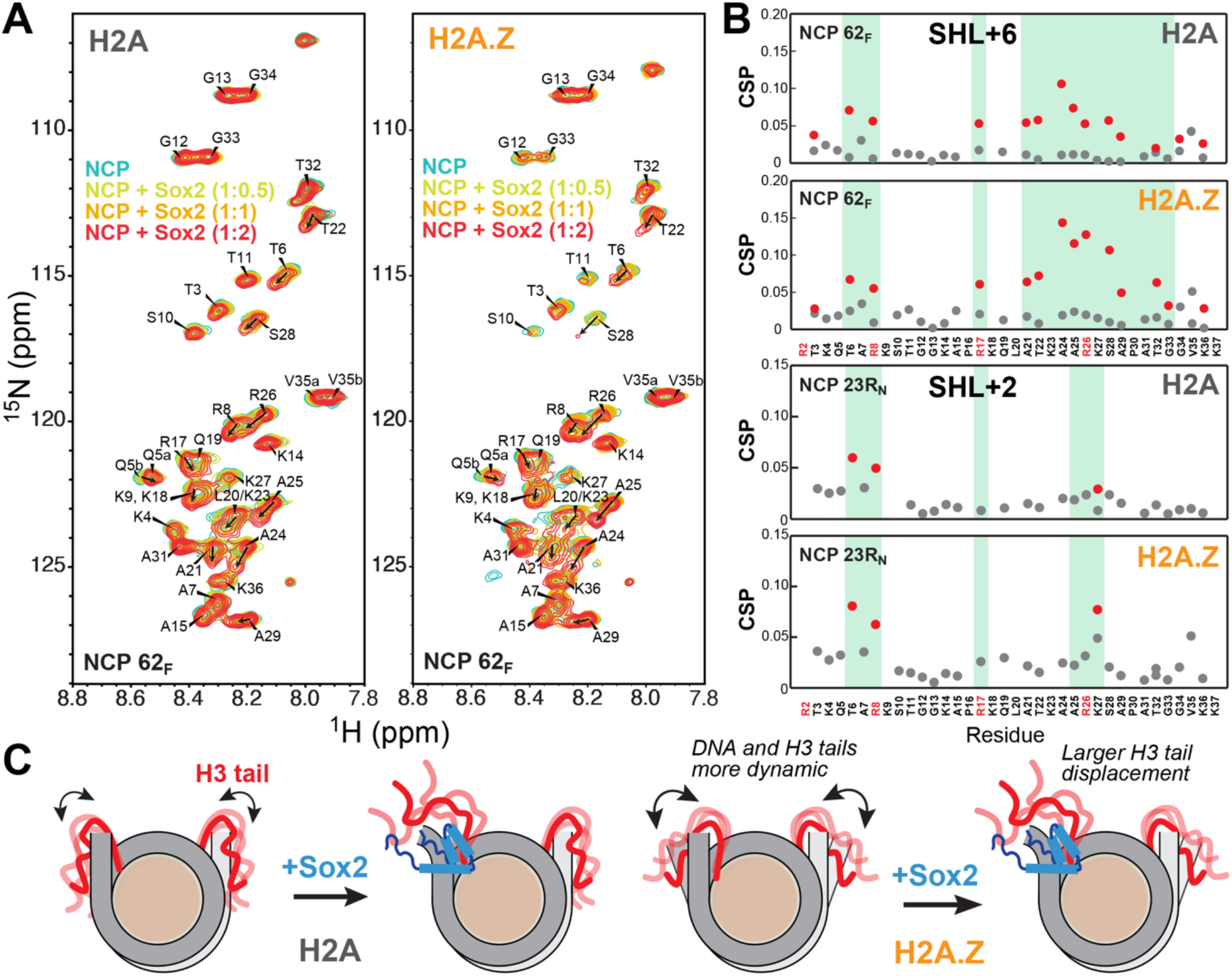
H2A.Z facilitates Sox2-mediated displacement of the H3 N-terminal tail. (A) ^1^H-^15^N HSQC spectral overlay of NCP 62_F_ (20 μM) containing ^15^N-H3 and H2A (left) or H2A.Z (right) with increasing concentrations of Sox2 (10, 20, and 40 μM) obtained at 37°C. Visible peaks represent the H3 N-terminal tail (residue 2-36). A second population of peaks is observed with Sox2 with chemical shifts closer to the free H3 N-tail peptide,^42^ consistent with tail dissociation from DNA. (B) Chemical shift perturbations (CSPs) for the H3 N-tail derived from Sox2 titrations of NCP 62_F_ (top) and NCP 23R_N_ with H2A or H2A.Z calculated at 1:2 NCP:Sox2 ratio. The second population is shown with red circles. Large CSP values for residues around R8 and R26, which tend to become more pronounced with H2A.Z, are highlighted in green. (C) Cartoon showing H3 N-tail displacement upon Sox2 binding, facilitated by enhanced tail dynamics with H2A.Z.

Interestingly, we also observed smaller but distinct chemical shift changes in the H3 N-tail when the Sox2 motif was positioned at internal sites in 601* NCPs, SHL+5 (NCP 54R_F_) or SHL+2 (NCP 23R_N_), or with Lin28B NCPs (Fig. 7B and Fig. S15). For these constructs, only a few peaks split into two discernible populations, consistent with more limited conformational changes. Notably, chemical shifts for Lin28B without Sox2 were shifted more towards the free H3 peptide relative to 601*, indicating partial H3 N-tail dissociation. Overall, a greater effect was detected with H2A.Z and destabilizing mutants. For NCP 23R_N_, large CSPs were clustered around the first few residues (Fig. 7B), suggesting that Sox2 mainly displaces the tip of the H3 N-tail. Because SHL+2 lies near the edge of the adjacent DNA gyre (SHL-7), Sox2 binding could directly or allosterically disrupt interactions between DNA and the H3 N-tail, observed in MD simulations (Fig. S10). Thus, our NMR data suggest that Sox2 competes with the H3 tails for DNA binding at multiple nucleosome positions, which could affect its interaction with nucleosomes and impact chromatin structure and histone tail recognition.

### Sox2 more strongly perturbs the H2A C-terminal tail than for H2A.Z

We next asked if Sox2-nucleosome association at SHL6 and SHL2 could be affected by competition with other histone tails, including the H2A C-tail and H4 N-tail. To test this, we performed titrations of canonical and H2A.Z nucleosomes containing either ^15^N,^13^C-labeled H2A/H2A.Z or ^15^N-labeled H4 with Sox2 (Fig. 8, Figs S16 and S17). Resonance assignments for H2A and H4 in NCPs were transferred from published studies of human histones and partially confirmed by assignment experiments conducted here.^95, 100^ For H2A.Z, we were able to unambiguously assign several residues and guess the location of others based on characteristic chemical shifts and comparison between H2A, H2A.Z, and C-tail swap mutant H2A.Z^C-H2A^ NCP spectra (Fig. 8 and Fig. S16).

**Figure 8.**
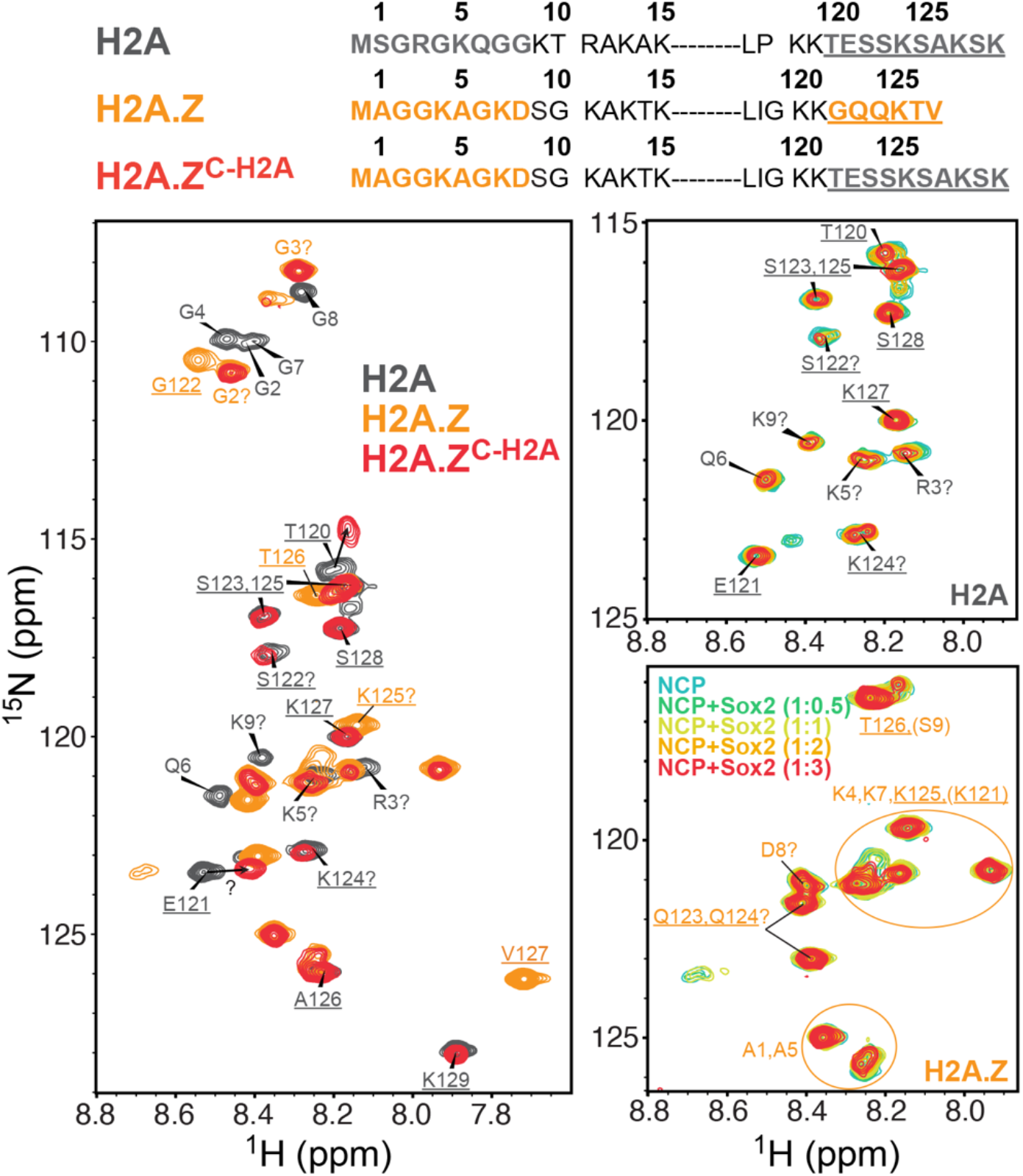
Sox2 perturbs the H2A C-terminal tail more than the H2A.Z C-terminal tail. Amino acid sequences (top) showing residues of the N- and C-terminal tails (bold), with the C-tail underlined. ^1^H-^15^N HSQC spectral overlay (bottom) of free NCP 62_F_(~20 μM) containing ^15^N/^13^C-labeled H2A (grey), H2A.Z (orange), or H2A.Z^C-H2A^ (red), shown on the left. Titrations of H2A and H2A.Z NCPs with increasing concentrations of Sox2 (0.5, 1, 2, and 3X), recorded at 37°C, shown on the right (see Fig. S16 for full and H2A.Z^C-H2A^ spectra). Peak assignments indicated by “?” represent our best guess based on published or predicted chemical shifts (BMRB).

Addition of Sox2 caused visible chemical shift changes and line broadening in multiple residues of the H2A C- and N-tail in canonical NCPs (Fig. 8 and Fig. S16). By contrast, the effect of Sox2 on the H2A.Z C-tail was minimal, suggesting that Sox2 binding at SHL+6 (NCP 62_F_) induces larger conformational changes in the longer H2A C-tail. Moreover, the destabilized H2A.Z^C-H2A^ mutant differed from that of the H2A C-tail (Fig. 8), with the E121 peak being either strongly perturbed (also T120) or completely line broadened due to chemical exchange. A similar E121 signature (i.e. large shift or broadening) was observed for E121 in an H2A-H2B dimer bound to DNA (Fig. S16B), implying an altered conformation and/or protonation state. Sox2 addition to H2A.Z^C-H2A^ NCP also produced smaller CSPs for the same residues affected in the H2A NCP (Fig. S16A). This data indicates that the H2A C-tail in the H2A.Z mutant adopts a distinct conformation than in canonical H2A, which is closer to the free H3 tail state and less responsive to Sox2. Given the greater unwrapping observed in H2A.Z^C-H2A^ (Fig. S14C), the reduced changes are likely due to weakened C-tail interactions with DNA and H3.

We also examined perturbations to the H4 N-tail upon Sox2 binding at SHL+2 (NCP 23R_N_) (Fig. S17). The H4 N-tail adopted similar conformations in H2A and H2A.Z NCPs, as indicated by their nearly identical spectra, and Sox2 addition produced negligible effects in both NCPs (Fig. S17). Thus, the enhanced Sox2 binding at SHL+2 in H2A.Z NCPs is unlikely to arise from changes in H4 N-tail dynamics or reduced competition for DNA. Instead, a destabilized H3 N-tail, coupled with increased DNA accessibility, may create more favorable energetics for Sox2 binding to H2A.Z nucleosomes.

### Sox2 globally perturbs nucleosomes and maintains unique H3 N-tail contacts with H2A.Z

Finally, to probe how Sox2 binding perturbs nucleosomes with atomic detail and how H2A.Z might facilitate these interactions, we performed three 1 μs MD simulations of the 601 NCP with Sox2 bound at SHL-6 (Fig. 9), based on a published cryo-EM structure.^65^ Sox2 binding induced global changes in nucleosome conformation, deforming both DNA and the histone core (Fig. 9A,B, and Fig. S6). Specifically, the two otherwise parallel DNA gyres were shifted laterally relative to one another, accompanied by a larger vertical compression in the case of H2A.Z (Fig. 9A,B, and Fig. S18). These changes suggest that Sox2 binding and DNA bending pulls and deforms one DNA gyre, leading to gyre misalignment and histone core distortion. Simultaneously, the H3 N-tails at SHL+7 (opposite side of Sox2) for both H2A and H2A.Z systems adopted extended conformations beyond SHL+5, partially filling the space created by the gyre misalignment (Fig. S6). This resulted in extensive new DNA contacts, likely compensating for loss of histone-DNA interactions and overall symmetry.

**Figure 9.**
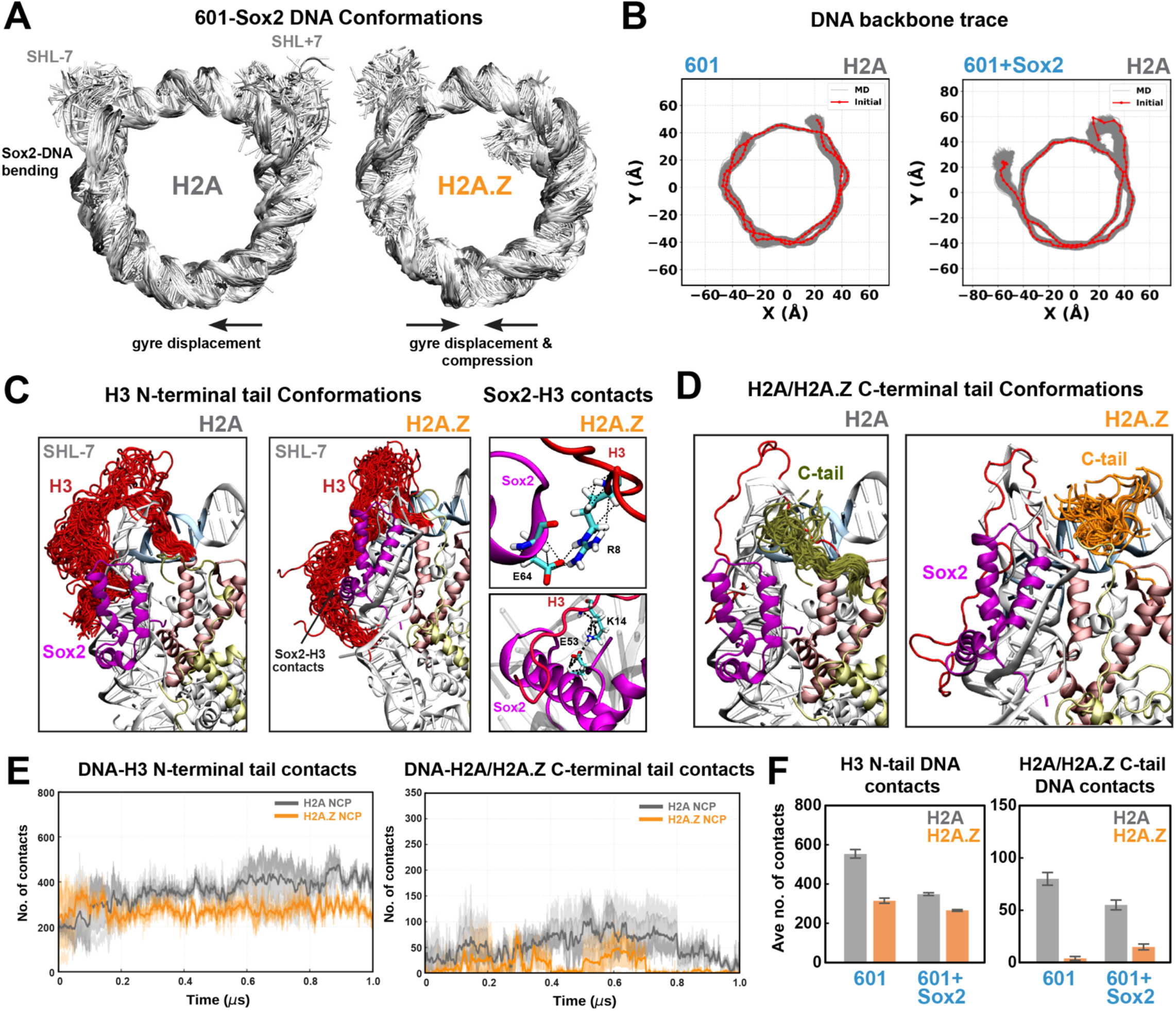
Sox2 binding redistributes histone-DNA contacts and enables unique Sox2-H3 tail interactions in the presence of H2A.Z. (A) Superimposed DNA conformations from MD simulations showing Sox2-induced DNA bending, gyre distortion, and increased DNA unwrapping. (B) DNA backbone traces for 601 NCP in the absence and presence of Sox2, showing significant distortion and gyre misalignment. (C) Superimposed H3 N-terminal tail conformations (red) depicting altered conformations with H2A.Z that facilitate unique Sox2-H3 tail interactions between specific Arg/Lys and Glu residues (right panel). (D) Superimposed conformations of H2A (olive) and H2A.Z (orange) C-terminal tails showing shifted ensembles for the H2A tail for the bound versus free NCP (see Fig. 4C). (E) Total contacts (4.5 Å cutoff) between DNA and the H3 N-tail (left) or the H2A/H2A.Z C-tail (right) as a function of time (averaged over 3 replicas), showing fewer contacts in the H2A.Z (orange) versus H2A (gray) system. (F) Average number of DNA contacts for the H3 N-tail (left) and H2A/H2A.Z C-tail (right) (near SHL-7) showing a large decrease with Sox2 binding and a larger effect for H2A versus H2A.Z.

We also observed that histone tail interactions with DNA, especially near the Sox2 binding site, were altered and reduced in the Sox2-NCP complexes (Fig. 9C,D, Figs S6, S19, and S20). The average H3 N-tail-DNA contacts decreased from roughly 550 to 300, while H2A C-tail-DNA contacts decreased from roughly 80 to 50 for canonical NCPs at the Sox2 site (Fig. 9E, F). The same trend was observed for H3 tail-DNA contacts on the opposite site (SHL+7) (Fig. S19). These results agree with NMR data showing Sox2-induced changes in both the H3 N-tail and H2A C-tail conformation (Figs 7 and 8), consistent with disruption of tail-DNA contacts. Importantly, fewer H3 tail-DNA contacts were on average disrupted by Sox2 in the H2A.Z system (Fig. 9D), and H2A.Z tail-DNA contacts were even increased, which could in part explain the increased Sox2 binding affinity.

Notably, we observed multiple Sox2-H3 N-tail interactions in H2A.Z NCPs that were absent in the canonical system (Fig. 9C and Fig. S21A,B). These were clustered in the Sox2 α3 helix and C-terminal tail and included hydrogen bonds between oppositely charged Arg/Lys and Glu residues – H3 R8 to Sox2 E64 (Fig. 9C), H3 K14 to Sox2 E53, and H3 E59 to Sox2 R74 (Fig. S21A,B). These unique contacts may arise from the altered DNA bending orientation in H2A.Z relative to canonical H2A (Fig. S21C). Our results suggest that Sox2-H3 interactions are more frequent with H2A.Z and, along with increased DNA accessibility and fewer disrupted histone-DNA contacts, could contribute to the enhanced Sox2-nucleosome association (Fig. 10).

**Figure 10.**
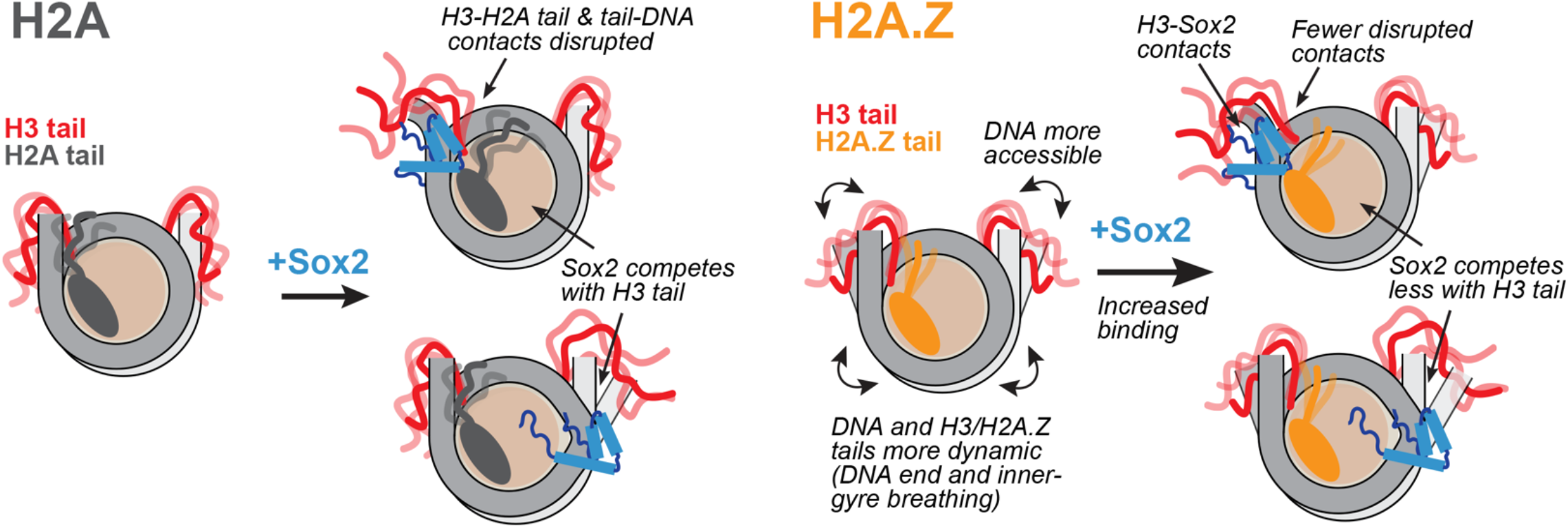
Proposed model for nucleosome destabilization and enhanced Sox2 binding by H2A.Z. Incorporation of H2A.Z increases DNA end and inner-gyre breathing and weakens key contacts between H3 N-tail, H2A C-tail, and nucleosomal DNA. This increases overall DNA mobility and accessibility and, together with new H3-Sox2 contacts, favors Sox2 binding at terminal or internal positions.

## Discussion

The histone variant H2A.Z can enhance chromatin accessibility and promote its interaction with chromatin factors to regulate gene expression. Here, we provide new molecular insights from experiments and computation that shed light on the role of H2A.Z in the interaction of pioneer factors Sox2 and Oct4 with nucleosomes. We find improved binding of Sox2 and Oct4 with H2A.Z at both end-positioned (SHL6) and internal (SHL2 for Sox2) nucleosome sites, which we correlate by NMR and simulation with more extensive DNA end and inner-gyre breathing and perturbed H3 N-terminal tail conformational dynamics. Simultaneously, we demonstrate that Sox2 competes with and displaces the H3 N-tail when associating at the same nucleosome positions. Sox2 and Oct4 binding could be favored by higher DNA accessibility and mobility and lower competition with H3 and, possibly, H2A tails for DNA. Thus, we show that pioneer factors can take advantage of H2A.Z-mediated changes in nucleosome conformation and dynamics to achieve more efficient engagement with chromatin. Other destabilizing histone tail disease-linked mutations, epigenetic modifications or proteolytic clipping could have similar and potentially greater impact on pioneer factor-chromatin interactions. Our findings support an emerging mechanism where histone tails and their epigenetics alter nucleosome dynamics and recognition to tune chromatin function.^36,37,42,45^

The destabilizing effect of H2A.Z on nucleosomes has been previously linked to its shorter and less positively charged C-terminal tail.^51-53^ Prior computational studies have shown that the H2A.Z C-tail forms fewer and more transient contacts with DNA, correlating with increased unwrapping, and that altered dynamics of the H2A.Z N-tail are associated with larger inter-gyre gapping.^53^ Moreover, deletion of the H2A C-tail also leads to increased histone exchange kinetics and nucleosome mobility *in vitro* and *in vivo*, implicating C-tail variants in similar functions.^46, 101^ While our results corroborate these findings, they provide new important insights into the H3 N-tail and its role in DNA mobility. We demonstrate that, with H2A.Z, the H3 N-tail is more dynamic and adopts distinct conformations with fewer DNA contacts. H3 N-tail conformations correlating with larger DNA breathing also bridge fewer contacts between adjacent DNA gyres, contributing to inter-gyre gapping, and rarely interact with SHL2. Prior studies have shown that high-salt-induced expansion of the inter-gyre distance similarly correlates with diminished H3 and other histone-DNA interactions,^38^ potentially mimicking the effects of H2A.Z. These molecular insights are validated by the striking agreement we observe between the H2A.Z-induced changes in H3 fluctuations and DNA contacts by simulation and changes in H3 N-tail dynamics obtained by NMR. Importantly, the conformational changes in H3 and nucleosomal DNA result in better DNA access at locations where we observe improved binding of Sox2 or Oct4 and where many other chromatin factors interact.^44, 45, 102^

Notably, our integrated NMR and MD data indicate a coupling between the H3 and H2A tail regions that is impaired with H2A.Z. MD simulations reveal a potential salt bridge between H3 R42 and H2A E121, also visualized in a 1.9 Å NCP structure,^99^ consistent with our NMR data. In the destabilized H2A.Z^C-H2A^ NCP mutant, which shows increased H3 N-tail mobility and DNA unwrapping, we observe extreme line broadening or a large change in E121 chemical shift, similar to the behavior of an isolated H2A-H2B dimer bound to DNA, and a smaller C-tail response to Sox2 binding. This altered NMR signature likely reflects disruption of the local 3_10_-helix^99^ and/or loss of C-tail contacts with H3 R42 or DNA. We propose that a potential H3 R42-H2A E121 interaction, missing in H2A.Z, could help stabilize H3 and H2A tails at the entry/exit DNA to limit unwrapping. H3 R42 can alternatively make hydrogen bonds with the DNA backbone.^37, 103^ Importantly, methylation at that site that abolishes H2A or DNA interactions has been shown to stimulate transcription *in vitro*.^104^ Moreover, mutations of H2A E121 (i.e. E121Q/K) are frequently encountered in human cancers and linked to disruption of higher-order chromatin structure.^48^ Thus, further targeted mutational studies for these residues could establish their role in nucleosome structure and dynamics.

We also demonstrate that the effect of H2A.Z is dependent on DNA sequence, with effects that are both significant and mechanistically distinct for the more labile Lin28B nucleosome. With H2A.Z, we find that Lin28B DNA is more dynamic or accessible around the dyad, consistent with the enhanced DNA unwrapping and flexibility observed in simulations. This could affect chromatin recognition by a variety of proteins targeting the nucleosome dyad region.^44,102^ Using experiment and computation, we further show that canonical Lin28B nucleosomes are inherently less stable and more dynamic than 601 constructs, with H3 N-tails making fewer DNA contacts, especially near SHL2. These pronounced increases in DNA and histone mobility in canonical contexts likely facilitate Sox2 binding at Lin28B sites that diverge from its high-affinity motif. Pre-existing dynamics may also explain the smaller H2A.Z effect on the apparent Sox2 binding affinity. Consistent with this interpretation, we previously showed,^66^ and further validated here, that mutations that disrupt DNA contacts and increase DNA mobility at the nucleosome edge (SHL6, H3 R49A) or internally (SHL2, H3 R83A) also enhance Sox2 binding. In Lin28B, Sox2 appears to remodel nucleosome structure globally by inducing DNA deformation and possibly sliding. This is in agreement with studies showing that Sox2 and Oct4 association entail substantial DNA distortion or repositioning^32,64,65,70,105^ and can evict linker histone H1,^105^ potentially altering local and higher-order chromatin packing. Moreover, H1 and H2A.Z seem to act antagonistically as the binding of H1 to nucleosomes is inhibited by the H2A.Z C-tail^106^ and the activity of H1 in yeast rDNA can be counteracted by H2A.Z.^107^ Additional studies will be required to define the impact of H2A.Z in multi-nucleosome assemblies and its interplay with histone H1. Together, our results support a model in which labile or variant genomic nucleosomes, acting in concert with epigenetic modifications, facilitate pioneer-factor engagement and promote structural transitions that yield more accessible and remodeling-competent chromatin.

Finally, we establish that Sox2 can globally perturb DNA and histones in both canonical and variant nucleosomes, providing new insights into how multiple histone tails are affected. We find that Sox2 can displace the N-tail of H3 when associating with modified 601* and Lin28B nucleosomes at both internal and end positions (SHL2, 5, 6; possibly SHL3 in Lin28B), with a larger effect in the presence of destabilizing H2A.Z and H3 mutant histones. Moreover, Sox2 binding can visibly perturb the C- and N-tail of H2A (SHL6) as well as the N-tail of H2B (SHL5, to be published elsewhere). While the H4 N-tail was not significantly impacted by Sox2 binding at SHL2, that might not be the case when other nucleosome positions are targeted. Collectively, our data illustrates that Sox2 can directly and allosterically disrupt the conformation and dynamics of nucleosomes, including multiple histone tails known to regulate DNA dynamics, inter-nucleosome contacts, and nucleosome sliding.^40,41,46,108-110^ Perturbations to histone tails by pioneer factors could have a major impact on chromatin structure, recognition, and epigenetic changes. Future research efforts should explore the role of these and other pioneer factors on DNA and histone tail conformation and dynamics, and how that might propagate to higher-order chromatin structure or tune the activity of chromatin readers and writers.

## Acknowledgements

We thank Prof. Gregory Bowman (JHU) for kindly providing materials and access to instrumentation and Prof. Carl Wu (JHU) for the H2A.Z plasmid. We acknowledge Fabiana Malaga Gadea for preparing protein and nucleosome samples. We are grateful to Dr. Ananya Majumdar, director of the Biomolecular NMR center (JHU), for assistance with NMR data collection and to the Integrated Imaging Center (JHU) staff for bioimager training and support. Anton 3 computer time was provided by the Pittsburgh Supercomputing Center (PSC) through Grant 1R24GM154042 from the National Institutes of Health. The Anton 3 machine at PSC is made available by D. E. Shaw Research. We thank the CUNY-HPCC at the College of Staten Island for computer time. R.P. would like to acknowledge Dr. Abhik Ghosh Moulick for helpful discussions. This research was supported by grants R01GM147642 awarded to E.N.N. and 1R15GM146228 awarded to S.M.L. R. P. is grateful for support from The Rosemary O’Halloran Scholarship.

## Author Contributions

Conceptualization: E.N.N., H.K.M., S.M.L. Methodology: E.N.N., H.K.M., R.P., S.M.L. Investigation: H.K.M., R.P., E.N.N., S.K.F, S.M.L. Formal analysis: H.K.M., R.P., E.N.N., S.M.L. Writing – original draft: E.N.N., H.K.M., R.P., S.M.L. Writing – review & editing: E.N.N., H.K.M., S.K.F., R.P., S.M.L. Visualization: H.K.M., R.P, E.N.N. Supervision: E.N.N., S.M.L. Funding acquisition: E.N.N., S.M.L. Resources: E.N.N., S.M.L.

## Supplementary Data

Supplementary Data are available online.

## Data Availability

Analysis codes are available on GitHub Repository https://github.com/CUNY-CSI-Loverde-Laboratory/H2A.Z_Variant_MD_Analysis. Code and simulation trajectories are available on Zenodo https://doi.org/10.5281/zenodo.19454347. NMR chemical shift perturbation data for histone tails have been deposited to the Biological Magnetic Resonance Bank (BMRB IDs: 52922-52935).

