## Supplemental Data for "H2A.Z facilitates Sox2-nucleosome interaction by promoting DNA and histone H3 tail mobility"

### Supplementary Methods

#### Additional Simulation Details

All systems were initially minimized to reduce unfavorable stress by using conjugate gradient and steepest descent gradient for 40 ps. Heating was performed by increasing the temperature of the system to 310 K under *NVT* conditions. Then, the systems were equilibrated to 100 ns under *NPT* conditions. The Langevin dynamics method,[1] with the collision frequency (friction constant of 1 $ps^{-1}$) was used to control the temperature of the system. The pressure of the system was controlled by the Berendsen[2] barostat. The simulation continued for 1 μs under *NPT* conditions with a 2 fs each timestep. Trajectories were stored after every 200 ps for analysis. All simulations used the SHAKE[3] algorithm to constrain hydrogen bonds. The Lennard-Jones cutoff value for nonbonded interactions was 12 Å, and electrostatic interactions were treated with the Particle Mesh Ewald (PME)[4] method using full periodic boundary conditions. The number of total molecules and ions are shown in Table S4.

#### Molecular Dynamics Data Analysis

Molecular dynamics (MD) trajectories for H2A and H2A.Z systems were first analyzed for root mean square fluctuations (RMSF) of the Cα atoms of the histone H3 N-terminal tails (1-43 residues) from the structure of the nucleosome. The entire trajectory of 1 μs after the 100 ns of equilibration was used. The RMSF data of both systems was plotted using Python. The number of contacts between DNA and histone H3 N-terminal tails was analyzed for 1 μs. The interactions were defined as between two heavy atoms of DNA and histone H3 tail residues with a distance cutoff value of 4.5 Å. The contact maps are obtained using MDAnalysis[5] based on the same distance cutoff. The number of contacts was analyzed for H2A and H2A.Z C-terminal tails. The secondary structure of the H3 N-terminal tails for both H2A and H2A.Z systems was determined using the AmberTools21 secstruct tool, which uses the DSSP[6] algorithm. This algorithm is based on hydrogen bonding patterns in the protein backbone amide (N-H) and carbonyl (C=O) positions. The algorithm provides secondary structure including α-helix, $3_{10}$-helices, turns, β-sheets, coils, or no structure. The gapping distance is quantitatively characterized by measuring the distance between the two DNA segments in SHL-4 and SHL+4, similar to *Panchenko et al.* The gap distance is measured as the center-of-mass separation between the base-pair groups of SHL-4 and SHL+4. For each frame of the trajectory, the center of mass of the SHL-4 and SHL+4 residues is computed to evaluate the distance. The reported distance is averaged over the entire 1 μs trajectory for H2A and H2A.Z systems. In addition, DNA backbone traces were generated by calculating the center of mass for each trajectory frame, and these positions were projected onto the X-Y plane, centered on the reference structure. The 2D plot of the backbone trace is visualized for backbone motion. All MD frames were overlaid and compared against the initial structure to capture the DNA fluctuations.

**Table S1.** Nucleotide sequences for Widom 601, modified 601* and Lin28B DNA constructs used for nucleosome preparation (the dyad (0$^{th}$) position is underlined). Cartoon showing the position of Sox2-Oct4 binding sites on the 601* sequence relative to nucleosome SHLs.

**Widom 601**
(601) 5'-TG GAGAATCCCG GTGCCGAGGC CGCTCAATTG GTCGTAGACA GCTCTAGCAC CGCTTAAACG CACGTACGCG C TGTCCCCCGC GTTTTAACCG CCAAGGGGAT TACTCCCTAG TCTCCAGGCA CGTGTCAGAT ATATACATCC TG-3'

**Modified Widom 601***
(FAM labeled strand; mutated region from Widom 601 is highlighted in gray)
(601*) 5'-TG GAGAATCCCG GTGCCGAGGC CGCTCAATTG GTCGTAGACA GCTCTAGCAC CGCTTAAACG CACGTACGCG C TGTCCCCCGC GTTTTAACCG CCAAGGGGAT TACTCCCTAG TCTCCAGGCA CGTGTGGTTA CGAGGCTATC GT-3'

*Oct4 POU$_S$ site shown in red. Oct4 POU$_{HD}$ site shown in green. Sox2 HMG site shown in blue.*

**601* mFgf4**
(m62$_F$) 5'-TG GAGACTTTGT TTGGATGCTA ATCTCAATTG GTCGTAGACA GCTCTAGCAC CGCTTAAACG CACGTACGCG C TGTCCCCCGC GTTTTAACCG CCAAGGGGAT TACTCCCTAG TCTCCAGGCA CGTGTGGTTA CGAGGCTATC GT-3'
(23R$_F$) 5'-TG GAGAATCCCG GTGCCGAGGC CGCTCAATTG GTCGTAGACA GCTCTAGCAC CGCTTAAACG CACGTACGCG C TGTCCCCCGC GTTTTAACCT TTGTTTGGAT GCTAATCTAG TCTCCAGGCA CGTGTGGTTA CGAGGCTATC GT-3'
(54R$_F$) 5'-TG GAGAATCCCG GTGCCGAGGC CGCTCAATTG GTCGTAGACA GCTCTAGCAC CGCTTAAACG CACGTACGCG C TGTCCCCCGC GTTTTAACCG CCAAGGGGAT TACTCCCTAG TCTCCAGGCC TTTGTTTGGA TGCTAATATC GT-3'
(62$_F$) 5'-TG GAGAATCCCG GTGCCGAGGC CGCTCAATTG GTCGTAGACA GCTCTAGCAC CGCTTAAACG CACGTACGCG C TGTCCCCCGC GTTTTAACCG CCAAGGGGAT TACTCCCTAG TCTCCAGGAT TAGCATCCAA ACAAAGTATC GT-3'

**601* mNanog**
(m62$_N$) 5'-TG GAGACATTGT AATGCAAAAC CGCTCAATTG GTCGTAGACA GCTCTAGCAC CGCTTAAACG CACGTACGCG C TGTCCCCCGC GTTTTAACCG CCAAGGGGAT TACTCCCTAG TCTCCAGGCA CGTGTGGTTA CGAGGCTATC GT-3'
(23R$_N$) 5'-TG GAGAATCCCG GTGCCGAGGC CGCTCAATTG GTCGTAGACA GCTCTAGCAC CGCTTAAACG CACGTACGCG C TGTCCCCCGC GTTTTAACCA TTGTAATGCA AAATCCCTAG TCTCCAGGCA CGTGTGGTTA CGAGGCTATC GT-3'
(54R$_N$) 5'-TG GAGAATCCCG GTGCCGAGGC CGCTCAATTG GTCGTAGACA GCTCTAGCAC CGCTTAAACG CACGTACGCG C TGTCCCCCGC GTTTTAACCG CCAAGGGGAT TACTCCCTAG TCTCCAGGCC ATTGTAATGC AAAAGCTATC GT-3'
(62$_N$) 5'-TG GAGAATCCCG GTGCCGAGGC CGCTCAATTG GTCGTAGACA GCTCTAGCAC CGCTTAAACG CACGTACGCG C TGTCCCCCGC GTTTTAACCG CCAAGGGGAT TACTCCCTAG TCTCCAGGCA CTTTTGCATT ACAATGTATC GT-3'

**Lin28B**
(Cy3 labeled strand only)
(Lin28B) 5'-ACA TATCCTCAGT GGTGAGTATT AACATGGAAC TTACTCCAAC AATACAGATG CTGAATAAAT GTAGTCTAAG T GAAGGAAGAA GGAAAGGTGG GAGCTGCCAT CACTCAGAAT TGTCCAGCAG GGATTGTGCA AGCTTGTGAA TAA-3'

**DNA40 (mFgf4)**
5'-ACGATACTTTGTTTGGATGCTAATCCTGGAGACTAGGGAG-3'

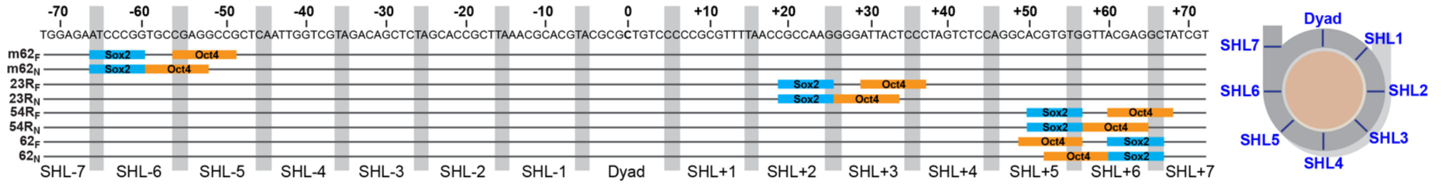

**Table S2.** Amino acid sequences for protein constructs used in this study. Mutated regions from the canonical sequence are shown in red and underlined.

| ***X. laevis* H2A** |
|---|
| MSGRGKQGGK TRAKAKTRSS RAGLQFPVGR VHRLLRKGNY AERVGAGAPV YLAAVLEYL TAEILELAGNA ARDNKKTRII PRHLQLAVRN DEELNKLLGR VTIAQGGVLP NIQSVLLPKK TESSKSAK SK<br><br>H2A$^{\text{C-H2A.Z}}$:<br>MSGRGKQGGK TRAKAKTRSS RAGLQFPVGR VHRLLRKGNY AERVGAGAPV YLAAVLEYL TAEILELAGNA ARDNKKTRII PRHLQLAVRN DEELNKLLGR VTIAQGGVLP NIQSVLLPKK GQQKTV |
| ***H. sapiens* H2A.Z-1** |
| MAGGKAGKDS GKAKTKAVSR SQRAGLQFPV GRIHRHLKSR TTSHGRVGAT AAVYSAAILE YLTAEVLELA GNASKDLKVK RITPRHLQLA IRGDEELDSL IKATIAGGGV IPHIHKSLIG KKGQQKTV<br><br>H2A.Z$^{\text{N-H2A}}$:<br>MSGRGKQGGK TRAKAKTRSQ RAGLQFPVGR IHRHLKTRTT SHGRVGATAA VYSAAILEYL TAEVLELAGN ASKDLKVKRI TPRHLQLAIR GDEELDSLIK ATIAGGGVIP HIHKSLIGKK GQQKTV<br><br>H2A.Z$^{\text{L-H2A}}$:<br>MAGGKAGKDS GKAKTKAVSR SQRAGLQFPV GRIHRHLRKG NYAERVGAGA AVYSAAILEY LTAEVLELAG NASKDLKVKR ITPRHLQLAI RGDEELDSLI KATIAGGGVI PHIHKSLIGK KGQQKTV<br><br>H2A.Z$^{\text{C-H2A}}$:<br>MAGGKAGKDS GKAKTKAVSR SQRAGLQFPV GRIHRHLKSR TTSHGRVGAT AAVYSAAILE YLTAEVLELA GNASKDLKVK RITPRHLQLA IRGDEELDSL IKATIAGGGV IPHIHKSLIG KKTESSKSAK SK |
| ***X. laevis* H2B** |
| MAKSAPAPKK GSKKAVTKTQ KKDGKKRRKT RKESYAIYVY KVLKQVHPDT GISSKAMSIM NSFVNDVFER IAGEASRLAH YNKRSTITSR EIQTAVRLLL PGELAKHAVS EGTKAVTKYT SAK |
| ***X. laevis* H3** |
| MARTKQTARK STGGKAPRKQ LATKAARKSA PATGGVKKPH RYRPGTVALR EIRRYQKSTE LLIRKLPFQR LVREIAQDFK TDLRFQSSAV MALQEASEAY LVALFEDTNL CAIHAKRVTI MPKDIQLARR IRGERA<br><br>H3 R49A:<br>MARTKQTARK STGGKAPRKQ LATKAARKSA PATGGVKKPH RYRPGTVALA EIRRYQKSTE LLIRKLPFQR LVREIAQDFK TDLRFQSSAV MALQEASEAY LVALFEDTNL CAIHAKRVTI MPKDIQLARR IRGERA<br><br>H3 R83A:<br>MARTKQTARK STGGKAPRKQ LATKAARKSA PATGGVKKPH RYRPGTVALR EIRRYQKSTE LLIRKLPFQR LVREIAQDFK TDLAFQSSAV MALQEASEAY LVALFEDTNL CAIHAKRVTI MPKDIQLARR IRGERA |
| ***X. laevis* H4:** |
| MSGRGKGGKG LGKGGAKRHR KVLRDNIQGI TKPAIRRLAR RGGVKRISGL IYEETRGVLK VFLENVIRDA VTYTEHAKRK TVTAMDVVYA LKRQGRTLYG FGG |

**Other Proteins:**

| ***H. Sapiens* Sox2 DBD (HMG)** |
|---|
| GPDRVKRPMN AFMVWSRGQR RKMAQENPKM HNSEISKRLG AEWKLLSETE KRPFIDEAKR LRALHMKEHP DYKYRPRRKT KT |
| ***H. Sapiens* Oct4 DBD (POU$_{\text{S}}$-POU$_{\text{HD}}$)** |
| GPEESQDIK ALQKELEQFA KLLKQKRITL GYTQADVGLT LGVLFGKVFS QTTICRFEAL QLSFKNMCKL RPLLQKWVEE ADNNENLQEI CKAETLVQAR KRKRTSIENR VRGNLENLFL QCPKPTLQQI SHIAQQLGLE KDVVRVWFCN RRQKGKRSSS DYAQRE<br><br>CtoS mutant:<br>GPEESQDIK ALQKELEQFA KLLKQKRITL GYTQADVGLT LGVLFGKVFS QTTISRFEAL QLSFKNMSKL RPLLQKWVEE ADNNENLQEI SKAETLVQAR KRKRTSIENR VRGNLENLFL QSPKPTLQQI SHIAQQLGLE KDVVRVWFSN RRQKGKRSSS DYAQRE |

**Table S3.** ExoIII digestion rates obtained from monoexponential fits of full-length DNA (SHL±7).

| | | **k (min$^{-1}$)** | **k (min$^{-1}$)** | **k (min$^{-1}$)** |
|---|---|---|---|---|
| **NCP construct** | | ***H2A*** | ***H2A.Z*** | ***H3 R49A*** |
| $62_F$ | SHL-7 | 0.050 ± 0.011 | 0.101 ± 0.026 | 0.198 ± 0.014 |
| | SHL+7 | 0.066 ± 0.018 | 0.082 ± 0.023 | 0.203 ± 0.053 |
| $62_N$ | SHL-7 | 0.051 ± 0.011 | 0.070 ± 0.017 | n/d* |
| | SHL+7 | 0.091 ± 0.020 | 0.109 ± 0.027 | n/d* |
| $m62_F$ | SHL-7 | 0.302 ± 0.084 | 0.244 ± 0.077 | 0.945 ± 0.162 |
| | SHL+7 | 0.060 ± 0.007 | 0.053 ± 0.011 | 0.198 ± 0.079 |
| $m62_N$ | SHL-7 | 0.196 ± 0.009 | 0.208 ± 0.019 | n/d* |
| | SHL+7 | 0.055 ± 0.024 | 0.082 ± 0.021 | n/d* |
| $54R_F$ | SHL-7 | 0.044 ± 0.012 | 0.075 ± 0.020 | n/a† |
| | SHL+7 | 0.031 ± 0.018 | 0.047 ± 0.012 | n/a† |
| $54R_N$ | SHL-7 | 0.056 ± 0.006 | 0.095 ± 0.008 | n/a† |
| | SHL+7 | 0.095 ± 0.018 | 0.091 ± 0.027 | n/a† |
| $23R_N$ | SHL-7 | 0.065 ± 0.012 | 0.079 ± 0.009 | n/a† |
| | SHL+7 | 0.113 ± 0.009 | 0.080 ± 0.013 | n/a† |
| $23R_F$ | SHL-7 | 0.043 ± 0.011 | 0.053 ± 0.011 | n/a† |
| | SHL+7 | 0.025 ± 0.004 | 0.033 ± 0.003 | n/a† |
| Lin28B | SHL-7 | 0.097 ± 0.040 | 0.075 ± 0.024 | n/a† |
| | SHL+7 | 0.043 ± 0.019 | 0.033 ± 0.026 | n/a† |

*n/d = could not be determined
†n/a = not available/measured

**Table S4.** Summary of calculated FRET ratios and statistics for different NCP constructs.

| NCP | Histone + Ligand | Mean | s.d. (sample) | s.d. (mean) | s.d. (total) | Reduced $\chi^2$ | *p*-value (H2A)[a] | *p*-value (H2A.Z)[a] | n |
|---|---|---|---|---|---|---|---|---|---|
| 601* | H2A | 0.2445 | 0.0082 | 0.0027 | 0.0037 | 0.9304 | | | 10 |
| 54R$_F$ | H2A | 0.2650 | 0.0049 | 0.0033 | 0.0039 | 0.3756 | | | 6 |
| | H2A.Z | 0.2028 | 0.0113 | 0.0036 | 0.0054 | 1.4286 | 6.15E-07 | | 8 |
| 54R$_N$ | H2A | 0.2083 | 0.0039 | 0.0047 | 0.0051 | 0.1699 | | | 6 |
| | H2A.Z | 0.1587 | 0.0083 | 0.0043 | 0.0055 | 0.6315 | 2.51E-06 | | 6 |
| 62$_F$ | H2A | 0.2297 | 0.0014 | 0.0040 | 0.0040 | 0.0259 | | | 7 |
| | H2A.Z | 0.2100 | 0.0076 | 0.0035 | 0.0046 | 0.6937 | 3.79E-04 | | 7 |
| | H3 R49A | 0.1036 | 0.0125 | 0.0036 | 0.0052 | 1.0729 | 5.17E-12 | 1.59E-13 | 11 |
| | H2A.Z$^{C-H2A}$ | 0.2008 | 0.0048 | 0.0034 | 0.0038 | 0.2433 | 8.91E-08 | 2.06E-02 | 8 |
| | H2A.Z$^{L1-H2A}$ | 0.2119 | 0.0050 | 0.0031 | 0.0035 | 0.2386 | 1.50E-06 | **0.588** | 9 |
| | H2A.Z$^{N-H2A}$ | 0.2108 | 0.0061 | 0.0035 | 0.0042 | 0.4192 | 1.05E-04 | **0.845** | 7 |
| 62$_N$ | H2A | 0.1951 | 0.0086 | 0.0031 | 0.0041 | 0.8280 | | | 10 |
| | H2A.Z | 0.1756 | 0.0080 | 0.0032 | 0.0041 | 0.6504 | 5.76E-05 | | 10 |
| | H3 R49A | 0.0769 | 0.0035 | 0.0064 | 0.0066 | 0.0772 | 1.57E-13 | 1.11E-12 | 4 |
| | H2A.Z$^{C-H2A}$ | 0.1573 | 0.0041 | 0.0053 | 0.0057 | 0.1431 | 2.00E-07 | 1.58E-04 | 6 |
| m62$_F$ | H2A | 0.1645 | 0.0081 | 0.0033 | 0.0042 | 0.5929 | | | 10 |
| | H2A.Z | 0.1732 | 0.0107 | 0.0034 | 0.0049 | 1.0872 | **0.065** | | 9 |
| | H3 R49A | 0.0733 | 0.0052 | 0.0046 | 0.0053 | 0.1661 | 7.42E-15 | 1.17E-11 | 4 |
| | H2A.Z$^{C-H2A}$ | 0.1370 | 0.0148 | 0.0056 | 0.0093 | 1.7948 | 2.74E-02 | 9.05E-03 | 6 |
| | H2A.Z$^{L1-H2A}$ | 0.1360 | 0.0019 | 0.0046 | 0.0046 | 0.0284 | 5.67E-07 | 5.59E-06 | 6 |
| | H2A.Z$^{N-H2A}$ | 0.1776 | 0.0067 | 0.0051 | 0.0058 | 0.4361 | **0.0179** | **0.393** | 6 |
| m62$_N$ | H2A | 0.1977 | 0.0086 | 0.0023 | 0.0030 | 0.8411 | | | 18 |
| | H2A.Z | 0.1794 | 0.0078 | 0.0023 | 0.0029 | 0.5899 | 6.17E-08 | | 18 |
| | H3 R49A | 0.0757 | 0.0051 | 0.0064 | 0.0069 | 0.1587 | 1.01E-09 | 2.90E-08 | 4 |
| | H2A.Z$^{C-H2A}$ | 0.1484 | 0.0020 | 0.0054 | 0.0055 | 0.0340 | 2.86E-15 | 1.53E-12 | 6 |
| | H2A.Z$^{L1-H2A}$ | 0.1444 | 0.0086 | 0.0039 | 0.0049 | 0.6272 | 1.22E-09 | 3.96E-07 | 8 |
| | H2A.Z$^{N-H2A}$ | 0.1663 | 0.0162 | 0.0043 | 0.0079 | 2.3974 | 0.00394 | 0.105 | 6 |
| | H2A$^{C-H2A.Z}$ | 0.1558 | 0.0097 | 0.0034 | 0.0046 | 0.8508 | 2.25E-09 | 6.87E-06 | 10 |
| 23R$_F$ | H2A | 0.2193 | 0.0034 | 0.0037 | 0.0040 | 0.1388 | | | 6 |
| | H2A.Z | 0.2116 | 0.0069 | 0.0031 | 0.0039 | 0.5668 | 0.0138 | | 9 |
| 23R$_N$ | H2A | 0.1988 | 0.0055 | 0.0034 | 0.0038 | 0.3224 | | | 10 |
| | H2A.Z | 0.1881 | 0.0123 | 0.0033 | 0.0051 | 1.5308 | 0.0368 | | 10 |
| | H3 R49A | 0.0972 | 0.0136 | 0.0061 | 0.0092 | 1.2681 | 0.0001 | 0.0001 | 4 |
| | H3 R84A | 0.1933 | 0.0051 | 0.0049 | 0.0055 | 0.2637 | **0.129** | **0.303** | 4 |
| Lin28B | H2A | 0.2017 | 0.0045 | 0.0034 | 0.0037 | 0.2191 | | | 8 |
| | H2A.Z | 0.1776 | 0.0051 | 0.0036 | 0.0040 | 0.2594 | 1.08E-07 | | 8 |
| | H2A$^{C-H2A.Z}$ | 0.1492 | 0.0029 | 0.0038 | 0.0040 | 0.0710 | 3.01E-12 | 2.90E-08 | 8 |
| **NCP** | **-/+ Sox2/Oct4[b]** | | | | | | **(free NCP)** | **(+Sox2)** | |
| 62$_N$ | H2A | 0.1951 | 0.0086 | 0.0031 | 0.0041 | 0.8280 | | | 10 |
| | H2A+Sox2 | 0.1759 | 0.0082 | 0.0032 | 0.0000 | 0.6555 | 7.73E-05 | | 8 |
| | H2A+Sox2/Oct4 | 0.0860 | 0.0287 | 0.0044 | 0.0258 | 5.4233 | 6.26E-06 | 2.75E-05 | 9 |
| | H2A.Z | 0.1756 | 0.0080 | 0.0032 | 0.0041 | 0.6504 | 5.76E-05 | | 10 |
| | H2A.Z+Sox2 | 0.1589 | 0.0172 | 0.0038 | 0.0135 | 2.6566 | 0.0303 | | 8 |
| | H2A.Z+Sox2/Oct4 | 0.0616 | 0.0164 | 0.0054 | 0.0097 | 1.5737 | 6.61E-06 | 2.98E-07 | 8 |
| m62$_N$ | H2A | 0.1977 | 0.0086 | 0.0023 | 0.0030 | 0.8411 | | | 18 |
| | H2A+Sox2 | 0.1788 | 0.0104 | 0.0029 | 0.0042 | 1.0373 | 2.41E-04 | | 12 |
| | H2A+Sox2/Oct4 | 0.0865 | 0.0280 | 0.0026 | 0.0085 | 4.9113 | 9.83E-09 | 4.15E-08 | 12 |
| | H2A.Z | 0.1794 | 0.0078 | 0.0023 | 0.0029 | 0.5899 | 6.17E-08 | | 18 |
| | H2A.Z+Sox2 | 0.1552 | 0.0091 | 0.0027 | 0.0038 | 0.7094 | 1.66E-09 | | 12 |
| | H2A.Z+Sox2/Oct4 | 0.0706 | 0.0068 | 0.0053 | 0.0057 | 0.2739 | 3.59E-11 | 1.45E-11 | 10 |

[a]*p*-value is reported relative to the same NCP containing either H2A or H2A.Z; bold p-values (> 0.05) indicate the result is not statistically significant at the 95% confidence level.
[b]FRET measurements in the presence or absence of 10X Sox2 and/or 10X Oct4. Free NCP values are the same as above.

**Table S5.** Summary of NCP systems MD simulation setup box sizes and atoms/ions.

| **Structures** | **Widom 601** | | **Lin28B** | | **SOX2** | |
|---|---|---|---|---|---|---|
| **Systems** | **H2A** | **H2A.Z** | **H2A** | **H2A.Z** | **H2A** | **H2A.Z** |
| **Box Size (Å$^3$)** | 171x124x185 | 171x127x185 | 142x227x241 | 137x227x241 | 196x142x202 | 191x142x202 |
| **No. of atoms** | 449702 | 459742 | 874536 | 838568 | 687086 | 686856 |
| **No. of Water Molecules** | 105978 | 108512 | 211891 | 202946 | 219716 | 219698 |
| **No. of $Na^+$ ions** | 484 | 496 | 857 | 836 | 763 | 769 |
| **No. of $Cl^-$ ions** | 360 | 366 | 707 | 680 | 624 | 630 |
| **NaCl Concentration (mM)** | 150 | 150 | 150 | 150 | 150 | 150 |

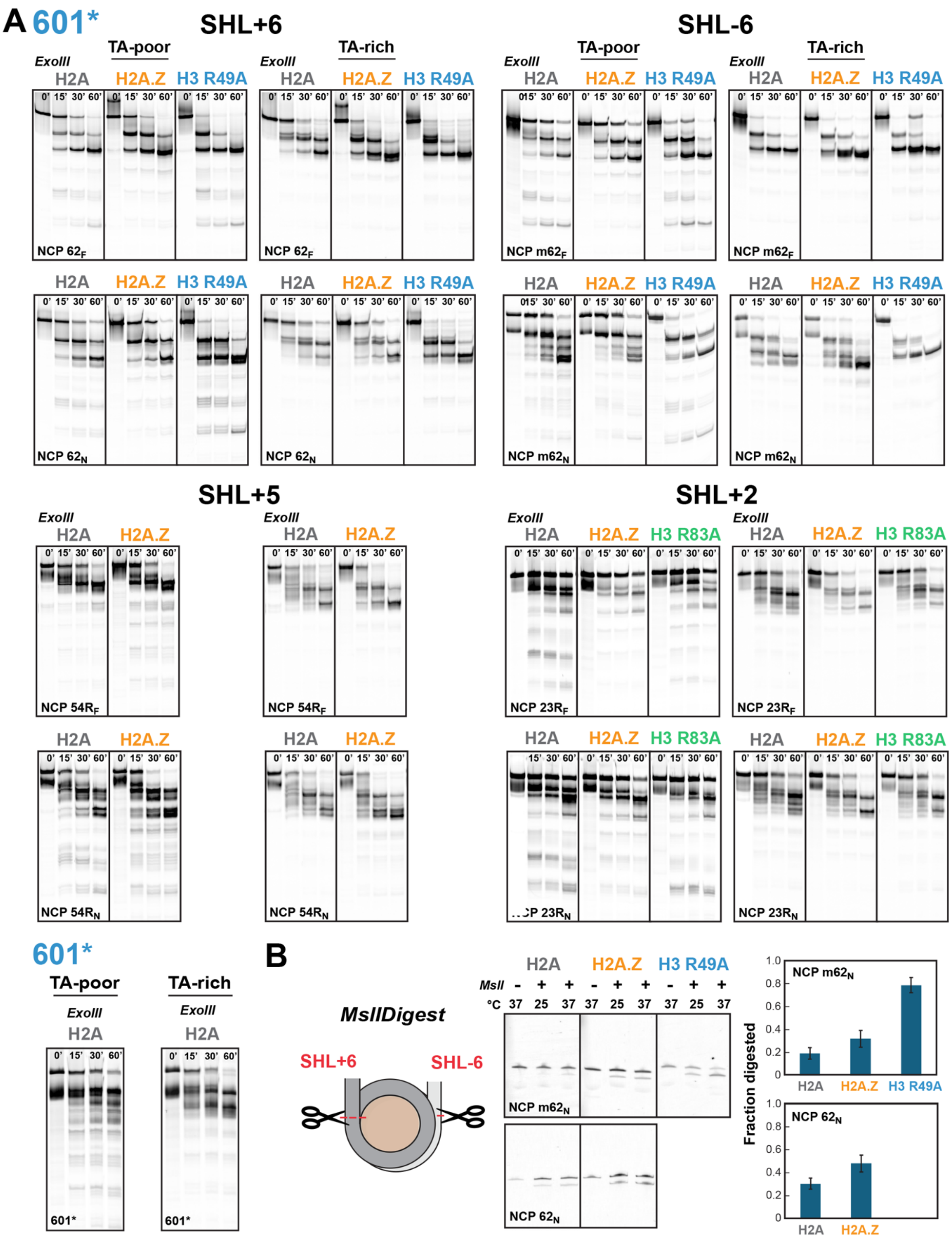


**Figure S1.** H2A.Z promotes DNA unwrapping in 601* nucleosomes. (A) Exonuclease III digestion time course for canonical H2A, H2A.Z, or H3 R49A 601* nucleosomes containing a Sox2-Oct4 motif at SHL+6 (NCP $62_F$, $62_N$), SHL-6 (NCP $m62_F$, $m62_N$), SHL+5 (NCP $54R_F$, $54R_N$), SHL+2 (NCP $23R_F$,$23R_N$) or no motif (601*). DNA digestion was monitored on denaturing gels using Cy3 (left, TA-rich side) or FAM (right, TA-poor side) fluorescence. More extensive digestion, consistent with increased DNA unwrapping, is typically observed for H2A.Z versus H2A near the nucleosome edge. The H3 R49A control displays strong unwrapping on both sides. On the TA-poor side, internal digestion (~SHL4-2) is increased for H2A, H3 R49A, and H3 R83A, but suppressed with H2A.Z. (B) MslI digestion assay monitored by denaturing gel electrophoresis for nucleosomes containing the Sox2-Oct4 motif at either SHL+6 (NCP $62_N$) or SHL-6 (NCP $m62_N$). Quantified fraction of digested DNA at 37°C (mean ± s.d.) shows cutting is enhanced with H2A.Z and further elevated with H3 R49A.

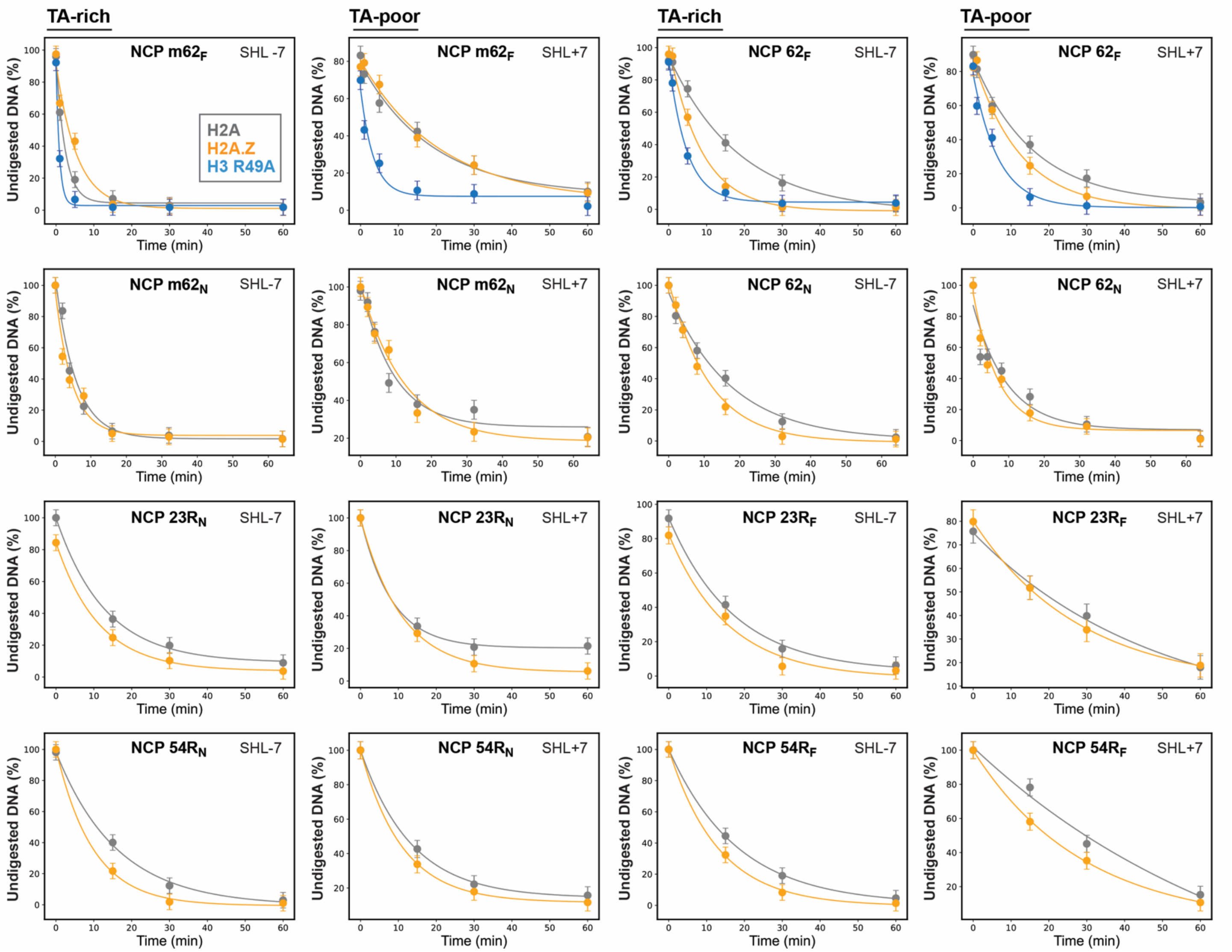


**Figure S2.** Representative plots of Exonuclease III digestion time dependence for 601* nucleosomes. Digestion profiles for H2A (gray), H2A.Z (orange), or H3 R49A (blue) 601* NCPs were quantified as DNA decay at SHL+7 (TA-poor side) and SHL-7 (TA-rich side). Average rates were determined by monoexponential fitting (solid lines) for 3-4 independent measurements (see Table S3). With some exceptions, faster digestion is generally observed with H2A.Z versus H2A in 601* NCPs.

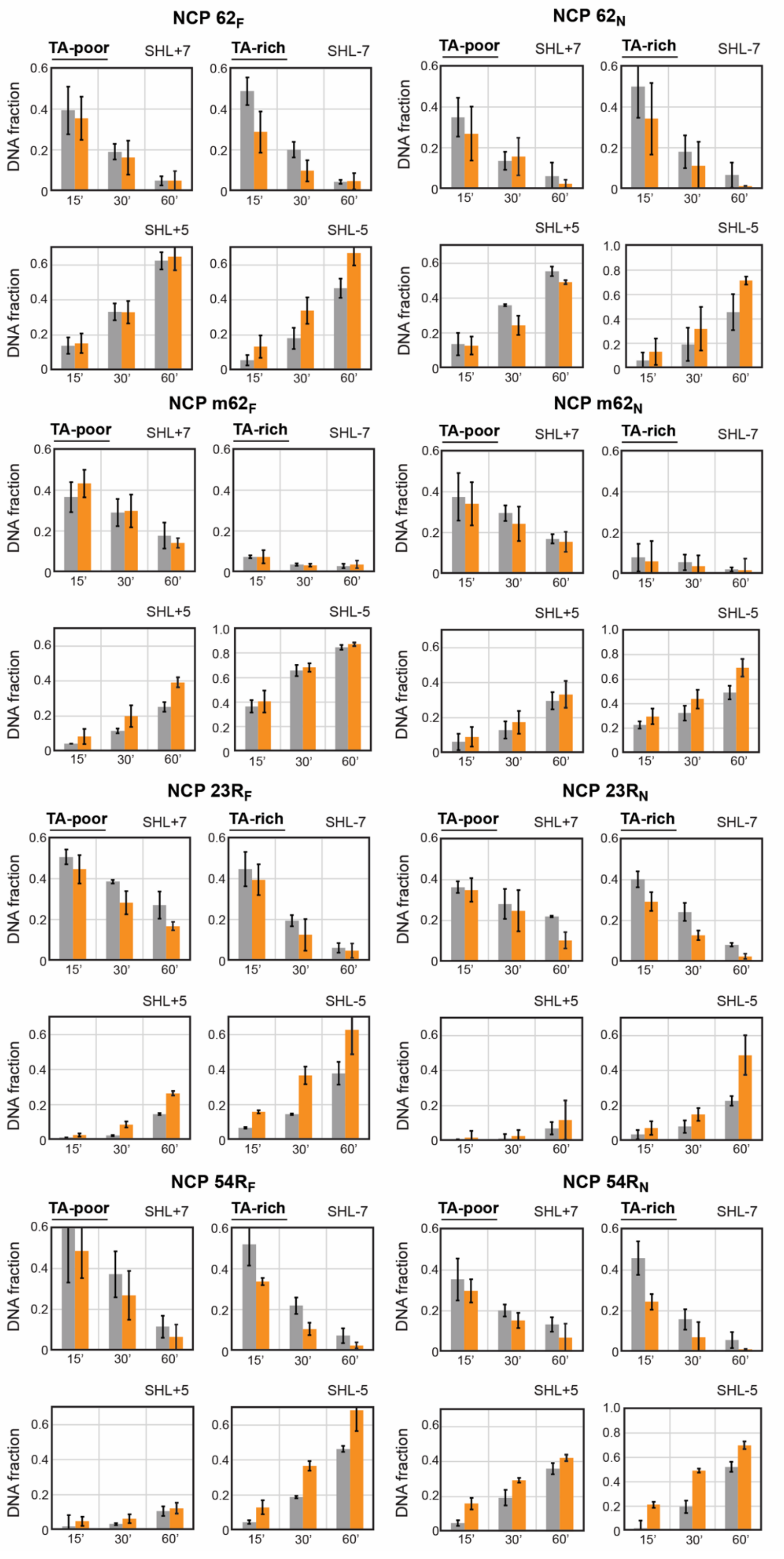


**Figure S3.** DNA fraction (mean ± s.d.) as a function of time (15, 30, and 60 min) at SHL±7 and SHL±5 obtained from ExoIII digestion of 601* NCPs containing H2A (gray) or H2A.Z (orange). More extensive digestion at SHL5 is generally observed with H2A.Z and on the TA-rich side, particularly when the Sox2-Oct4 motifs are inserted on that side (NCP $m62_F$ and $m62_N$).

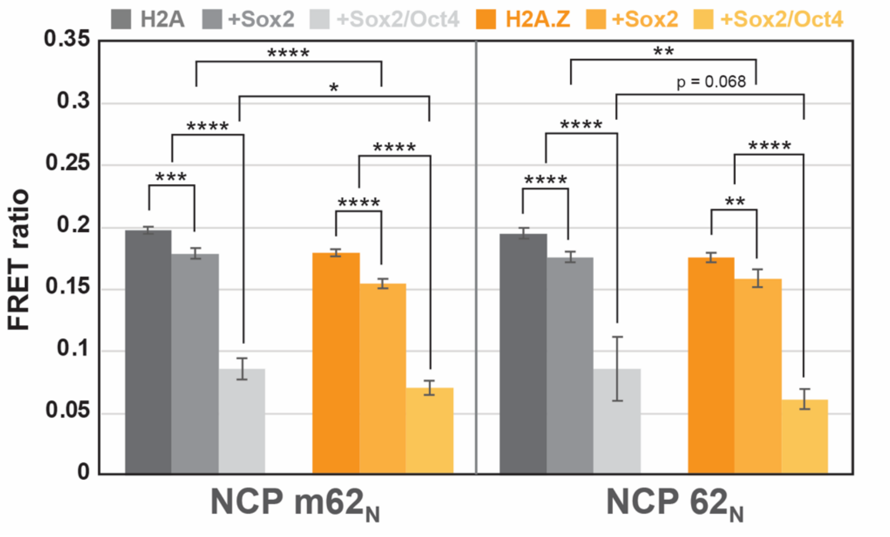


**Figure S4.** Effect of Sox2 and Oct4 binding on DNA unwrapping in nucleosomes. Changes in FRET ratios for H2A or H2A.Z NCPs (20 nM) containing the mNanog motif at SHL-6 ($m62_N$) or SHL+6 ($62_N$) upon addition of Sox2 (10X) or Sox2/Oct4 (10X/10X). Sox2 alone slightly decreases the FRET ratio, consistent with possibly dynamic DNA bending. Sox2+Oct4 lead to larger decreases in the FRET ratio, consistent with Oct4 further stabilizing the highly bent DNA conformation, as in the Cryo-EM structure (PDB ID: 6T90). Statistical significance is indicated by *p*-values obtained from a two-tailed t-test (****$p$<0.0001;***$p$< 0.001;**$p$<0.01; *$p$<0.05).

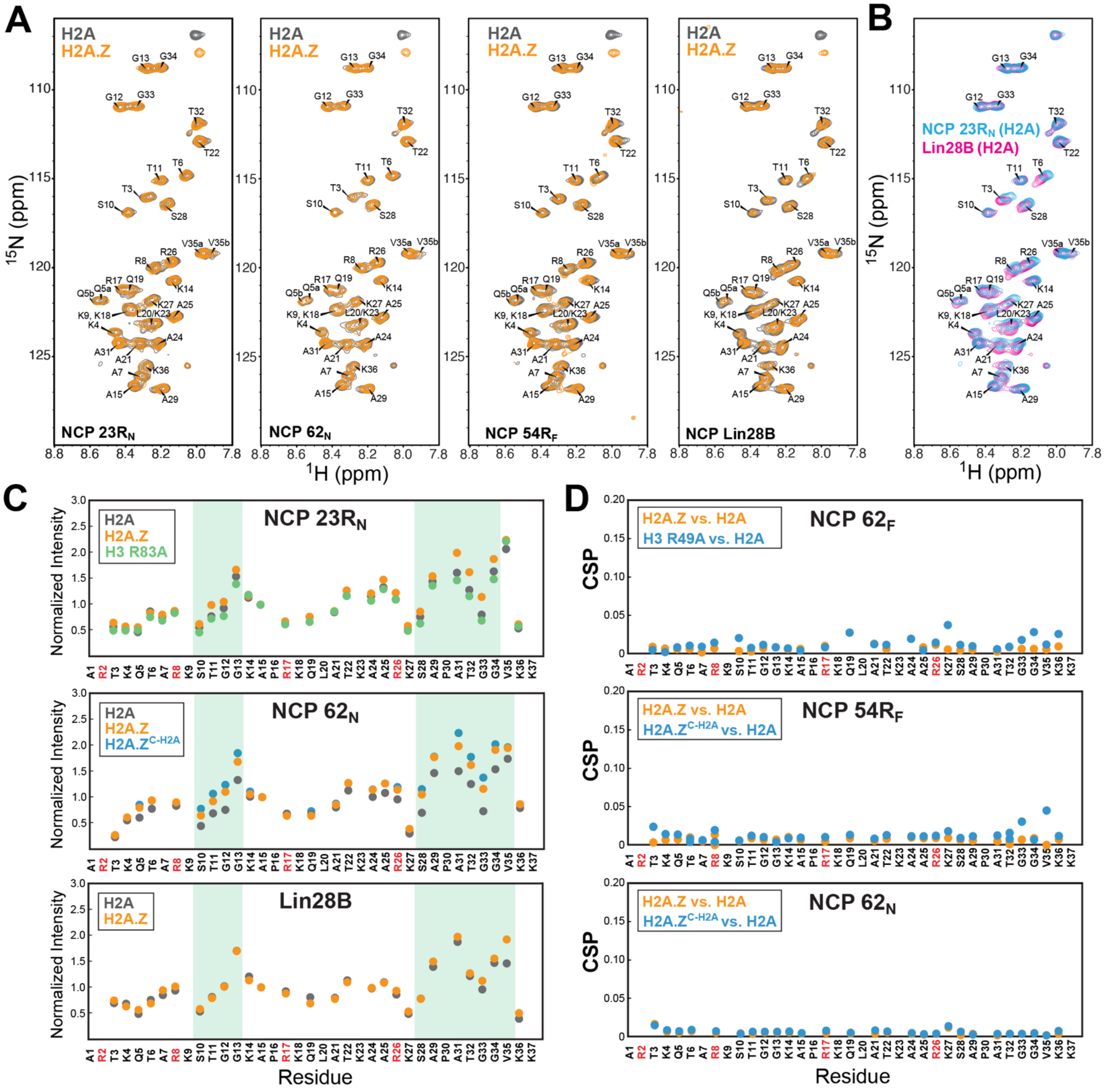


**Figure S5.** H2A.Z perturbs the dynamics of the H3 N-terminal tail. (A) Overlay of $^1$H-$^{15}$N HSQC spectra of $^{15}$N-H3 labeled NCP 23R$_N$, 62$_N$, 54R$_F$, and Lin28B containing H2A or H2A.Z variant. (B) Overlay of $^1$H-$^{15}$N HSQC spectra of NCP 23R$_N$ and Lin28B containing H2A, showing changes in chemical shifts in multiple peaks in Lin28B in the direction of a histone H3 peptide.[7] (C) Normalized peak intensities obtained from spectra in (A) (except for NCP 52R$_F$ included in Figure 3) for NCPs containing H2A, H2A.Z, H3 R83A and H2A.Z$^{C-H2A}$ mutants. Plots depict generally larger changes in peak intensity for flexible hinge regions (highlighted in green) in H2A.Z and mutant histones. (D) Chemical shift perturbations (CSPs) for select NCPs from Figure 3 and panel (A) showing overall small changes with H2A.Z, H2A.Z$^{C-H2A}$, or H3 R49A mutants relative to H2A NCPs.

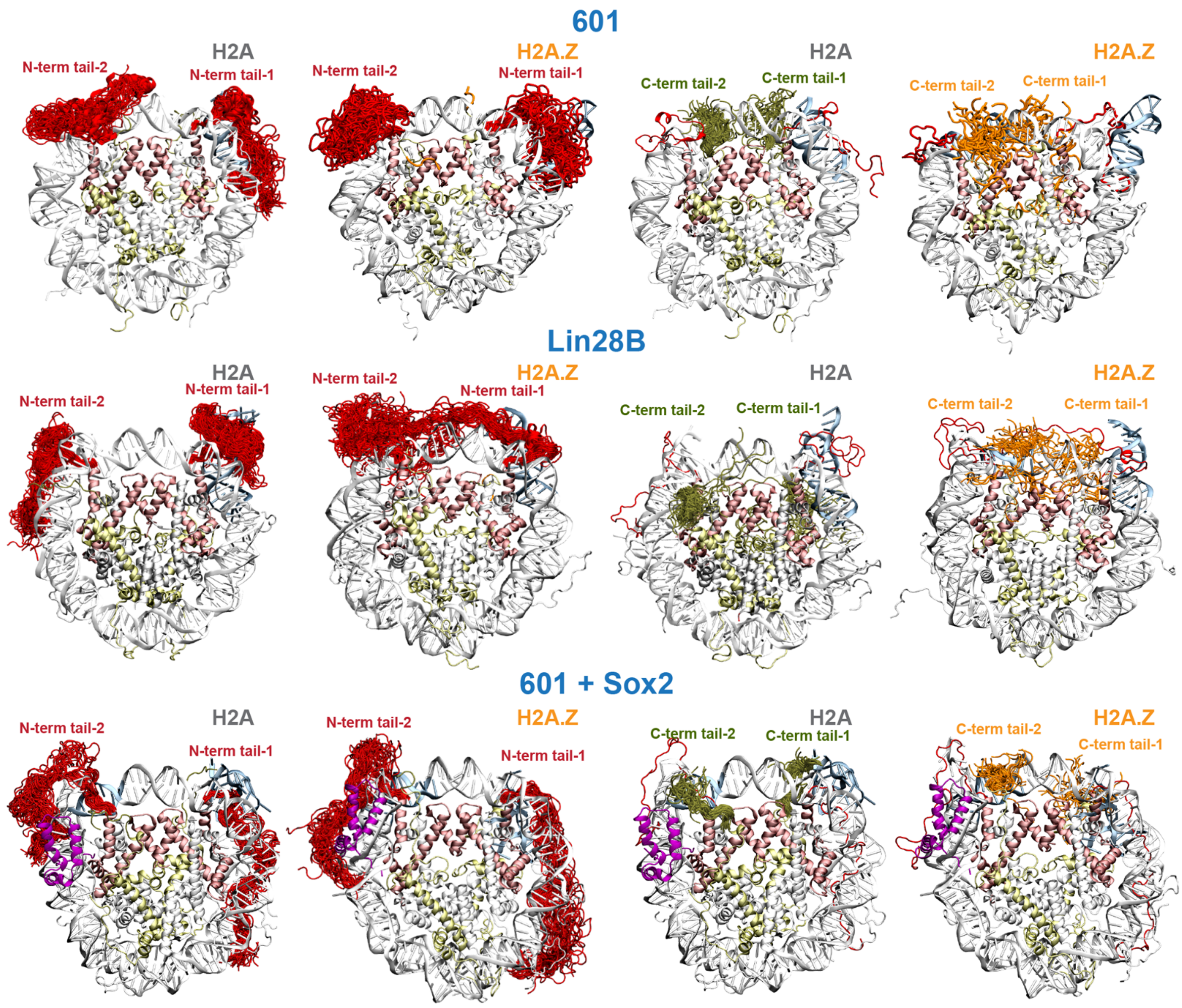


**Figure S6.** Histone tail conformations in 601, Lin28B, and Sox2-bound 601 nucleosome systems. Shown are the conformations of the H3 N-terminal tails (red), H2A C-terminal tails (olive) and H2A.Z C-terminal tails (orange) in 601 (top), Lin28B (center), and 601-Sox2 complex (bottom) nucleosomes. H3 (in H2A.Z) and H2A.Z tails tend to exhibit larger fluctuations than in corresponding H2A systems. The H2A C-tail conformations are shifted in the presence of Sox2 relative to the free 601 NCP.

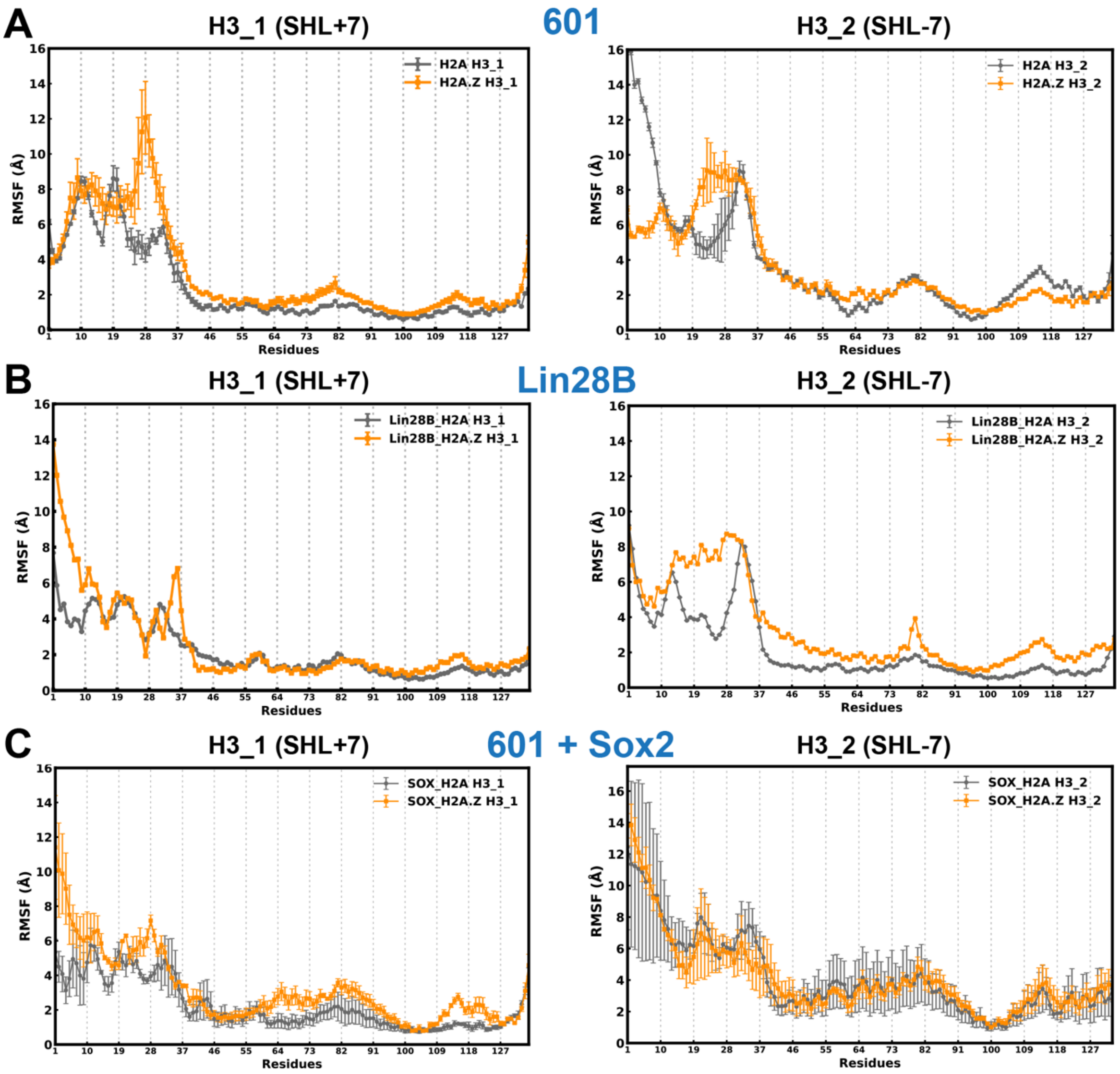


**Figure S7.** Increased histone H3 mobility in the presence of H2A.Z. Root-mean-square fluctuations (RMSF) for Cα atoms of histone H3 (mean ± s.d. for 3 replicas) in (A) 601, (B) Lin28B, and (C) Sox2-bound 601 in H2A (grey) and H2A.Z (orange) systems. Overall greater fluctuations are observed with H2A.Z, particularly in the H3 N-tail. H3_1 and H3_2 designate the histone whose N-terminal tail is located near SHL+7 and SHL-7, respectively.

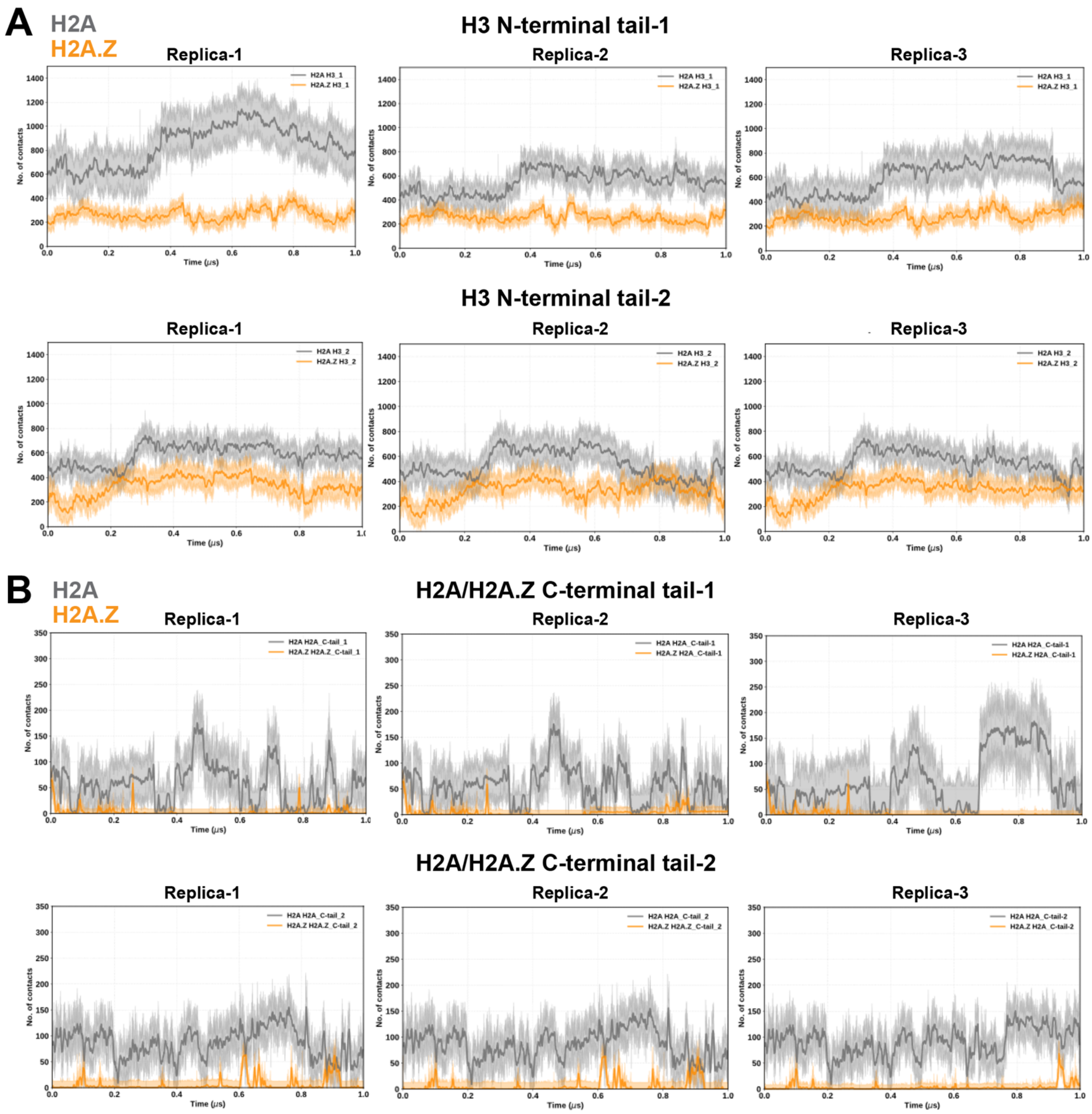


**Figure S8.** H2A.Z reduces histone tail-DNA contacts in 601 nucleosomes. Total number of contacts (4.5 Å cutoff) between DNA and (A) H3 N-terminal tails and (B) H2A/H2A.Z C-terminal tails as a function of time in H2A (grey) and H2A.Z (orange) NCP constructs. The positions of Tail-1 and Tail-2 are near SHL+7 and SHL-7, respectively.

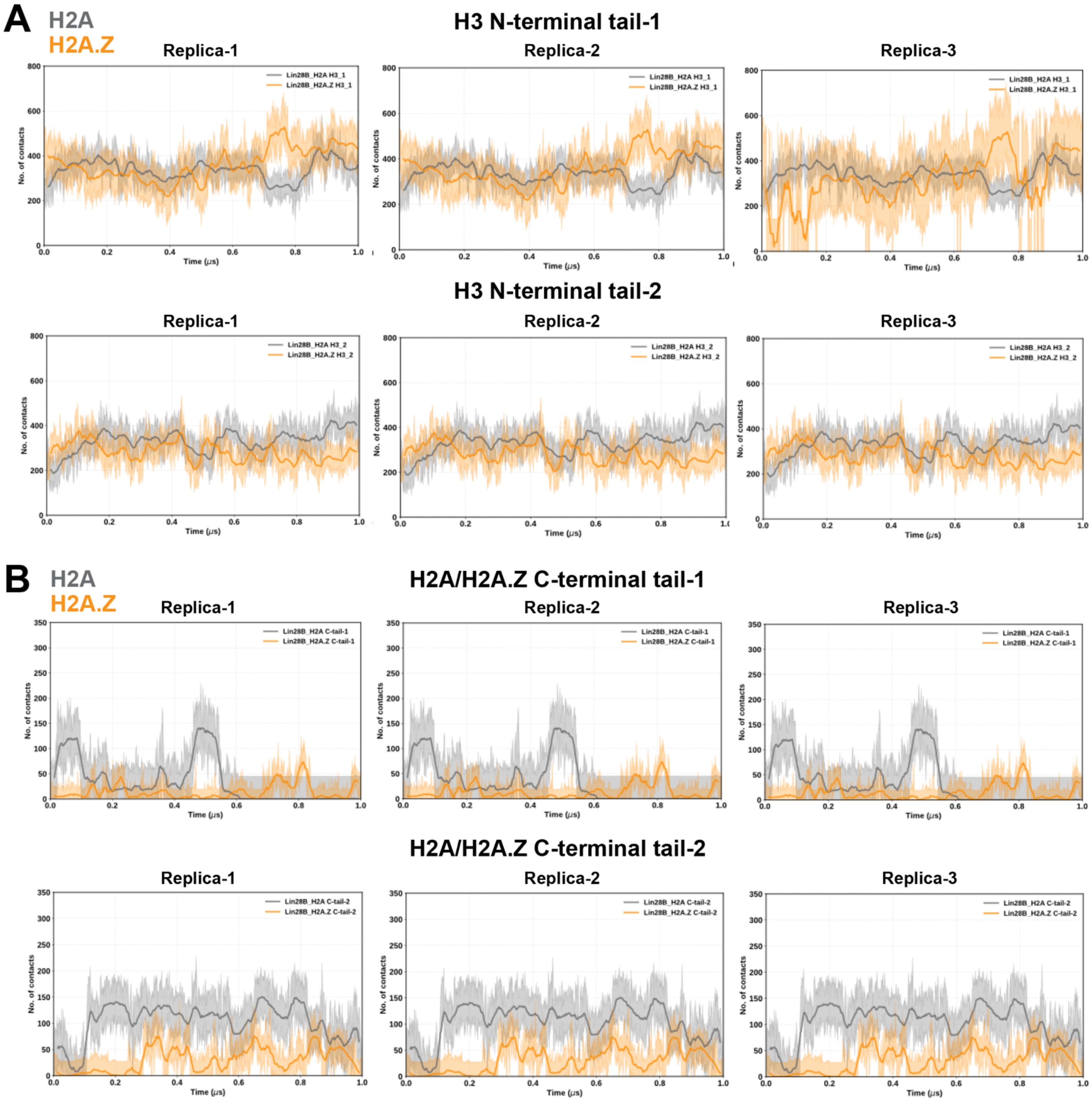


**Figure S9.** H2A.Z reduces histone tail-DNA contacts in Lin28B nucleosomes. Total number of contacts (4.5 Å cutoff) between DNA and (A) H3 N-terminal tails and (B) H2A/H2A.Z C-terminal tails as a function of time in H2A (grey) and H2A.Z (orange) NCP constructs. The positions of Tail-1 and Tail-2 are near SHL+7 and SHL-7, respectively.

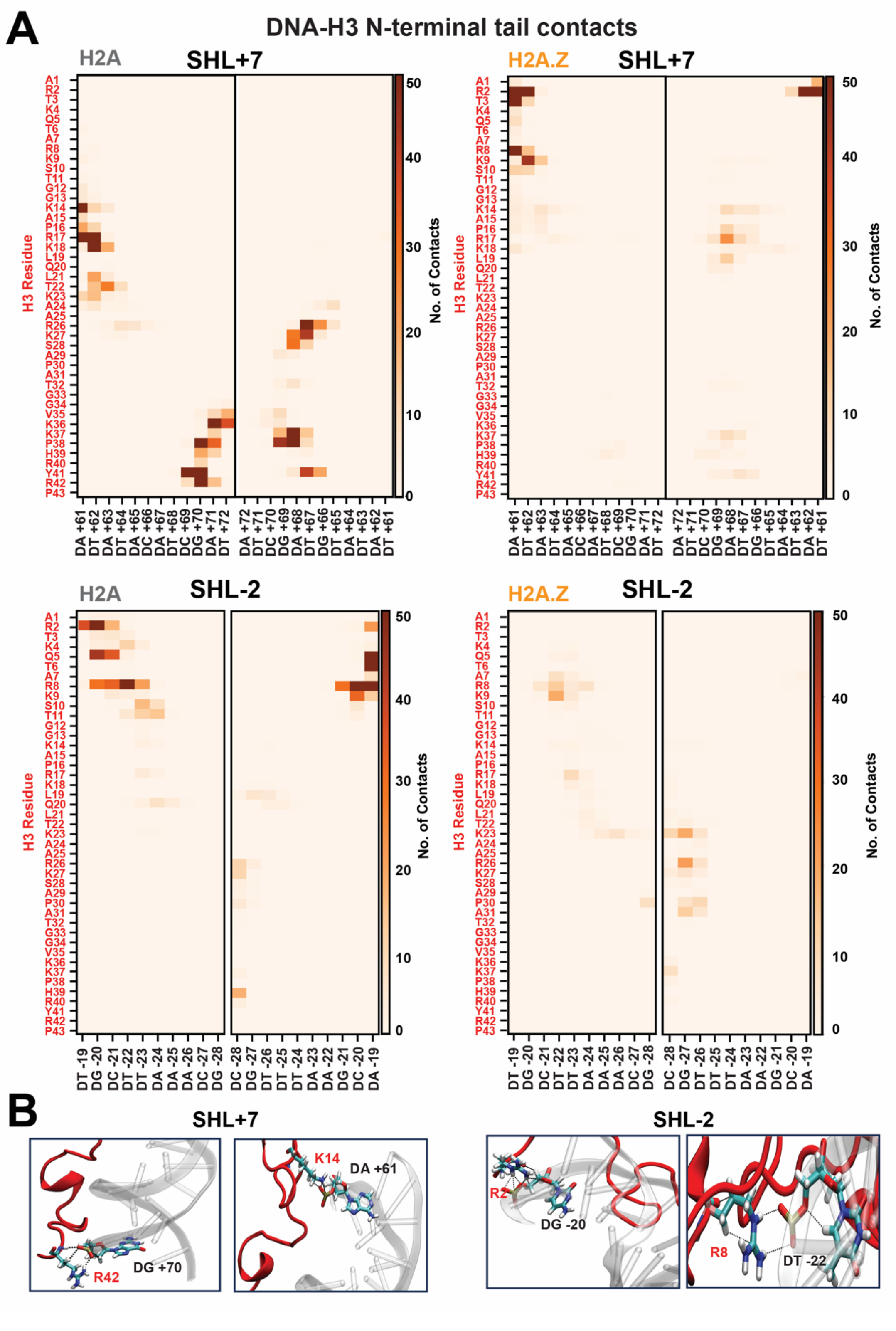


**Figure S10.** H2A.Z reduces H3 tail-DNA contacts at terminal and internal SHLs. (A) A map of the H3 N-tail (residues 1-43) contacts with DNA base pairs at SHL+7 and SHL-2, obtained from 601 NCP simulations, showing a decrease in contacts with H2A.Z. (B) Examples of hydrogen bonds between H3 N-tail residues and DNA at SHL+7 and SHL-2.

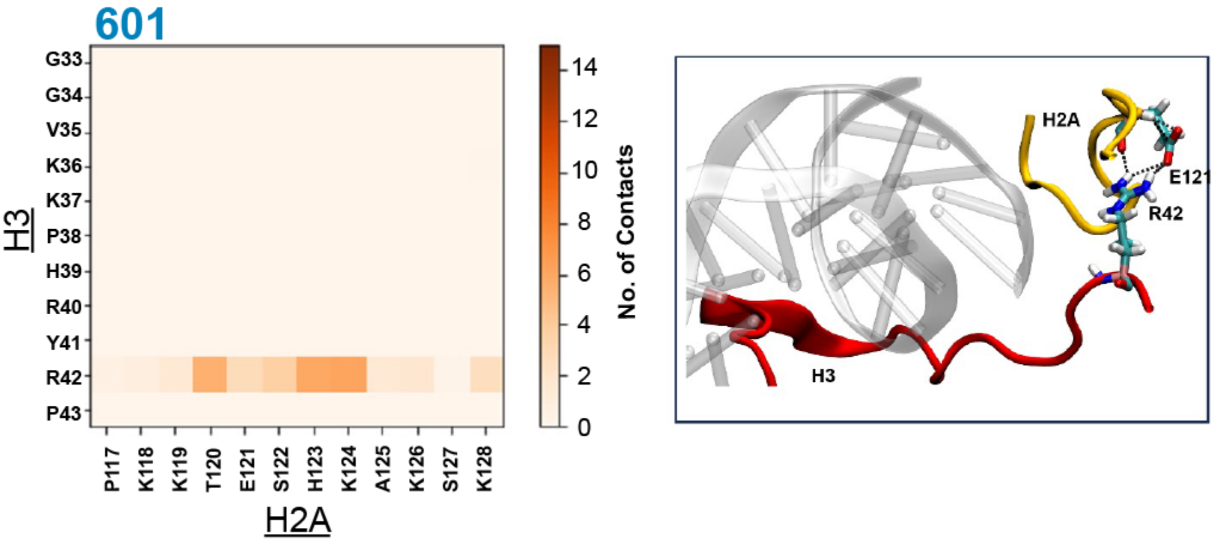


**Figure S11.** The H3 N-terminal tail interacts extensively with the H2A C-terminal tail. (A) Contact map (left) showing interactions between residues in the H3 N-tail (Tail-1) and H2A C-tail (Tail-1) in 601 nucleosomes. H3 R42 interacts with multiple residues in the H2A C-tail, particularly T120 to K124. The snapshot of the H3 and H2A (right) shows formation of hydrogen bonds between H3 R42 and H2A E121.

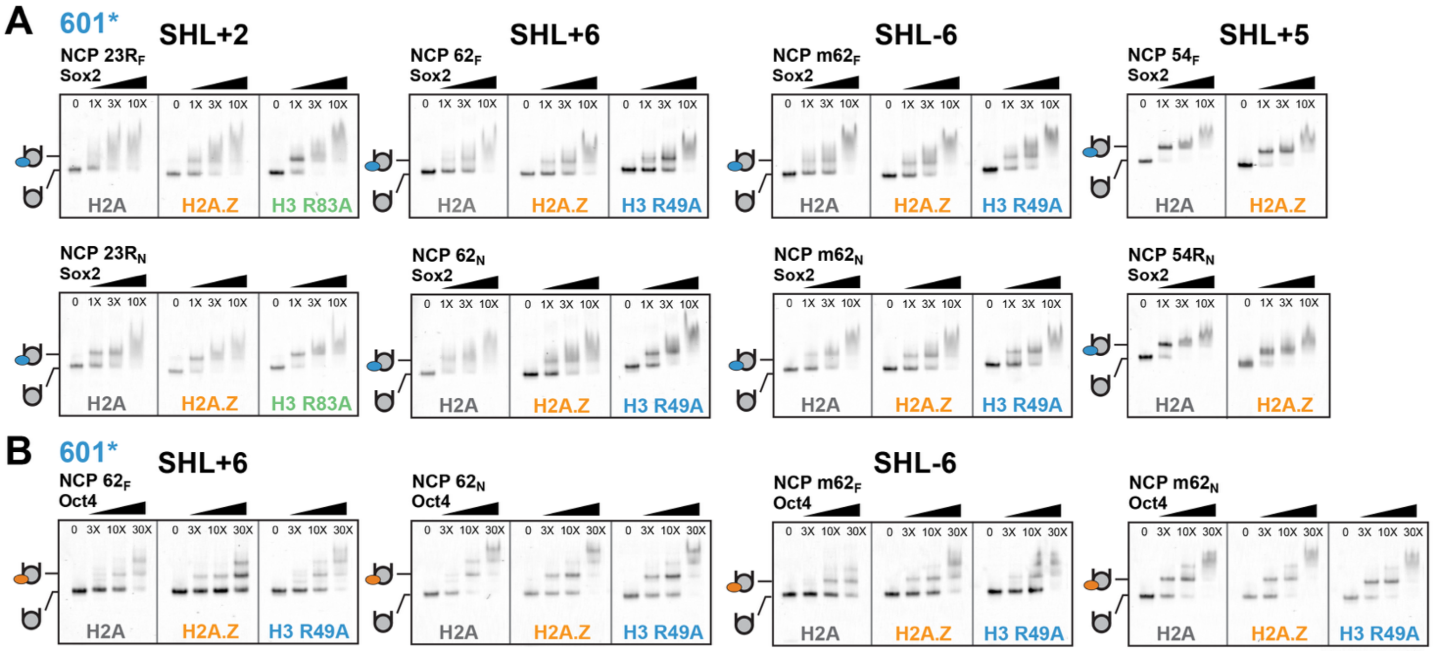


**Figure S12.** H2A.Z stabilizes Sox2 and Oct4 binding at end-positioned and internal nucleosome sites. (A) EMSA analysis of nucleosomes (10 nM) with increasing Sox2 concentrations (1X, 3X, and 10X). Sox2 binding is enhanced by H2A.Z at SHL+2 and SHL±6 and further improved in H3 R83A and H3 R49A mutant nucleosomes, which serve as controls for locally increased DNA mobility at SHL2 and SHL6, respectively. (B) EMSA analysis of nucleosomes (10 nM) in the presence of increasing Oct4 concentrations (3X, 10X, and 30X). H2A.Z promotes formation of a more stable Oct4-NCP complex at SHL+6 (NCP $62_N$ and $62_F$), where either the $POU_{HD}$ or $POU_S$ binding site is solvent exposed. The effect of H2A.Z is less pronounced at SHL-6 (TA-rich side), where DNA is found to be more unwrapped (see Figure S1-S3, Table S3).

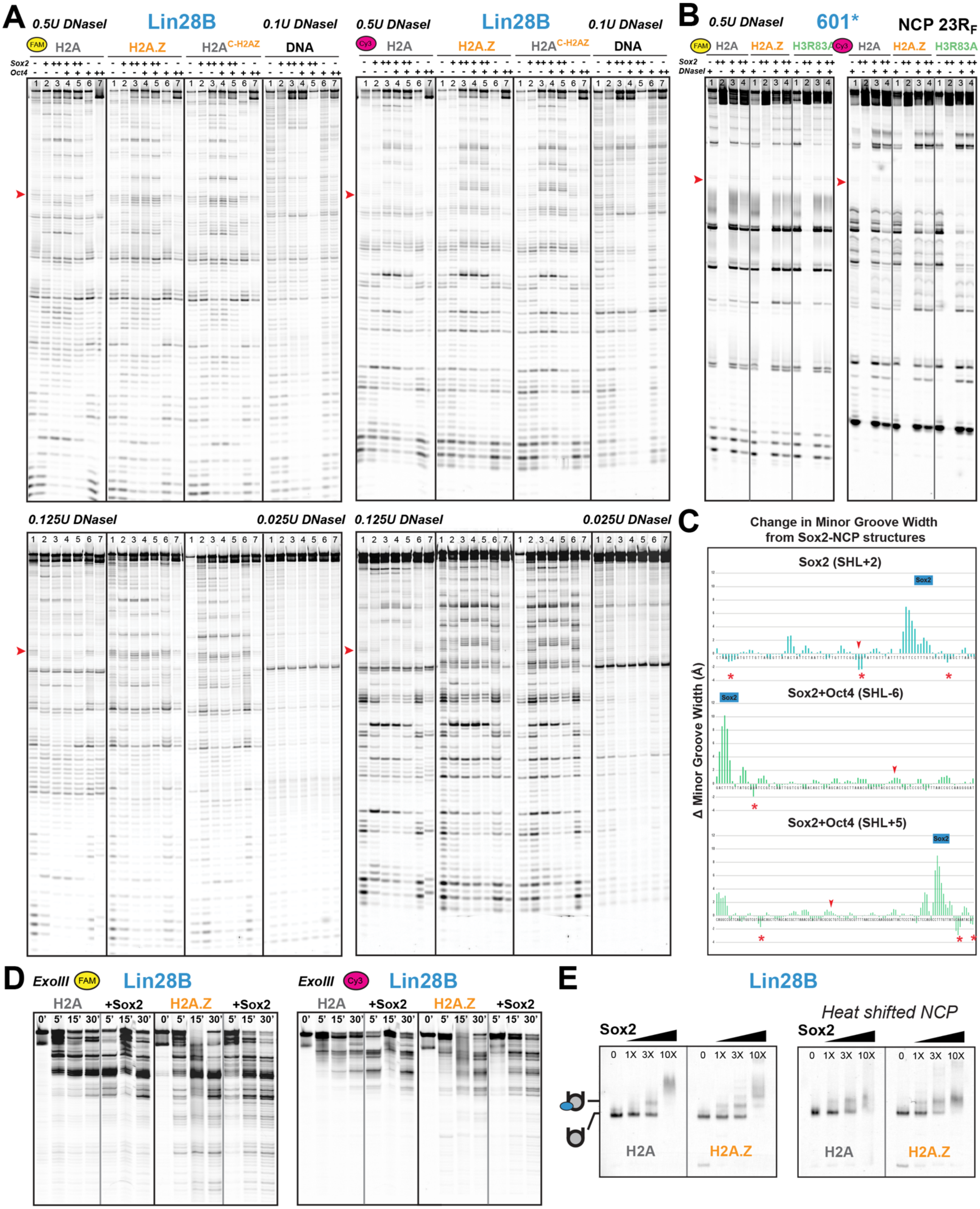


**Figure S13.** Sox2 binding alters the conformation of Lin28B nucleosomes, modulated by H2A.Z. (A) DNaseI digestion, at two enzyme concentrations and monitored by denaturing gel electrophoresis, of Lin28B NCPs containing H2A, H2A.Z, H2A-C$^{H2A.Z}$ chimera or Lin28B DNA in the absence (lane 1) or presence of Sox2 (1X(+), lane 2; 10X(++), lane 3), Sox2/Oct4 (1X/3X(+/+), lane 4; 10X/3X(++/+), lane 5), or only Oct4 (3X(+), lane 6; 30X(++), lane 7). Sox2 yields multiple footprints and hypersensitive sites on nucleosomes, further enhanced by Oct4 at low concentrations (compare lane 2 and lane 4). In contrast, binding of Oct4 alone to NCP or Sox2 to naked DNA results in localized footprints. The presumed nucleosome dyad is marked with a red arrow, consistent with the increased DNaseI signal at exposed minor groove sites. (B) DNaseI digestion of 601* 23$R_F$ NCPs containing H2A, H2A.Z, or H3 R83A in the absence or presence of Sox2 (3X(+), 10X(++)), showing the absence of hypersensitive sites with Sox2 and no visible changes in dyad digestion with H2A.Z, in contrast to Lin28B NCPs in (A). (C) Changes in minor groove width of NCP complexes with Sox2 bound at SHL+2 (PDB ID: 6T7B), SHL-6 (PDB ID: 6T90), or SHL+5 (PDB ID: 6Y0V) relative to the corresponding free NCP structures (PDB ID: 6T79 and 6T93), performed using the X3DNA. Larger minor groove narrowing is indicated by stars (*) and the dyad positions are indicated by red arrows. (D) Exonuclease III digestion time courses for canonical H2A and H2A.Z Lin28B nucleosomes in the absence or presence of Sox2 (10X), showing increased susceptibility of H2A NCPs with Sox2 and smaller effect for H2A.Z NCPs. (D) EMSA gels of Sox2 binding to Lin28B NCPs showing similar binding to H2A and H2A.Z constructs and little effect of heat shifting on Sox2 binding. (E) The heat-shifted NCPs (incubated at 37 $^{o}$C for 3 hrs to remove differently positioned, kinetically trapped species) still contained the slower-migrating impurity observed in non-heat-shifted NCPs.

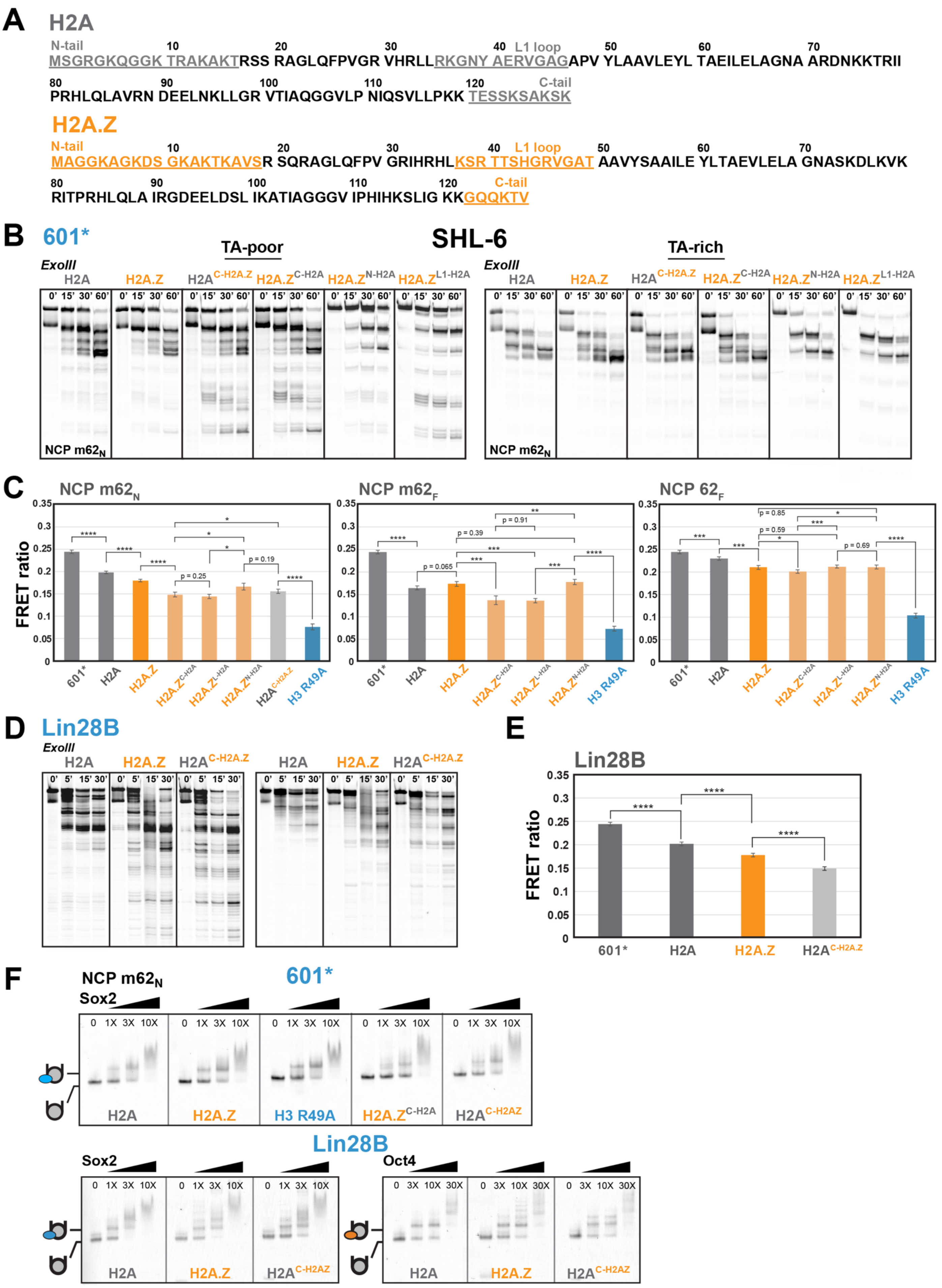
A
H2A
N-tail
MSGRGKQGGK TRAKAKTRSS RAGLQFPVGR VHRLLRKGNY AERVGAGAPV YLAAVLEYL TAEILELAGNA ARDNKKTRII
L1 loop
PRHLQLAVRN DEELNKLLGR VTIAQGGVLP NIQSVLLPKK TESSKSAKSK
C-tail
H2A.Z
N-tail
MAGGKAGKDS GKAKTKAVSR SQRAGLQFPV GRIHRHLKSR TTSHGRVGAT AAVYSAAILE YLTAEVLELA GNASKDLKVK
L1 loop
RITPRHLQLA IRGDEELDSL IKATIAGGGV IPHIHKSLIG KKGQQKTV
C-tail
B
601*
TA-poor
SHL-6
TA-rich
ExoIII
H2A
H2A.Z
H2A.Z N-H2A
H2A.Z L1-H2A
NCP m62N
C
NCP m62N
NCP m62F
NCP 62F
FRET ratio
601*
H2A
H2A.Z
H2A.Z C-H2A
H2A.Z L-H2A
H2A.Z N-H2A
H2A C-H2A.Z
H3 R49A
D
Lin28B
ExoIII
E
Lin28B
FRET ratio
F
NCP m62N
601*
Sox2
H3 R49A
Lin28B
Oct4

**Figure S14.** The H2A.Z C-terminal tail and L1 loop modulate nucleosome dynamics. (A) Amino acid sequences of H2A (grey) and H2A.Z (orange) showing the N-terminal tail, L1 loop, and C-terminal tail regions (highlighted and underlined) swapped in mutation constructs. (B) ExoIII digestion time courses (0, 15, 30, 60 min) for H2A, H2A.Z, H2A mutant containing an H2A.Z C-tail ($H2A^{C\text{-}H2A.Z}$), and H2A.Z mutants carrying the H2A N-tail ($H2A.Z^{N\text{-}H2A}$), L1 loop ($H2A.Z^{L1\text{-}H2A}$), or C-tail ($H2A.Z^{C\text{-}H2A}$). C-tail and L1 loop mutants show significant internal digestion on the TA-poor side that is suppressed with H2A.Z. (C) Corresponding FRET ratios for NCP $m62_N$ constructs as in (B), showing decreased FRET and increased DNA unwrapping for C-tail and L1 loop mutants. Also shown are FRET values for NCP $m62_F$ and $62_F$ constructs. Statistical significance is indicated by *p*-values obtained from a two-tailed t-test (****$p$<0.0001;***$p$< 0.001;**$p$<0.01; *$p$<0.05). (D) ExoIII digestion time courses (0, 5, 15, 30 min) for H2A, H2A.Z, and $H2A^{C\text{-}H2A.Z}$ Lin28B NCPs showing increased digestion with H2A Z and the C-tail mutant. (E) Corresponding FRET ratio of Lin28B NCPs showing decreased values of H2A.Z or C-tail swap mutant relative to canonical NCPs (*p*-values are indicated as in (C)). (F) Corresponding EMSA gels of $m62_N$ and Lin28B NCPs showing small effects of C-tail mutants on Sox2 or Oct4 binding to nucleosomes.

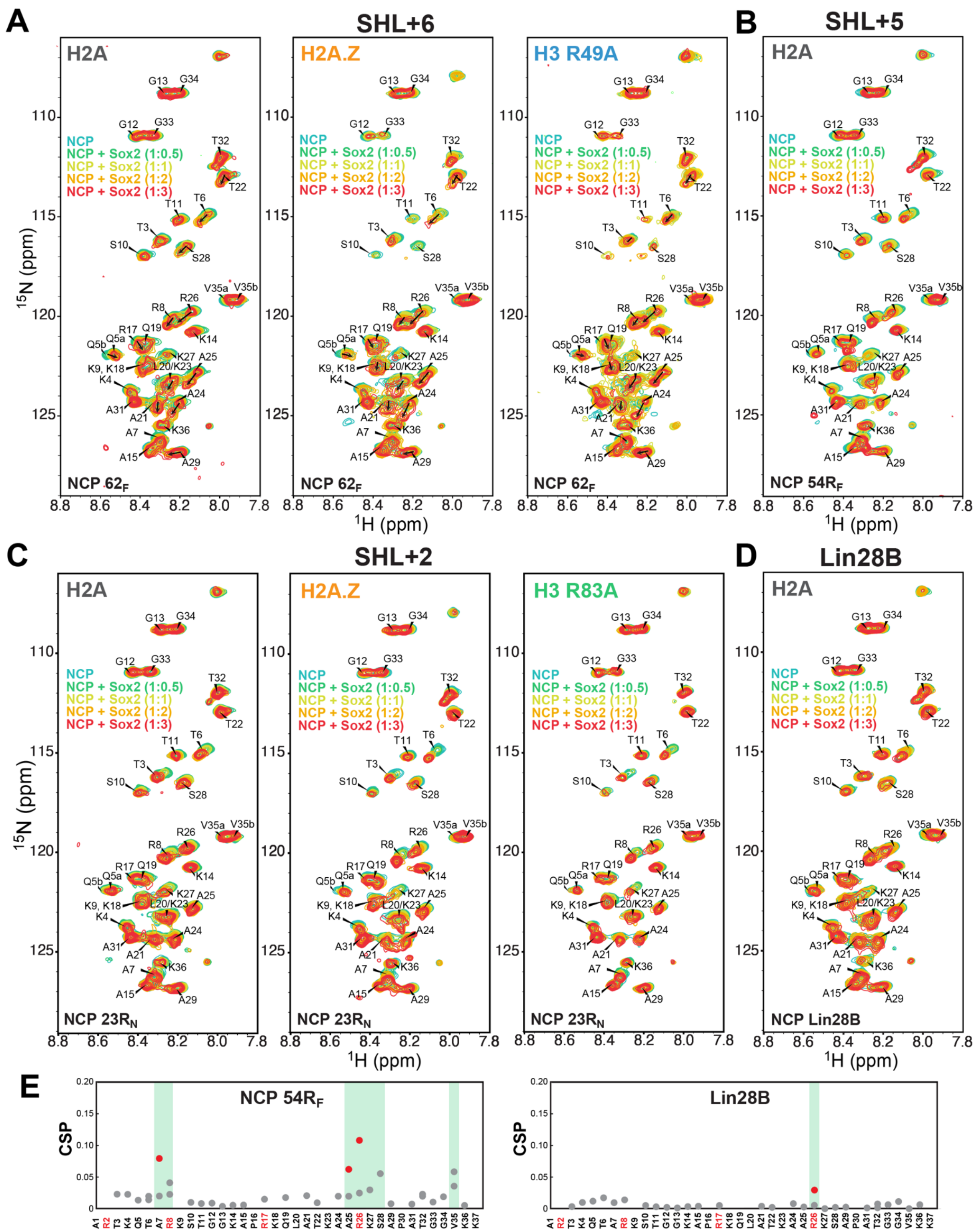


**Figure S15.** Sox2 perturbs the H3 N-terminal tail when binding at different nucleosome positions. (A) Overlay of $^{1}$H-$^{15}$N HSQC spectra of $^{15}$N-H3 NCP 62R$_F$ (20 μM) containing H2A (left), H2A.Z (center) or H3 R49A (right) with increasing concentrations of Sox2 (0.5X, 1X, 2X and 3X), collected at 37$^{o}$C. A second population of peaks appears with the addition of Sox2 and is consistent with the release of the H3 N-tail from DNA. Similar overlay of $^{1}$H-$^{15}$N HSQC spectra of (B) $^{15}$N-H3 NCP 54R$_F$, (C) $^{15}$N-H3 NCP 23R$_N$ containing H2A, H2A.Z or H3 R83A, and (D)$^{15}$N-H3 NCP Lin28B with Sox2. (E) Chemical shift perturbations (CSPs) for NCP 54R$_F$ and Lin28B at 1:2 Sox2:NCP ratio showing that the H3 N-tail is perturbed to a much smaller extent when Sox2 binds to internal sites at SHL+5 (and SHL+2 in Figure 7B) of 601* than at SHL6 or at putative sites on Lin28B (larger CSPs highlighted in green).

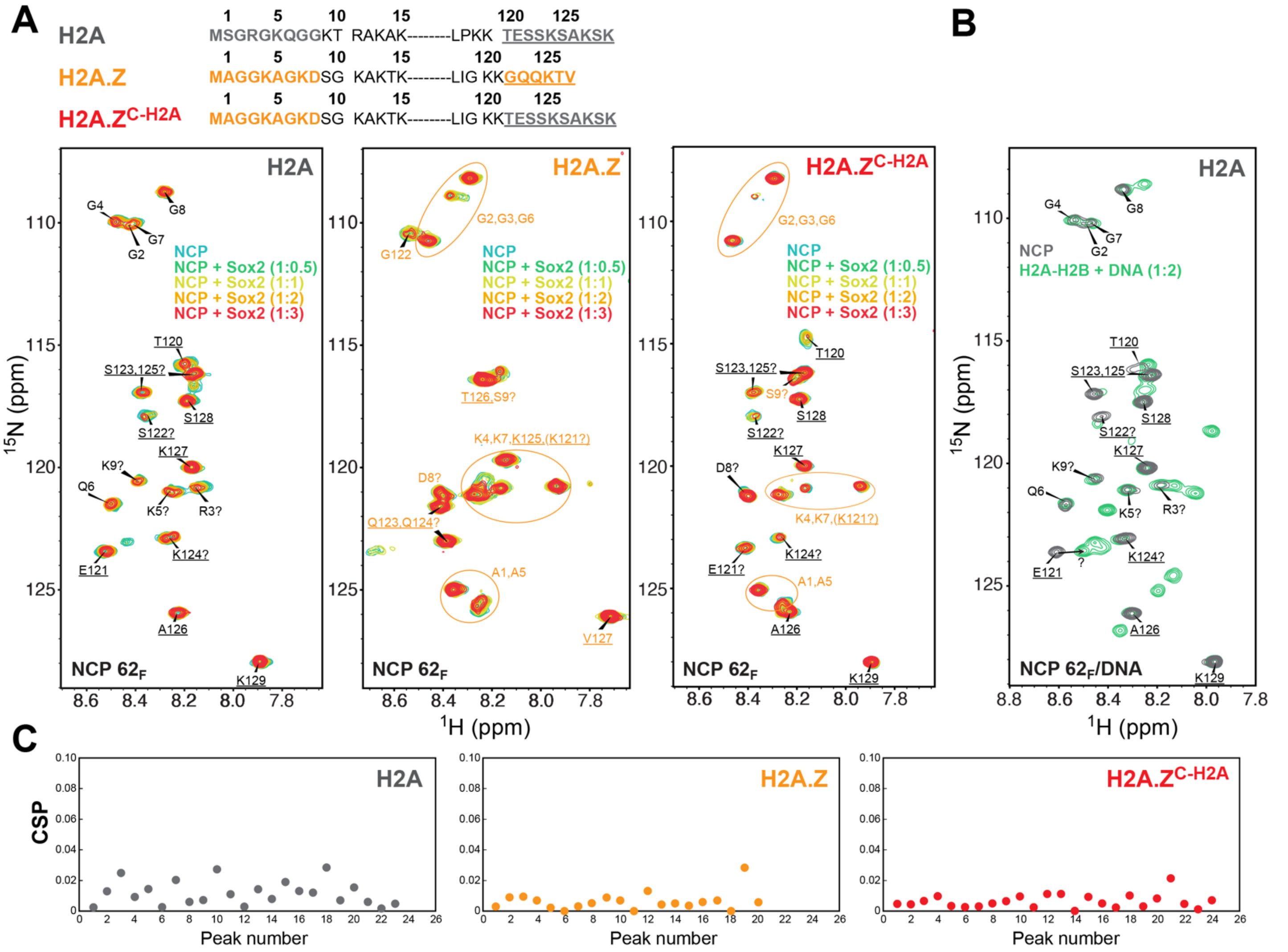


**Figure S16.** Sox2 perturbs the H2A C-terminal tail more than for H2A.Z. (A) $^{1}$H-$^{15}$N HSQC spectral overlay of NCP $62_F$ containing $^{13}$C/$^{15}$N-labeled H2A (grey), H2A.Z (orange), or H2A.Z$^{C-H2A}$ (red) with increasing concentrations of Sox2 (0.5X, 1X, 2X and 3X), collected at 37°C. Amino acid sequences showing visible N-tail and C-tail residues, with the C-tail underlined (note that H2A.Z$^{C-H2A}$ numbering matches that of H2A in the spectra). (B) $^{1}$H-$^{15}$N HSQC spectral overlay of NCP $62_F$ containing $^{15}$N-H2A (grey) and an ($^{15}$N-H2A)-H2B dimer (20 μM) bound to a 40 base pair DNA duplex (green), collected at 25°C. E121 is significantly shifted or broadened in the histone dimer-DNA complex relative to NCP $62_F$, indicating a change in E121 conformation, protonation of interactions. The predicted E121 peak position (indicated with "?") with DNA matches that with the predicted position in the H2A.Z$^{C-H2A}$ mutant, where the tail conformation is altered. (C) Chemical shift perturbations (CSPs) in NCP $62_F$ containing H2A, H2A.Z, or H2A.Z$^{C-H2A}$ with the addition of 3X Sox2, showing overall larger spectral changes in canonical versus H2A.Z nucleosomes (peaks are arranged in increasing $^{15}$N chemical shift).

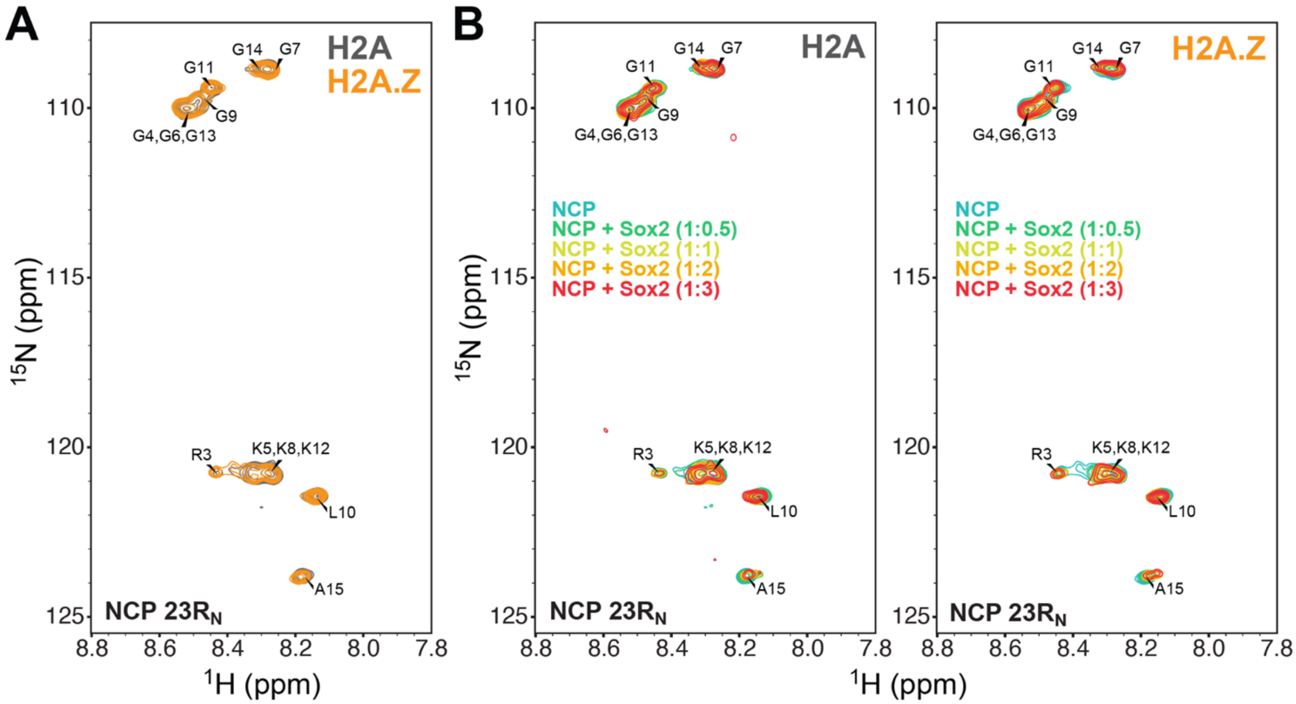


**Figure S17.** Sox2 binding at SHL2 has limited effect on the H4 N-terminal tail. (A) Overlay of $^{1}H$-$^{15}N$ HSQC spectra of $^{15}N$-H4 NCP 23R$_N$ (~20 μM) containing H2A (grey) or H2A.Z (orange). (B) Overlay of $^{1}H$-$^{15}N$ HSQC spectra of $^{15}N$-H4 NCP 23R$_N$ (~20 μM) with increasing concentrations of Sox2 (0.5X, 1X, 2X, and 3X), showing small chemical shift changes with either H2A (left) or H2A.Z (right).

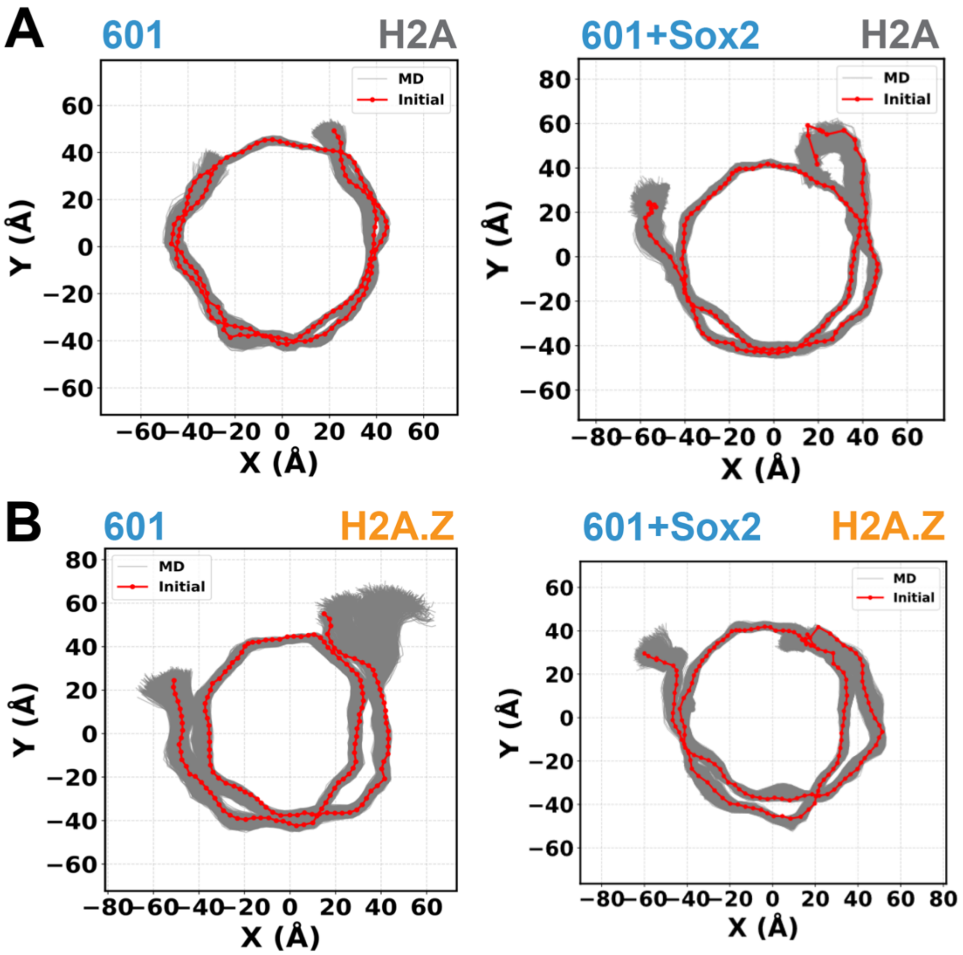


**Figure S18.** Sox2 binding globally perturbs DNA in canonical and H2A.Z 601 nucleosomes. DNA backbone traces for 601 NCP containing (A) H2A and (B) H2A.Z in the presence or absence of Sox2, showing significant backbone distortion, gyre misalignment, and measurable vertical compression, particularly for the H2A.Z NCP.

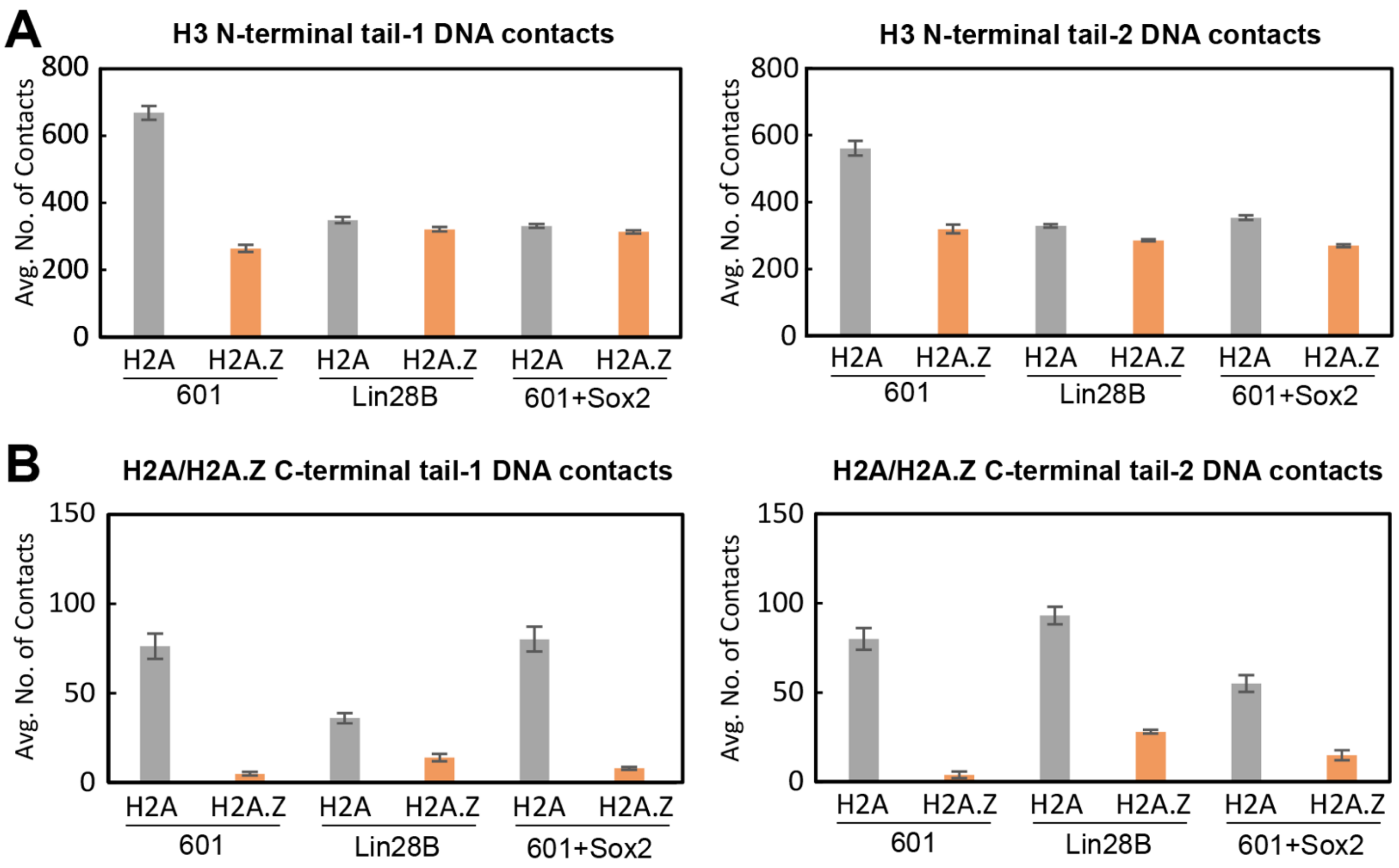


**Figure S19.** Sox2 binding disrupts H3 N-tail and H2A C-tail interactions with DNA. (A) Total DNA contacts for the H3 N-terminal tail-1 (SHL+7) and tail-2 (SHL-7) in 601, Lin28B and Sox2-bound 601 NCPs, showing (i) larger reduction for H2A versus H2A.Z 601 NCP with Sox2 binding (near tail-2, SHL-6) and (ii) larger reduction for unbound 601 versus Lin28B NCPs with H2A.Z. (B) Total DNA contacts for the H2A/H2A.Z C-terminal tail-1 (SHL+7) and tail-2 (SHL-7) in 601, Lin28B and Sox2-bound 601 NCPs, showing a large decrease for H2A 601 NCP but a small increase for H2A.Z 601 NCP with Sox2 binding (near tail-2, SHL-6). The Lin28B NCP exhibits increased H2A.Z C-tail-DNA contacts relative for 601, which could be dependent on the local DNA sequence.

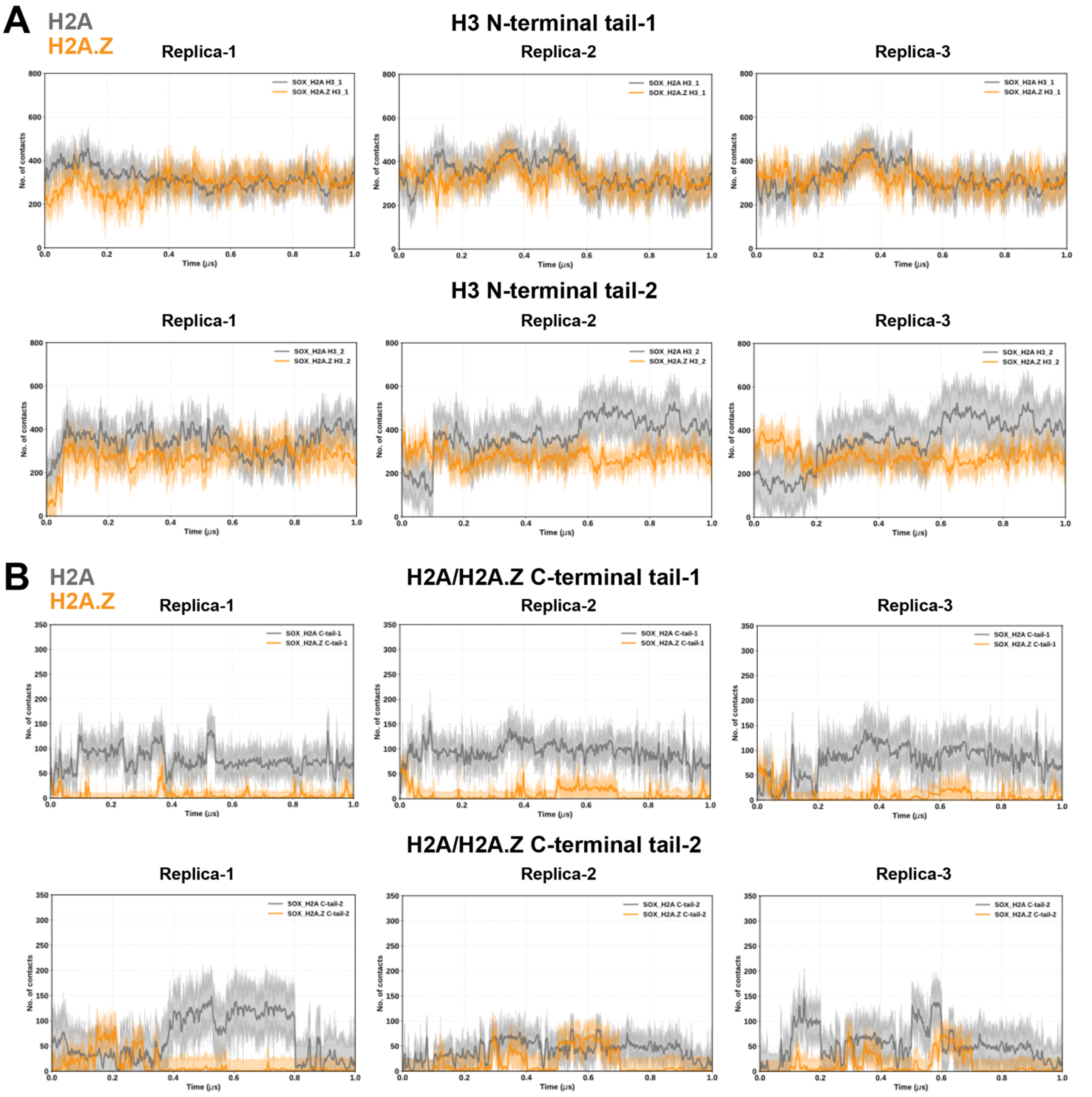


**Figure S20.** Sox2 binding at SHL-6 reduces histone tail-DNA contacts in H2A.Z 601 nucleosomes. Total number of contacts (4.5 Å cutoff) between DNA and (A) H3 N-terminal tails and (B) H2A/H2A.Z C-terminal tails as a function of time in H2A (grey) and H2A.Z (orange) NCP constructs. The positions of Tail-1 and Tail-2 are near SHL+7 and SHL-7, respectively.

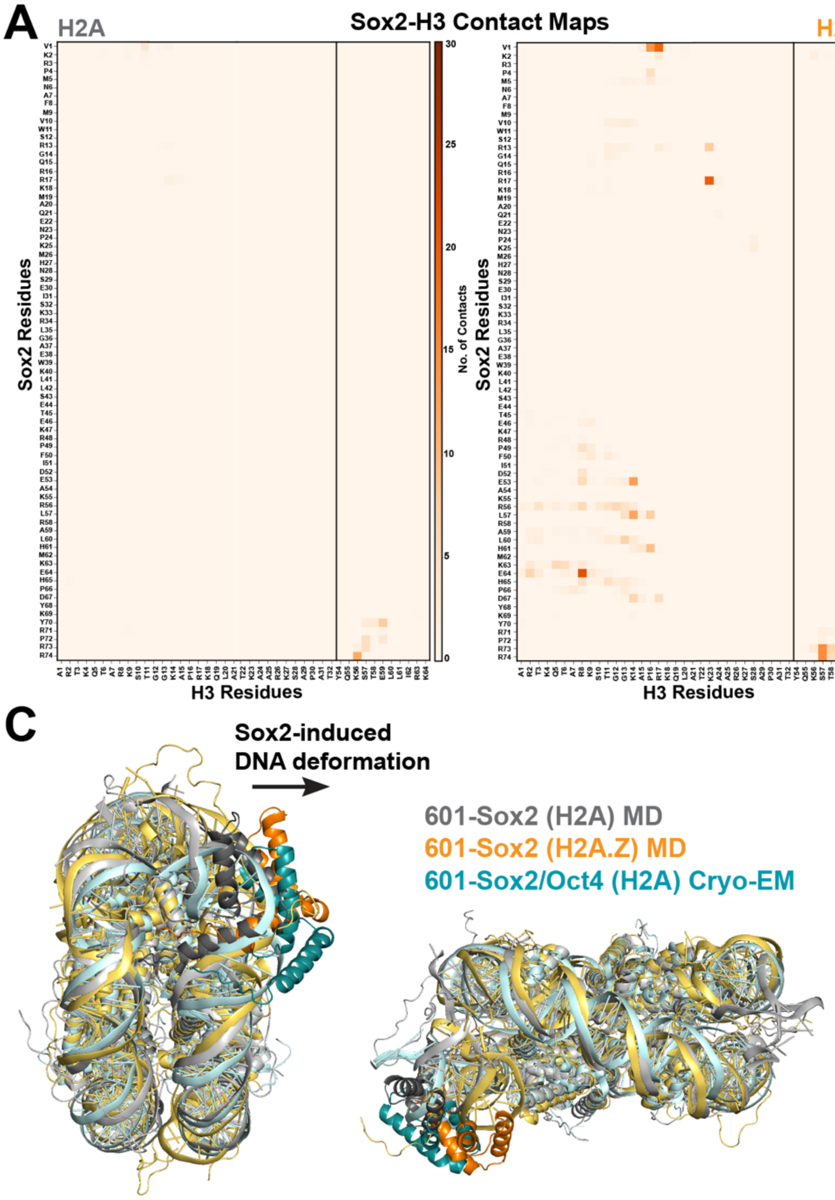


**Figure S21.** Sox2 induces greater DNA displacement and forms unique interactions with histone H3 in H2A.Z nucleosomes. (A) Contact maps of histone H3 (N-tail residues 1-32 and $\alpha$1-helix residues 54-64) and the Sox2 HMG domain from MD simulations of 601-Sox2 H2A and H2A.Z complexes, showing unique interactions in the presence of H2A.Z that are rare or absent in the presence of H2A. (B) Examples of potentially strong hydrogen bonding and electrostatic interactions between Arg/Lys (R8 and K14) residues in the H3 N-tail and Glu (E53 and E64) residues in the Sox2 $\alpha$3-helix. (C) Structural overlay of 601-Sox2/H2A complex (grey), 601-Sox2/H2A.Z complex (orange) from MD simulations and the Cryo-EM 601-Sox2/Oct4 complex (teal), showing that, with H2A.Z, Sox2 pulls and deforms DNA more extensively and more similar to the Sox2-Oct4-NCP ternary complex.
